# LIF is a pro-algesic mediator in human and mouse models of arthritis-related pain

**DOI:** 10.64898/2026.10.12.758476

**Authors:** Rosie Ross, Yuening Li, Julia Vlachaki Walker, Mosab Ali Awadelkareem, Irene Zebochin, George Goodwin, Alina-Cristina Marin, Teodora Trendafilova, Innes Montague, Zoe Rutter-Locher, Sarah Ryan, Friederike Baldeweg, Katja Freytag, Rophi Condor, Natalie Arridge, Christopher D. Buckley, David L. Bennett, Frances Humby, Bruce W. Kirkham, Franziska Denk, Leonie S. Taams

## Abstract

**Purpose:** Emerging clinical and experimental data suggest that JAK inhibitors (JAKi) are effective in reducing pain in Rheumatoid Arthritis (RA). We recently showed that JAK/STAT signalling cytokines, specifically IL-6, LIF and IL-11, are present in the RA joint and can act directly on human induced pluripotent stem cell (iPSC) derived sensory neurons by increasing their STAT3 phosphorylation (pSTAT3) levels and expression of select pain-relevant genes. In this study, we determined that LIF is the most consequential cytokine out of the three, altering neuronal signalling, outgrowth and pain-like behaviours in *in vitro* and *in vivo* models of arthritis.

**Methods:** Human iPSC-derived sensory neurons were stimulated with RA synovial fluid (SF) or with cytokine-treated RA synovial fibroblast conditioned media (CM) in the absence or presence of anti-cytokine antibodies. pSTAT3 and pSTAT1 levels, and gene expression patterns, were assessed by Western blot and RNA-sequencing (RNA-seq). The immortalised dorsal root ganglion (DRG)-derived cell line (HD 10.6) was assessed for LIF-induced JAK/STAT signalling in addition to neurite outgrowth assays as a functional readout of the sensory neurons. Lastly, mouse models were used to assess pain-related behaviours *in vivo*.

**Results:** LIF was found to be consistently present in the SF from painful joints from individuals with inflammatory arthritis. Western blotting revealed that the addition of RA SF to iPSC-derived sensory neurons led to LIF-dependent induction of pSTAT3 and pSTAT1, while RNA-seq demonstrated that LIF alters the transcriptome of sensory neurons more prominently than IL-6 and IL-11, up-regulating signature genes related to peripheral neuron sensitization, such as *TRPV1*, *NTRK1*, *BDNF*, *HRH1* and *SCN10A.* Similarly, cytokine-treated fibroblast conditioned media was shown to induce neuronal pSTAT3 and pSTAT1 and upregulate expression of *BDNF*, *HRH1* and *VGF* in a LIF-dependent manner. Additionally, the immortalised dorsal root ganglion (DRG)-derived cell line (HD 10.6), which also exhibited LIF-induced pSTAT1 and pSTAT3, showed increased neurite outgrowth after axotomy when treated with LIF. Finally, in a mouse model of inflammatory arthritis, LIF was shown to be significantly increased in the affected joint. Furthermore, intraarticular injection of LIF into a healthy mouse knee resulted in increased responsiveness to painful pressure stimuli in that joint.

**Conclusions:** Our data indicate that LIF may be a pro-algesic mediator and a possible therapeutic target to improve pain management in individuals living with inflammatory arthritis.

## Introduction

Rheumatoid Arthritis (RA) is a chronic autoimmune disease characterised by synovial tissue inflammation and joint pain and is often associated with systemic comorbidities. Over the last two decades, there has been tremendous progress in disease management and treatment strategies for people living with RA, through the advent of biologic therapies and the shift towards early intensive therapy in newly diagnosed individuals with the goal to decrease disease activity (Alivernini et al., 2022). Whilst these developments have led to substantial improvements in managing joint inflammation, the effective management of joint pain remains more challenging. For example, a 10-year prospective study on almost 1,700 individuals with RA receiving routine care showed that only some components of the disease activity score 28 (DAS28), such as the erythrocyte sedimentation rate and number of swollen joints decreased significantly over this period. In contrast, measures which are considered more closely indicative of pain (tender joint counts, patient global assessment) did not change (Gullick et al., 2019). Similarly, in a cohort of 477 people with early RA, established in 2005, ≥30% still experienced unacceptable pain after 15 years (Svensson et al., 2020). Those with unacceptable pain, defined as ≥40mm on a 0-100mm visual analogue scale (VAS), also displayed the most substantial impairments of functions of daily life (Svensson et al., 2020). Another study confirms this, with moderate to severe pain in RA (VAS ≥40mm) associated with worse quality of life and higher depression and anxiety scores (Vergne-Salle et al., 2020). Indeed, improvement in pain is consistently ranked as one of the top priorities in treatment outcomes by people living with RA (Sanderson et al., 2010, Hewlett et al., 2005, Carr et al., 2003). These data highlight that pain is a critical unmet clinical need in the management of RA.

Pain in RA is complex and has predominantly been linked, at least in the early stages of disease, to inflammation-driven pain referred to as nociceptive pain. When pain becomes chronic, it is often considered an issue of maladaptive processing within the central nervous system (CNS), so-called ‘central sensitisation’. However, various strands of evidence are emerging that indicate that persistent pain in RA, for a subset of patients, can still be driven by peripheral mediators. Previous work from our group utilised lidocaine injections to better assess peripherally mediated pain in a cohort of patients with active inflammatory arthritis (IA). We observed that people who lack other indicators of possible central sensitisation, like fibromyalgia, experienced better temporary pain relief in the injected joint (Rutter-Locher et al., 2024b). Further analysis into the complex mechanisms of pain within 158 individuals with active RA (DAS28 >3.2) revealed an unexpected group (27.2%), who did not have fibromyalgia or joint inflammation as assessed by ultrasound, but exhibited similar tender joint counts and VAS scores to people with joint inflammation (Chaabo et al., 2024). Multiple other studies have observed this uncoupling of pain and inflammation, when measured clinically, systemically or via joint ultrasound (Bai et al., 2024, Orange et al., 2018, Sanderson et al., 2010, Svensson et al., 2020, Vergne-Salle et al., 2020). One potential mechanistic explanation for this disconnect, beyond central sensitisation, is the pro-algesic potential of connective tissue cells, like fibroblasts.

Fibroblast dysfunction is not captured in routine clinical measures of inflammation in arthritis. And yet, when assessed, it is frequently observed in RA joints (Zhang et al., 2023) and has been shown to be an important driver of chronic inflammation in the disease (Rutter-Locher et al., 2024a, Wei et al., 2020, Mizoguchi et al., 2018, Marsh et al., 2021). More recently, synovial fibroblasts have been linked to pain in an RA cohort that displayed low joint inflammation (Bai et al., 2024). Consistent with this, previous work from us and others has shown that RA synovial fibroblasts treated with inflammatory cytokines are a major source of possible pain mediators such as interleukin-6 (IL-6), interleukin-11 (IL-11), leukemia inhibitory factory (LIF) and type 1 interferons (Li et al., 2025a, Okamoto et al., 1997, Nguyen et al., 2017). We recently showed that human sensory neurons express receptors for these cytokines and that direct stimulation of neurons with these cytokines can induce neuronal JAK/STAT3 signalling (Li et al., 2025a). Furthermore, both IL-6- and LIF-induced neuronal pSTAT3 and subsequent pain-associated gene upregulation was blocked by JAK inhibitors (JAKi), such as tofacitinib, upadacitinib and baricitinib (Li et al., 2025a). JAKi are a current treatment option for patients with RA, and post hoc analyses of clinical trials indicate that JAKi may be superior to adalimumab at reducing pain in IA (Fleischmann et al., 2019, Dougados et al., 2022, Fautrel et al., 2019, Taylor et al., 2026).

IL-6 is probably best-known as a pain-mediator, having been studied in the context of neuronal hyperexcitability and chronic pain conditions for many years (Sommer and Kress, 2004, Schaible, 2014, Choy and Calabrese, 2018, Kelleher et al., 2017). LIF is better known for its importance in sensory neuron differentiation and outgrowth (Ostasov et al., 2015, Hendry et al., 1992, Murphy et al., 1994, Cafferty et al., 2001). Literature on its role in pain is conflicting, with some reporting that LIF can induce mechanical sensitivity in rats (Thompson et al., 1996), while others suggesting that its actions might be analgesic (Banner et al., 1998). Meanwhile, IL-11 remains largely unstudied in the context of pain. Here, we focus our attention on LIF, demonstrating that blockade of LIF can inhibit most RA SF-induced neuronal pSTAT3, that LIF induces profound gene expression changes consistent with peripheral neuron sensitisation and that recombinant LIF injections can increase rodent responsiveness to joint pressure stimuli.

## Results

### LIF is a dominant driver of neuronal pSTAT3 and pSTAT1 induction in the context of RA

We previously showed that IL-6, IL-11 and LIF are enriched in RA SF and that all three cytokines and RA SF can enhance pSTAT3 levels in iPSC-derived sensory neurons (Li et al., 2025a). Here, we additionally show that LIF, IL-6+sIL-6R and IL-11+sIL-11R can induce pSTAT1 in iPSC-derived sensory neurons, and that LIF is the strongest inducer of both pSTAT3 and pSTAT1 compared to IL-6 and IL-11 even when added to the sensory neurons at 10 fold lower dose (10ng/ml LIF vs 100ng/ml IL-6 and IL-11) (Supplementary Figure 1).

This finding led us to investigate whether LIF is the dominant driver of RA SF-induced neuronal pSTAT3 compared to IL-6 and IL-11. For this, we cultured iPSC-derived sensory neurons with RA SF (n=8 donors) in the absence or presence of anti-LIF, anti-IL-11 or tocilizumab, which blocks the IL-6 receptor. A significant increase in neuronal pSTAT3 was seen in the SF only control compared to the media only control (Figure 1A, B), supporting our previous observations that RA SF can directly induce pSTAT3 in sensory neurons (Li et al., 2025a). A similar effect was observed for pSTAT1 induction (Supplementary Figure 2). Furthermore, we observed the addition of anti-LIF antibody (Ab) was the most effective at reducing RA SF-induced pSTAT3 levels, compared to anti-IL-11 or tocilizumab (Figure 1A, B) or isotype control Ab (Supplementary Figure 3).

**Figure 1:**
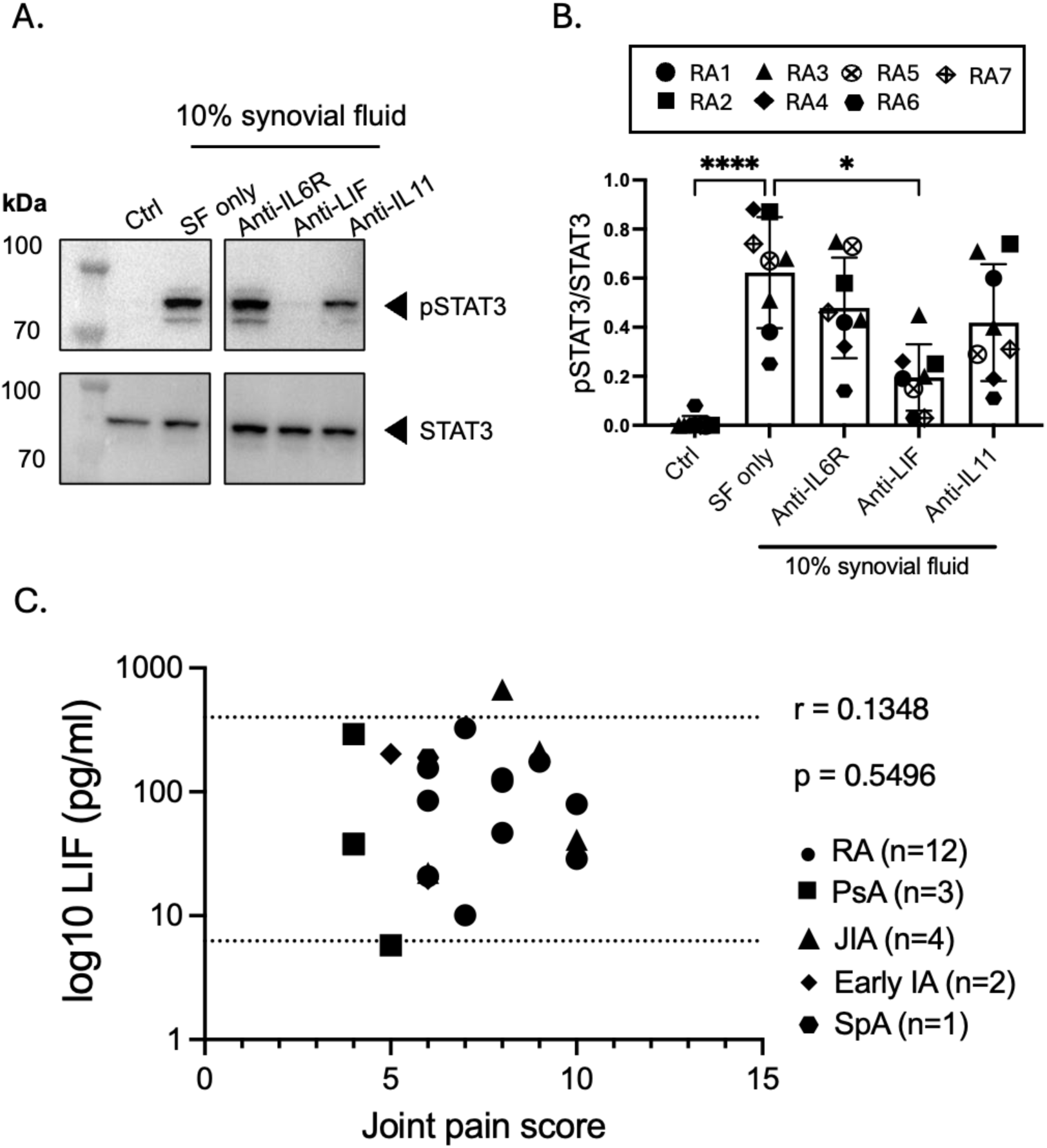
Anti-LIF is the most effective at reducing RA SF-induced neuronal pSTAT3. **A.** Representative Western blot showing pSTAT3 levels in iPSC-derived sensory neurons after a 1hr incubation with 10% v/v RA SF alone or with anti-IL-6R (100 μg/ml), anti-LIF (4 μg/ml) or anti-IL-11 (6 μg/ml) and B. corresponding densitometry of n=5 different neuronal differentiations, across n=3 different iPSC lines (UKB, K4 and AD2) exposed to a total of n=7 RA SF donors. Statistical analysis was performed using a Kruskal Wallis test followed by Dunn’s multiple comparison test comparing means to the mean of SF only condition, *padj* = <0.0001, *padj* = 0.0276. C. LIF levels within SF from patients with inflammatory arthritis (n=22) plotted against joint pain score (NRS 1-10) of aspirated knee joint (dotted lines indicate upper and lower detection point of ELISA; Spearman correlation test was applied but no significant correlation was observed). RA = rheumatoid arthritis, PsA = psoriatic arthritis, JIA = juvenile idiopathic arthritis, IA = inflammatory arthritis, SpA = spondyloarthritis.

To deduce whether LIF levels are associated with joint pain, we collected synovial fluid from people living with inflammatory arthritis (IA) who were asked to rate their pain within the aspirated knee, using a numerical rating scores (NRS) between 1 and 10. LIF was detectable in all but one SF sample (Figure 1C), and all patients reported a joint pain NRS of ≥3, which is considered unacceptable pain (Svensson et al., 2020, Tubach et al., 2012, Salaffi et al., 2015). We failed to observe a direct correlation between LIF levels and joint pain scores, but this was not surprising, given that our sample size of n=22 would only have permitted detection of large effects.

Synovial fibroblasts are one of the cell types present within the RA joint that could be responsible for producing and secreting LIF into the synovial fluid (Li et al., 2025a, Okamoto et al., 1997, Nguyen et al., 2017). When RA synovial fibroblasts were stimulated with the inflammatory cytokines IL-1β, or a combination of TNFα and IL-17A, we observed a significant increase in LIF compared to the unstimulated control (Figure 2A). Transfer of 10% (v/v) of the conditioned media (CM) from these cytokine-treated RA synovial fibroblasts onto human iPSC-derived sensory neurons led to a significant increase in neuronal pSTAT3 (Figure 2B, C). This CM-induced increase in neuronal pSTAT3 was completely abolished by the addition of anti-LIF Ab, an effect comparable to the blocking effects of the JAKi tofacitinib or the gp130 inhibitor SC144 (Figure 2D, E). These results were replicated for neuronal pSTAT1 (Supplementary Figure 4).

**Figure 2:**
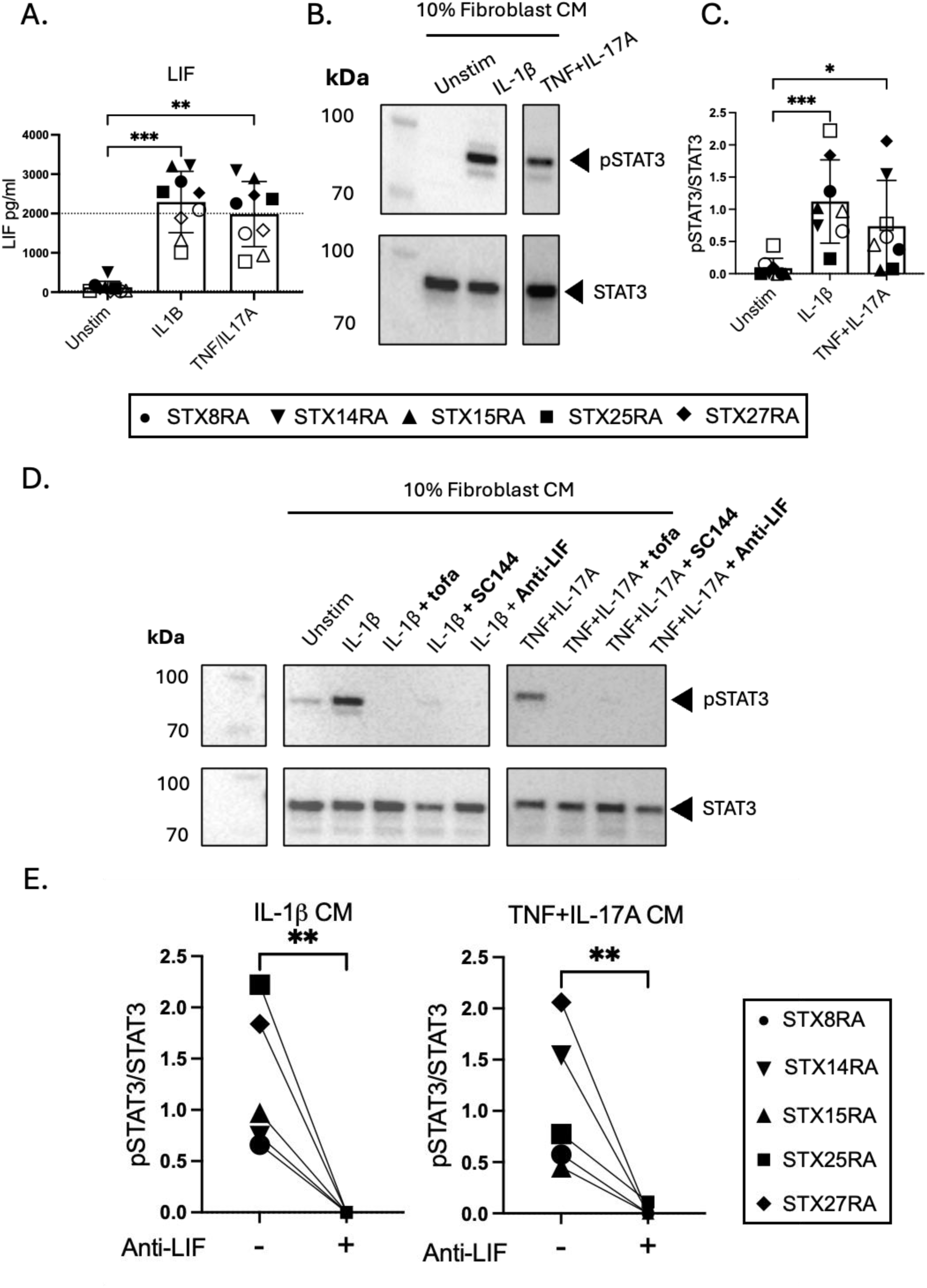
Anti-LIF blocks RA synovial fibroblast CM-induced neuronal pSTAT3. **A.** Quantities of LIF within RA synovial fibroblast CM after treatment with IL-1β (10 ng/ml) or TNF*a* (10 ng/ml) with IL-17A (10 ng/ml) for 24 hrs. Fibroblasts were obtained from n=5 different RA donors, indicated by the different solid symbols with technical repeats of the same cells indicated by open symbols. The dotted line indicates the upper detection limit of the ELISA. Statistical analysis was performed using a Kruskal Wallis test followed by Dunn’s multiple comparison test comparing means to the mean of unstimulated fibroblast CM condition, *padj* = 0.0002, *padj* = 0.0020. B. Representative Western blot showing pSTAT3 levels in iPSC-derived sensory neurons after 1hr exposure to 10% v/v conditioned media from unstimulated or cytokine treated fibroblasts, and C. corresponding densitometry of n=5 RA synovial fibroblast CM indicated by the different solid symbols with technical repeats of the same cells indicated by open symbols across n=5 different neuronal differentiations in n=1 iPSC line (K4). Statistical analysis was performed using a Kruskal Wallis test followed by Dunn’s multiple comparison test comparing means to the mean of the unstimulated CM condition, *padj* = 0.0008, *padj* = 0.0315. D. Representative Western blot showing pSTAT3 levels in sensory neurons after 1hr exposure to 10% v/v conditioned media from unstimulated or cytokine treated fibroblasts, with anti-LIF (4 μg/ml), the JAKi tofacitinib (2 μM) or the gp130 inhibitor SC144 (20 μM) and E. corresponding densitometry analysis of n=5 RA synovial fibroblast CM across n=3 different neuronal differentiations in n=1 iPSC line (K4). Statistical analysis was performed using a Mann-Whitney U test, *p* = 0.0079, *p* = 0.0079.

Collectively, these data indicate that LIF is a dominant mediator in driving neuronal pSTAT3 and pSTAT1 in the context of RA.

### LIF drives the expression of genes linked to neuronal hyperexcitability and peripheral sensitisation in iPSC-derived sensory neurons in a JAKi-dependent manner

We sought to determine how sensory neurons are transcriptionally affected by LIF as compared to other members of the IL-6 cytokine family. For this, we performed bulk RNA-seq of iPSC-derived sensory neurons that were incubated for 24 hours with recombinant LIF, IL-6 + sIL-6R or IL-11 + sIL-11R in the absence or presence of tofacitinib (Figure 3A; Supplementary Figure 5A & 6A). We performed pairwise comparisons between the LIF, IL-6 and IL-11 conditions versus media-only control and utilised a cut-off of FDR ≤ 0.05 to indicate differential gene expression. LIF induced up-regulation of a large number of genes (n=1,441), a strikingly prominent number of which have been linked to peripheral neuron sensitisation and hyperexcitability (Figure 3B, Supplementary Excel Table 1). These included *TRPV1* and *TRPV2,* ion channels known for detecting noxious stimuli such as capsaicin and high temperature (Julius, 2013, Mickle et al., 2015) and have been prominently linked to peripheral neuron sensitisation (Li et al., 2025b, Moriyama et al., 2005). There were also receptors for the constituents of so-called ‘inflammatory soup’, a substance shown to induce pain experimentally in humans (Steen et al., 1996): *NTRK1,* which encodes the high-affinity NGF receptor TrkA, and *HRH1,* which encodes the histamine receptor 1. Finally, various sodium and potassium channel genes involved in sensory neuron hyper-excitability were also upregulated, most prominently *SCN10A*, encoding for the sodium channel subunit Na_v_1.8, the target of the novel FDA-approved analgesic suzetrigine (Bertoch et al., 2025), and *SCN5A*, encoding for the sodium channel subunit Na_v_1.5.

**Figure 3:**
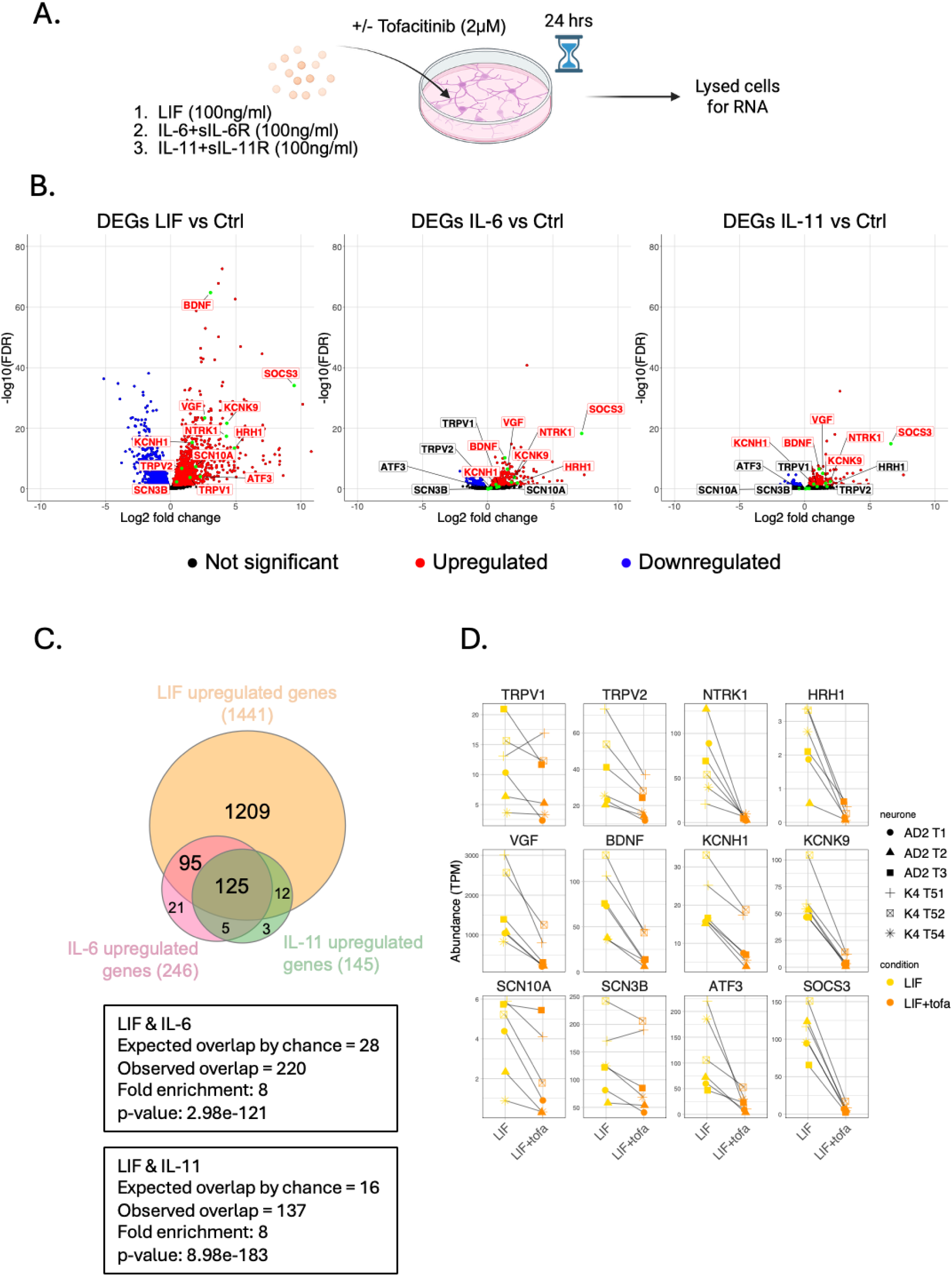
LIF signalling is a dominant driver of differential gene expression in iPSC-derived sensory neurons compared to IL-6/sIL-6R and IL-11/sIL-11R signalling. **A.** Schematic showing experimental design (n=6 neuronal differentiations, n=2 iPSC lines). B. Volcano plots showing differentially expressed genes from three different pairwise comparisons: LIF vs Ctrl, IL-6+sIL-6R vs Ctrl and IL-11+sIL-11R vs Ctrl (significant DEGs FDR ≤ 0.05, Wald test). Hyperexcitability and/or neuronal sensitisation-associated genes are highlighted with a green dot and either blue (downregulated), red (upregulated), or black (non-significant) text. C. Venn diagram showing the overlap between IL-6+sIL-6R upregulated genes, IL-11+sIL-11R upregulated genes and LIF upregulated genes. Statistical analysis was performed using a Fisher’s exact test on the gene overlap between IL-6 and LIF and IL-11 and LIF. D. Neuron hyperexcitability/sensitisation-associated genes significantly upregulated by LIF were also significantly downregulated by the JAKi tofacitinib (tofa) at padj < 0.05, with the exception of TRPV1 and SCN3B. Different symbols indicate n=3 different neuronal differentiations each, from two different iPSC lines (AD2 and K4).

In addition, and in support of our previous observations (Li et al., 2025a), *BDNF* and *ATF3* were upregulated alongside the well-known STAT3 regulated gene and JAK/STAT signalling inhibitor *SOCS3*. In contrast, IL-6 and IL-11 upregulated fewer genes (n=246 and n=145, respectively), with only a subset of the aforementioned neuron hyperexcitability/sensitisation genes affected. The overlap between LIF, IL-6 and IL-11 regulated genes was substantial, with almost all IL-6 and IL-11 regulated genes also regulated by LIF (Figure 3C). A Fisher’s exact test indicated 8x more genes overlapped between LIF and IL-6, and LIF and IL-11 than expected by chance (p-values of p = 2.98e^-121^ and p = 8.98e^-183^, respectively, Figure 3C). Critically, all these neuron hyperexcitability/sensitisation-associated genes were downregulated in the presence of tofacitinib, indicating a JAK-dependent mechanism for the LIF-induced upregulation (Figure 3D). This supports our findings that compared to IL-6 and IL-11, LIF has a greater impact on JAK/STAT signalling and on downstream pain-relevant gene transcription in iPSC-derived sensory neurons.

### LIF drives the expression of genes linked to neuronal hyper-excitability and sensitisation in iPSC-derived sensory neurons in a RA context

To further elucidate LIF-regulated gene expression within an RA context, we performed bulk RNA-seq on iPSC-derived sensory neurons that were cultured with RA SF either in the presence of anti-LIF or isotype control Abs (both at 4 μg/ml) for 24 hours (Figure 4A; Supplementary Figure 5B & 6B). We performed a pairwise comparison between the RA SF with anti-LIF condition and RA SF with isotype control condition (cut-off of FDR ≤ 0.05 to indicate differential gene expression). This analysis resulted in 281 RA SF-induced differentially expressed genes (DEGs) that were downregulated in response to anti-LIF which included many of the aforementioned genes associated with peripheral neuron hyperexcitability and/or sensitisation (Figure 4B, Supplementary Excel Table 2). To robustly identify the LIF-dependent genes that were induced by RA SF, we intersected the 281 RA SF-induced anti-LIF downregulated genes with the 1,441 LIF upregulated genes which provided us with a statistically significant overlap of 136 genes (Fisher’s exact test, fold enrichment 4.1, p-value 1.12e^-53^, Figure 4C). Functional annotation analysis using DAVID (Sherman et al., 2022) revealed that many of these 136 genes were associated with NGF receptor (NTRK1) signalling, neurotrophin signalling pathways, membrane depolarisation and ion transport, further supporting the presence of peripheral neuron sensitisation (Figure 4D; Supplementary Excel Table 3). As expected, we also found several genes that were significantly enriched in the JAK/STAT signalling pathway (Figure 4D).

**Figure 4:**
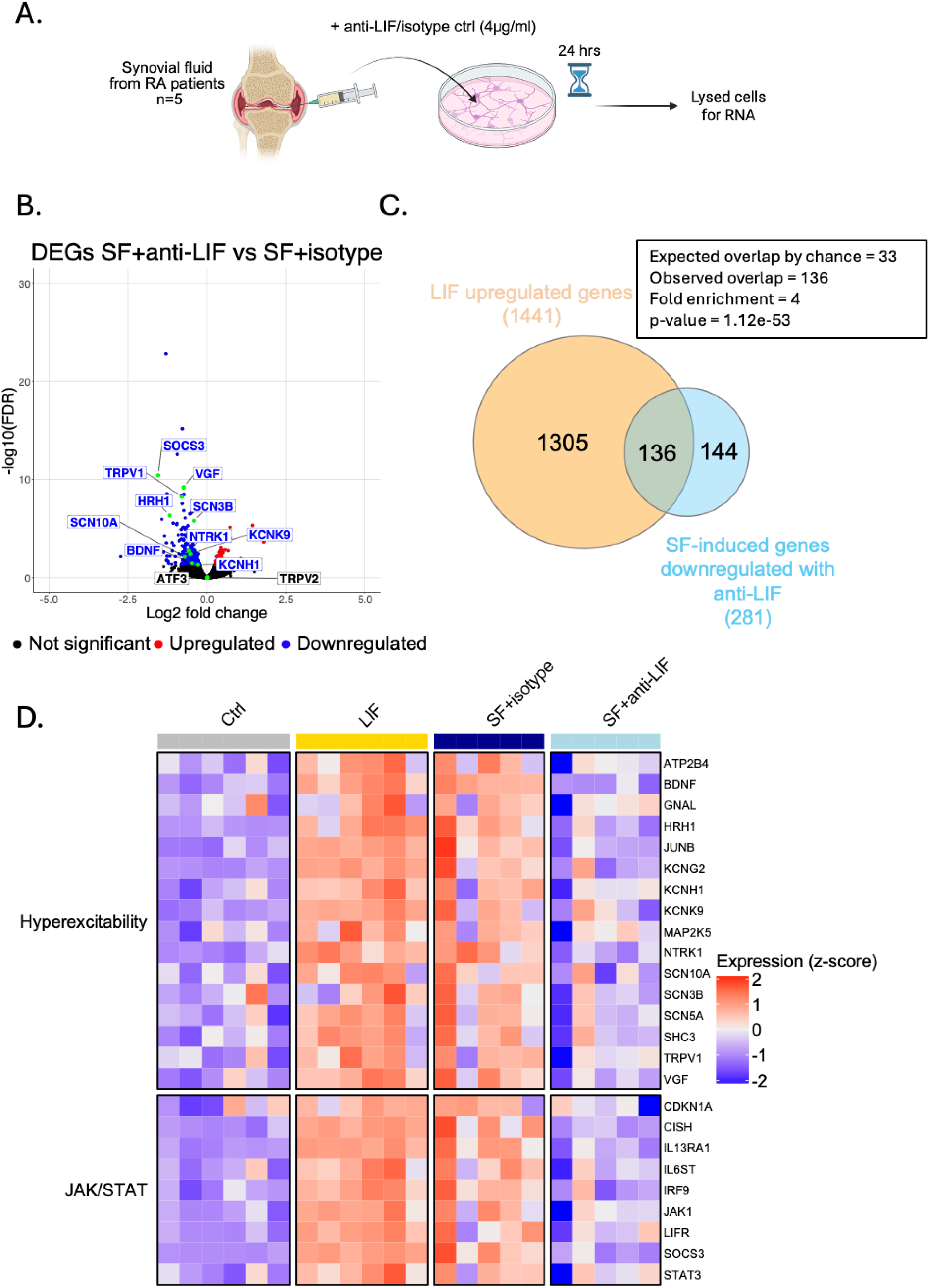
RA SF drives the expression of LIF dependent neuronal hyperexcitability/sensitisation and JAK/STAT-associated genes. **A.** Schematic showing experimental design. B. Volcano plots showing differentially expressed genes from pairwise comparisons: RA SF with anti-LIF vs RA SF with isotype control (significant DEGs FDR ≤ 0.05, Wald test). Hyperexcitability and/or neuronal sensitisation-associated genes are highlighted with a green dot and either blue (downregulated), red (upregulated), or black (non-significant) text. C. Venn diagram showing the overlap between LIF upregulated genes and RA SF induced genes downregulated with anti-LIF. Statistical analysis was performed using a Fisher’s exact test. D. Heatmap of genes identified as enriched in DAVID pathway analysis and associated with neuronal hyperexcitability/sensitisation or the JAK/STAT pathway. Only LIF-dependent genes significantly regulated in RA SF are plotted. For control and LIF conditions, each square is an independent iPSC-differentiation from n=2 iPSC lines. For SF conditions, each square is the same iPSC-derived sensory neuron differentiation (AD2 line) treated with n=5 different SF in the presence of either isotype control or anti-LIF Abs.

We then explored how LIF produced by RA synovial fibroblasts would affect the transcriptional profile of iPSC-derived sensory neurons. For this, we performed bulk RNA-seq of iPSC-derived sensory neurons that were cultured with CM from unstimulated or IL-1β treated RA fibroblasts with or without addition of anti-LIF for 24 hours (Figure 5A; Supplementary Figure 5C & 6C). Pairwise comparison between the IL-1β CM and the unstimulated CM revealed 152 upregulated DEGs including *BDNF, VGF, HRH1, KCNK9, KCNH1* and *SOCS3* (Supplementary Figure 7A, Supplementary Excel Table 4), whilst 113 genes were downregulated when comparing the IL-1β fibroblast CM in the presence vs. the absence of anti-LIF conditions; these included *BDNF, VGF, HRH1 and SOCS3* (Supplementary Figure 7B). We took forward the 55 intersecting genes from the two comparisons (fold enrichment 44.4, p-value 1.25e^-80^, Supplementary Figure 7C), which we considered to be robustly induced by the IL-1β fibroblast CM in a LIF-dependent manner.

**Figure 5:**
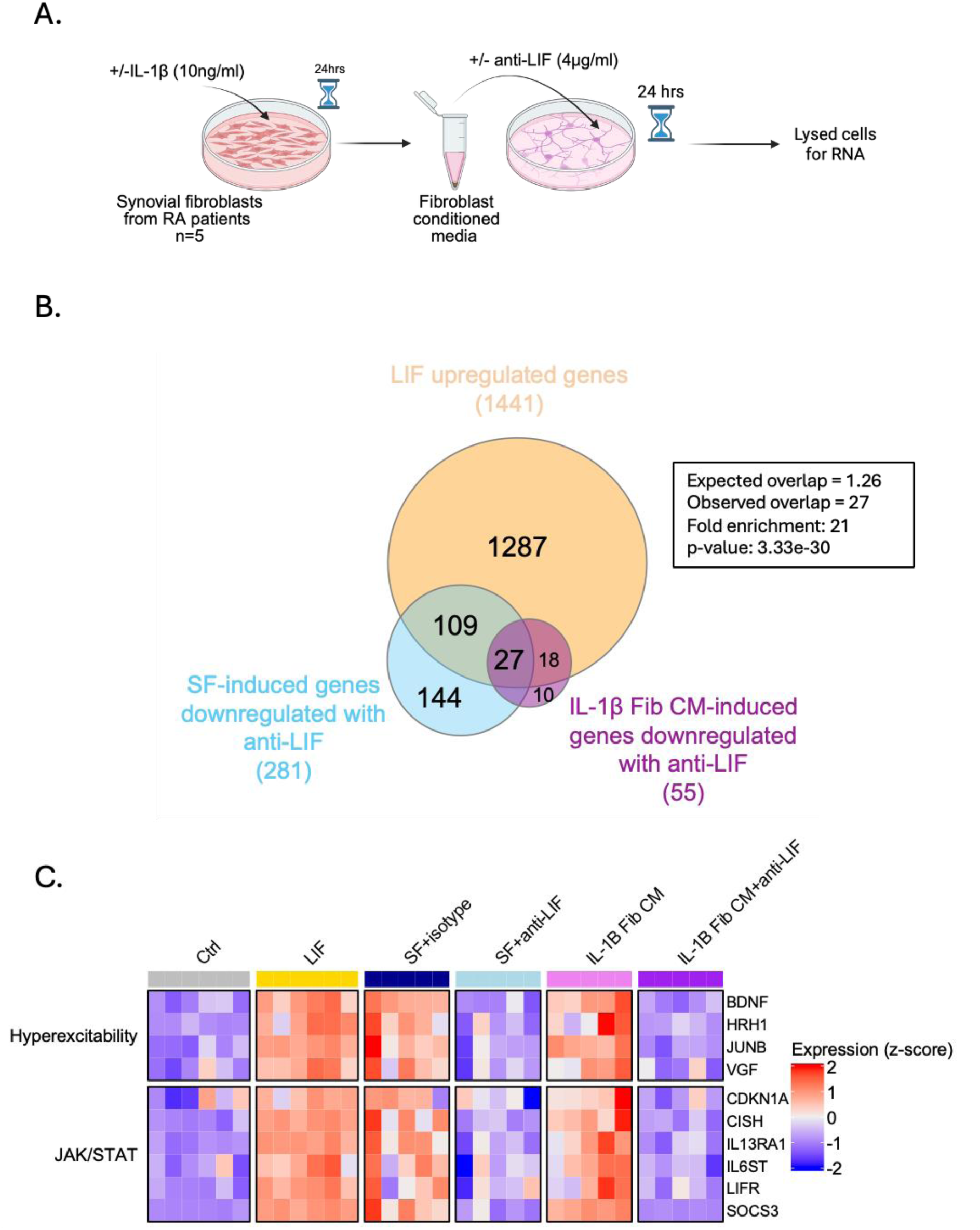
LIF dependent neuronal hyperexcitability/sensitisation and JAK/STAT associated genes are also induced by conditioned media from cytokine treated RA synovial fibroblasts. **A.** Schematic showing experimental design. B. Venn diagram showing the overlap between LIF significantly upregulated genes, RA SF-induced genes significantly downregulated with anti-LIF and IL-1β-treated fibroblast CM-induced genes downregulated with anti-LIF. Statistical analysis was performed using a super exact test. C. Heatmap of neuron hyperexcitability/sensitisation-associated genes and JAK/STAT associated genes. Only LIF-dependent genes induced by both RA SF and RA synovial fibroblast CM are plotted. For control and LIF conditions, each square is an independent iPSC-differentiation from n=2 iPSC lines. For SF conditions, each square is the same iPSC-derived sensory neuron differentiation (AD2 line) treated with n=5 different SF in the presence of either isotype control or anti-LIF Abs. For the IL-1β fibroblast CM, each square is n=5 different RA synovial fibroblast with or without anti-LIF added to n=3 neuronal differentiations across n=1 iPSC line (AD2).

We then integrated the three components of our bulk RNA-seq analysis by identifying the intersection between the LIF upregulated genes, the RA SF-induced genes downregulated by anti-LIF and the synovial fibroblast IL-1β CM-induced genes downregulated by anti-LIF. This resulted in a statistically significant overlap of 27 genes (Fisher’s exact test, fold enrichment 21, p-value 3.33e^-30^, Figure 5B). Of these 27 genes, 10 overlapped with the genes from the above-mentioned functional annotation analysis by DAVID (Figure 5C). Together, this analysis demonstrates that RA SF and RA synovial fibroblasts induce the expression of genes associated with neuronal hyperexcitability and/or sensitisation in a LIF-dependent manner.

We validated our key findings in the HD10.6 cell line. HD10.6 is a conditionally immortalised human dorsal root ganglion (DRG)-derived neuronal cell line established from embryonic male human DRG tissue through transfection with a tetracycline-regulated *v-Myc* expression construct (Raymon et al., 1999). It has recently been shown by us and others (Dhillon et al., 2026, Dochnal et al., 2026, Al-Abbasi et al., 2025) that prolonged maturation of these cells generates human sensory neurons that more faithfully recapitulate some features of nociceptive sensory neurons than iPSC-derived models; this includes higher TRPV1 expression. Using this independent cell model, we confirmed by Western blot and qPCR that LIF induces pSTAT3 and pSTAT1 within differentiated HD10.6 neurons (Supplementary Figure 8A-D) and upregulates some of the top LIF-dependent genes (*SOCS3*, *LIFR* and *HRH1*) from our RNA-seq analysis (Supplementary Figure 8E).

### LIF increases neurite outgrowth after axotomy

Changes in joint innervation during arthritis remain an active area of investigation (Pongratz and Straub, 2013, Xenophontos et al., 2025), not least since it would likely accompany vascular remodelling that occurs in affected joints (Buckley et al., 2006). Since LIF is known to affect sensory neuron development and neurite outgrowth in rodents, we tested whether it might have similar effects in our human HD10.6 neurons using a microfluidic device. Cells were seeded into the soma compartment of the device, and differentiation was initiated 24 hours later. The cells were then transduced with pAAV-CAG-tdTomato 7 days later to visualise all neuronal processes, and at day 21 axotomy was performed by vacuum aspiration confined to the axonal compartment followed by LIF or control treatment for 24 and 48 hours. A statistically significant increase in neurite outgrowth with LIF treatment compared to control was observed at both 24 (p-value 0.0002) and 48 hours (p-value 0.0026) after axotomy (Figure 6).

**Figure 6:**
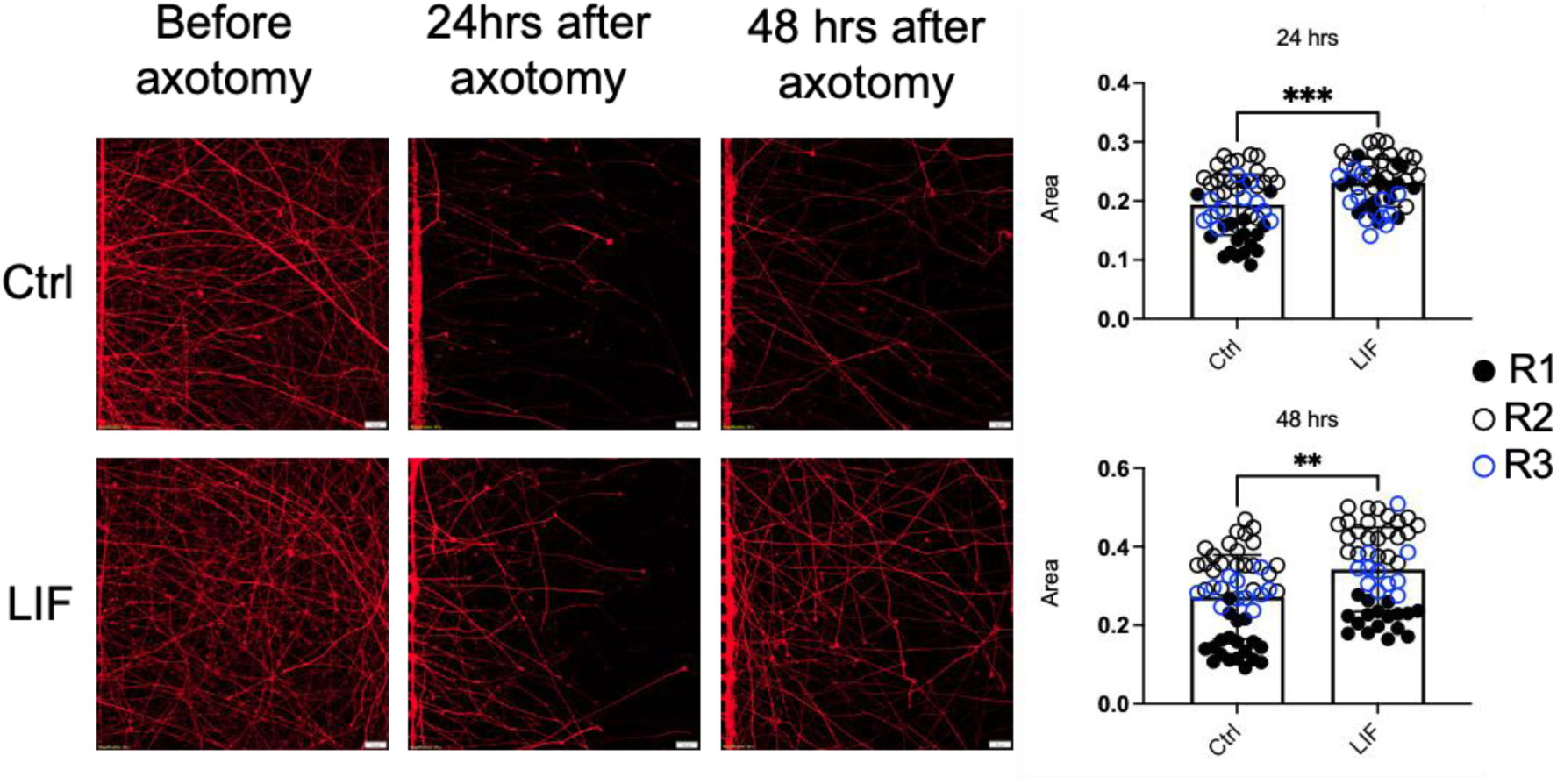
LIF increases neurite outgrowth in differentiated HD10.6 DRG neurons. Representative microscopy images of neurites before axotomy, 24 hours, and 48 hours after axotomy with and without LIF treatment alongside corresponding analysis of the area covered by tdTomato-positive neurites. Four different fields of view were assessed for n=3 independent HD10.6 differentiations (R1-3) and normalised to their corresponding pre-axotomy values. Each dot is a field of view, with different differentiations indicated by colour. Statistical analysis was performed using a Mann-Whitney U test (*p* = 0.0002, *p* = 0.0026).

### LIF is induced in the joints of arthritic mice and, in healthy animals, causes neuronal pSTAT3 induction and increased sensitivity to painful joint pressure

Finally, to examine whether LIF plays a role in a whole animal setting, we conducted a series of experiments in mice. We were able to show that, as in humans, LIF is significantly increased in the affected joint after induction of antigen-induced arthritis (Figure 7A, B). As expected, this was accompanied by joint swelling and a reduction in weight bearing on the affected leg, indicative of pain (Figure 7C, D). We also confirmed that akin to their human counterparts, mouse knee synovial fibroblasts release LIF in response to IL-1β stimulation *in vitro* (Supplementary Figure 9A & B) and that cultured mouse DRG neurons respond to stimulation with LIF, as observed by an increase in pSTAT3 (Supplementary Figure 9C & D).

**Figure 7:**
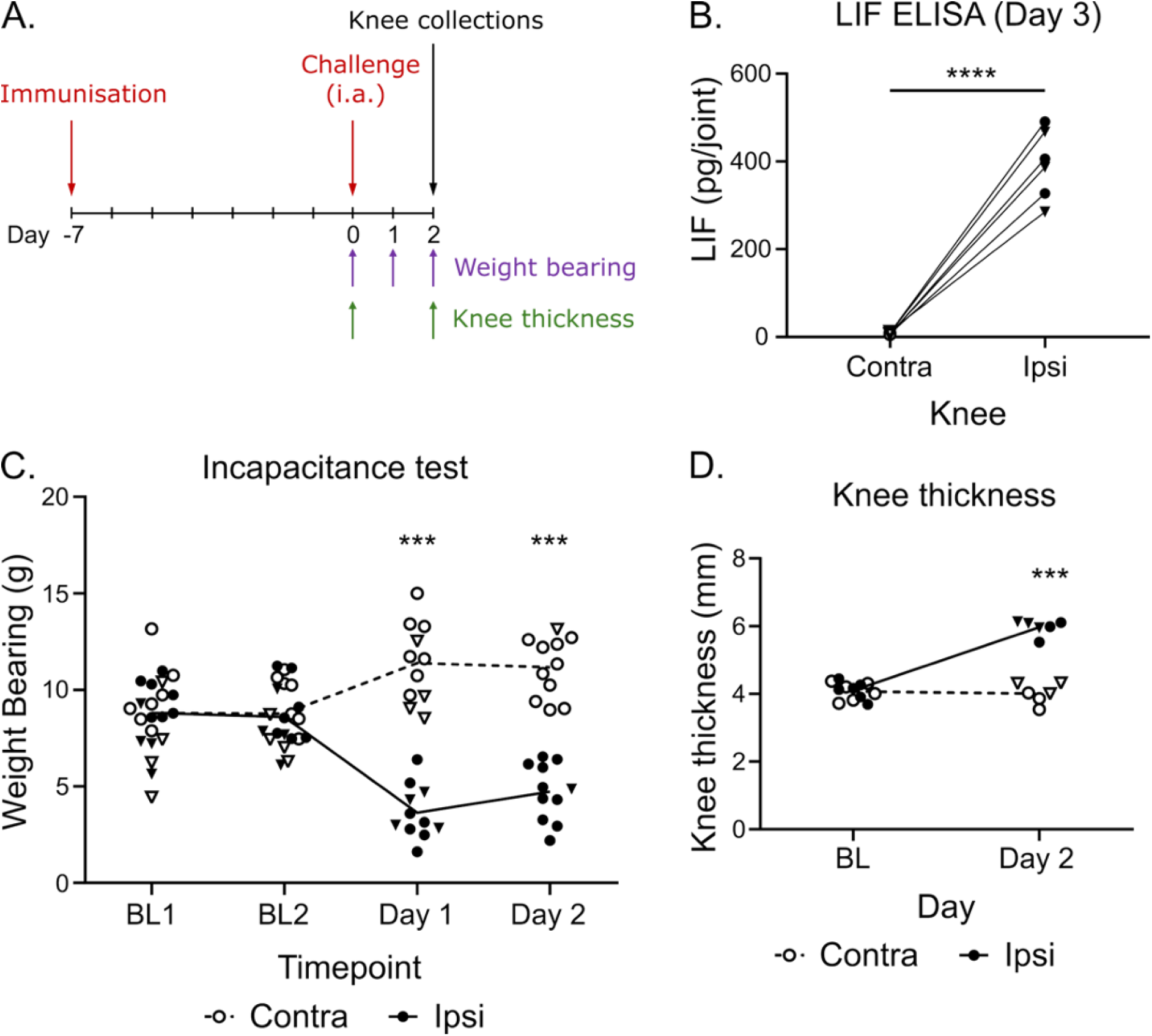
Inflammation and pain in a mouse model of inflammatory arthritis is accompanied by an increase in LIF in the affected joint. **A.** Schematic representation of experimental plan. Briefly, antigen induced arthritis (AIA) was induced in wildtype mice, and an incapacitance test to measure weight bearing was carried out daily. Knee thickness was also measured on the day of arthritis induction (day 0) and the day of peak inflammation (day 2). i.a. = intraarticular. B. LIF measured via ELISA in the supernatant of digested AIA mouse knees, 3 days after the induction of arthritis. n=6. Paired t test comparing ipsi-to contralateral knees (p < 0.0001). C. Weight placed on each leg in AIA mice (n=11), measured using the incapacitance test; two baseline measurements taken on day 1 and day 2; 2-way ANOVA with Turkey’s multiple comparisons test (day: p = 0.0009, knee: p < 0.0001, day x knee: p < 0.0001). D. Knee thickness of the contralateral and ipsilateral leg of AIA mice on day 0 (BL) and day 2 after induction of the model; n=6, paired t-test between ipsi and contra knees at day 2 (p < 0.0001). Circles = males, triangles = females.

To investigate whether the cytokine has an effect on pain in healthy mice, 0.5 μg of LIF was injected into the joints of mice and successful delivery was confirmed via ELISA (Figure 8A, Supplementary Figure 10A & B). LIF did not alter weight bearing, i.e. likely failed to elicit spontaneous sensory neuron firing (Supplementary Figure 10C & D). However, in line with what we would expect from the human gene expression results presented here, LIF did appear to induce functional peripheral neuron sensitisation; specifically, mice that had received intra-articular LIF rather than vehicle injections more readily responded to painful joint pressure (Figure 8B, 2-way ANOVA, significant interaction between day and treatment, p = 0.04, with Sidak’s multiple comparison tests revealing the most prominent difference to be at 72h, p = 0.0012). In these same mice, nerves innervating the knee joint were retrogradely labelled with Fast Blue (FB) dye to identify them in subsequent immunofluorescence staining for pSTAT3. The absolute number of joint-innervating neurons per sensory ganglion is very small, thus increasing variability between mice; in spite of this, and matching our *in vitro* observations, more than 60% of FB+ joint neurons were pSTAT3+ in 4 out of 6 LIF injected mice, compared to a maximum of 50% in all vehicle-injected mice (Figure 8C, D). Concordantly, there were significantly fewer pSTAT3 negative joint neurons in LIF-treated compared to vehicle mice (p = 0.0097, Supplementary Figure 10E-H).

**Figure 8:**
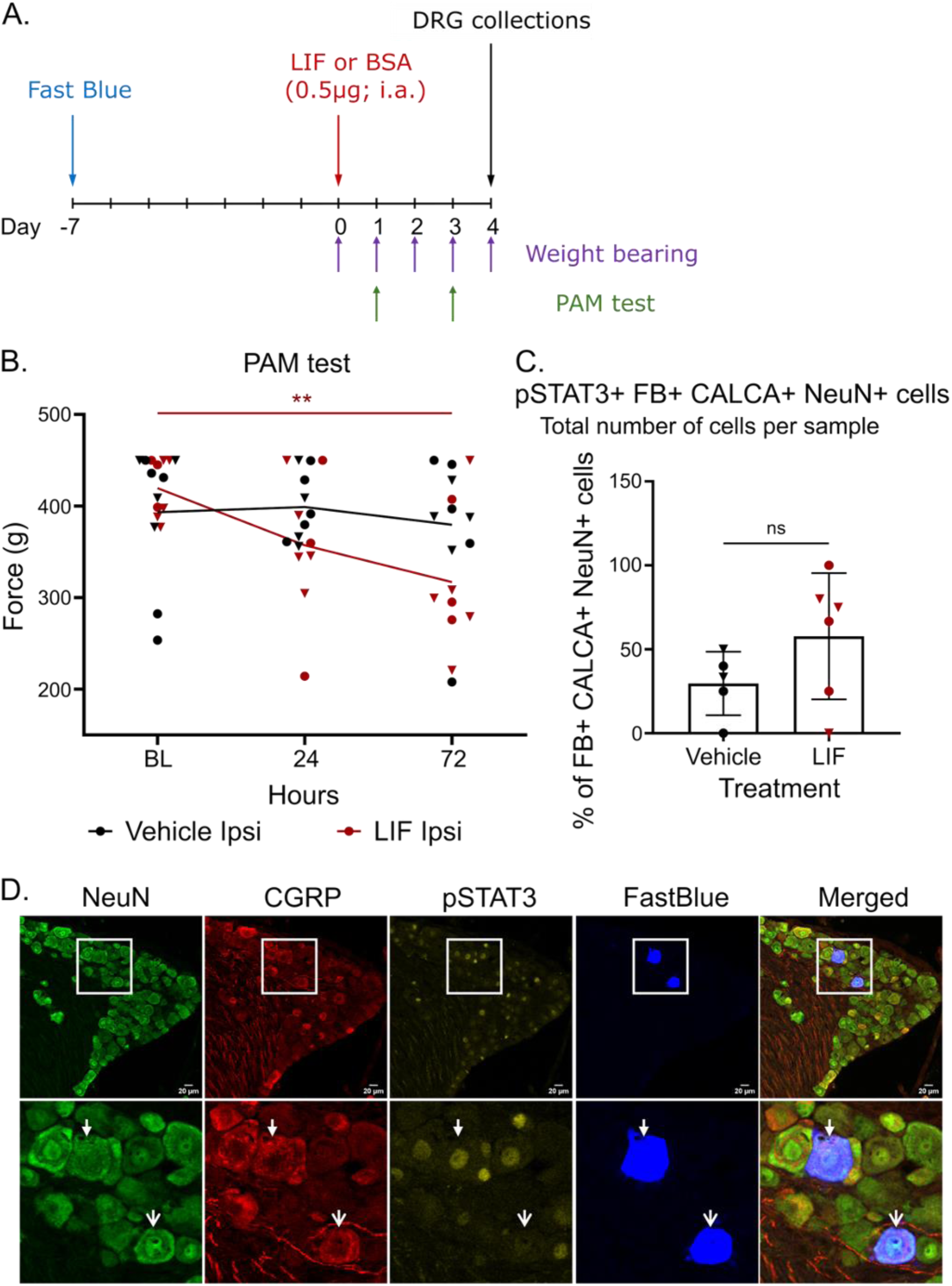
Intraarticular injection of LIF results in a reduced joint pressure pain threshold in healthy mice. **A.** Schematic representation of experimental plan: Fast Blue (FB) was injected intraarticularly in the left knee (ipsilateral) of wildtype mice. Seven days later, LIF (0.5μg) or vehicle (0.1% BSA) was injected in the ipsilateral knee, and weight bearing was assessed daily until day 4 via an incapacitance test. Pressure Application Measurement (PAM) was taken for joints at baseline (BL) and on days 1 and 3 (24 & 72hr after injection of LIF). Sensory ganglia (DRG) were collected on day 4 for immunohistochemistry. i.a. = intraarticular. B. PAM test results for ipsilateral knees, showing at which force a withdrawal response was elicited. Each symbol represents an individual mouse (circles = males, triangles = females; vehicle: n=9, LIF: n=8). 2-way ANOVA with Šídák’s multiple comparisons test (interaction between day x treatment, p = 0.04; baseline vs. LIF at 72h, p = 0.0012). C. Percentage of pSTAT3 positive cells as a proportion of all Fast Blue, CALCA and NeuN triple positive cells in sensory ganglia which send projections to the joint (L4 DRG), 4 days after intraarticular injection with LIF (n=6) or vehicle (n=5). D. Representative image from the analysis in Figure 8C, showing a field of view containing a pSTAT positive (yellow, filled arrow) and a pSTAT3 negative (open arrow) Fast Blue, CALCA (red) and NeuN (green) triple positive cell.

## Discussion

Effective pain management, which is consistently ranked as a top priority for treatment improvement amongst people living with RA, is still an unmet clinical need. To address this, we need a better understanding of the mechanisms underlying chronic pain in RA. Our study has revealed LIF as a potent and prominent driver of JAK/STAT signalling in sensory neurons within an RA context. Downstream, LIF induces upregulation of genes associated with peripheral neuron sensitization, increases neurite outgrowth and decreases pressure pain thresholds *in vivo*. Altogether, these pro-algesic changes suggest that LIF may be a promising mediator for drug targeting in RA.

As reported previously (Li et al., 2025a, Okamoto et al., 1997, Nguyen et al., 2017, Lotz et al., 1992), our results confirm synovial fibroblasts as a prominent source of LIF. Importantly, we have now additionally demonstrated that fibroblast conditioned media are able to induce both pSTAT3 and pSTAT1 in human sensory neurons in a LIF-dependent manner, leading to induction of genes associated with neuronal hypersensitivity and sensitisation. This finding is particularly important in the context of chronic intractable pain, as evidence has been converging to suggest that dysfunctional fibroblasts are associated with pain in the absence of clinically measurable inflammation. This is not just true in rheumatoid arthritis (Bai et al., 2024), but also other chronic pain conditions, such as osteoarthritis and neuropathy (Shinotsuka and Denk, 2022, Kreß et al., 2021, Wijesinghe et al., 2024, Villa-Hernandez et al., 2025).

We detected LIF protein in RA joints as previously described (Li et al., 2025a) and specifically confirmed that LIF is present in painful knee joints (joint pain score of ≥3 on NRS) in a heterogeneous population of individuals with IA. Furthermore, we show that levels of LIF are significantly higher in the arthritic and painful knee of the AIA mice, making this a useful model for further study of the role LIF plays in RA pain. Additionally, we observed that joint injections of LIF significantly decreased the threshold at which healthy mice were willing to sustain a painful pressure stimulus. This finding is in line with a report of LIF inducing mechanical sensitivity when injected into the hindpaws of rats (Thompson et al., 1996). A decrease in joint pressure pain thresholds is also routinely observed in RA study cohorts (Trouvin et al., 2022, Lee et al., 2013).

In both our *in vivo* and *in vitro* model systems, we have observed that the functional effects of LIF are more pronounced after several days; concordantly, LIF-injected joints of mice were most sensitive 72h after injection, while neuronal firing rates increased most prominently in iPSC-derived sensory neurons 96h after first exposure (Li et al., 2025a). A mechanistic explanation for this phenomenon readily presents itself in our sequencing results. We show that LIF induces profound alterations in the transcriptome of sensory neurons, consistent with the cells undergoing so-called ‘peripheral sensitisation’, whereby pain-relevant ion channels and receptors are regulated in a direction that renders the neurons more sensitive to mechanical, thermal and noxious stimuli. Peripheral sensitisation is a hallmark of chronic pain states (Basbaum et al., 2009), but until now, it has been surprisingly difficult to identify mediators that can clearly induce it at transcriptional level in isolated cultured sensory neurons. For example, mediators present in the above-mentioned pain inducing inflammatory soup altered expression of just a single gene in iPSC-derived sensory neurons and did not result in convincing alterations in neuronal firing properties, neither in mouse nor human neuron models (Li et al., 2025b). To our knowledge, LIF is therefore the first cytokine able to induce peripheral sensitisation at transcriptional level in sensory neurons in the absence of any other supporting cell types. We show that its effects are more profound than IL-6 and IL-11, and crucially for RA pain, that similar LIF-dependent changes can be observed in sensory neurons exposed to human RA synovial fluid.

In conclusion, our study indicates that, in contrast to other pro-algesic cytokines, LIF is able to directly induce peripheral neuron sensitisation in models of RA, without involvement of other cell types. Its effects are more pronounced than those of IL-6 and IL-11, with anti-LIF able to reverse gene expression changes associated with peripheral neuron sensitisation induced by both RA synovial fluids and RA synovial fibroblasts. Based on this, we propose that LIF inhibition may be a promising therapeutic avenue for individuals with low inflammation but persistent peripheral pain in RA.

## Materials and Methods

**Table 1.** – Antibodies and inhibitors used within this study.

| Antibody/inhibitor | Clone | Company | Catalogue # | Concentration | Usage |
| --- | --- | --- | --- | --- | --- |
| **Human samples & model systems** |  |  |  |  |  |
| STAT3 | D3Z2G | Cell signalling | 12640S | 1 in 1000 | Western blot |
| pSTAT3 (Tyr705) | D3A7 | Cell signalling | 9145S | 1 in 1000 | Western blot |
| GAPDH | D16H11 | Cell signalling | 5174T | 1 in 1000 | Western blot |
| Anti-Rabbit HRP | - | Life Technologies | 32460 | 1 in 5000 | Western blot |
| LIF | polyclonal | R&D Systems | AF-250-NA | 4 µg/ml | Neutralisation |
| IL-11 | polyclonal | R&D Systems | AF-218-NA | 6 µg/ml | Neutralisation |
| Goat IgG | polyclonal | R&D Systems | AB-108-C | 4 µg/ml | Neutralisation |
| IFNAR1 | Anifrolumab | InVivoSIM | SIM0022 | 100 µg/ml | Neutralisation |
| IL-6R | Tocilizumab | Roche | - | 100 µg/ml | Neutralisation |
| Tofacitinib |  | Cayman Chemical | 11598 | 2 uM | Inhibition |
| SC144 |  | Sigma-Aldrich | SML0763 | 20 uM | Inhibition |
| **Mouse samples & model systems** |  |  |  |  |  |
| STAT3 | 79D7 | Cell signalling | 4904 | 1 in 1000 | Western blot |
| pSTAT3 (Tyr705) | D3A7 | Cell signalling | 9145S | 1 in 2000 | Western blot |
| pSTAT3 (Tyr705) | D3A7 | Cell signalling | 9145S | 1 in 100 | Immunostaining |
| NeuN | polyclonal | Merck | ABN91 | 1 in 750 | Immunostaining |
| CGRP | polyclonal | Enzo Life Sciences | BML-CA1137-0025 | 1 in 400 | Immunostaining |

For details of all other reagents used within this study refer to Supplementary Excel Table 5.

### Patient samples

Synovial fluid (SF) samples were obtained from patients with inflammatory arthritis attending Guy’s Hospital Rheumatology Department. Cell free SF was generated by centrifuging 1 ml aliquots of the SF at 8000 rpm for 3.5 minutes to pellet the cells then collecting and storing (-80°C) the supernatant. Synovial fibroblasts were obtained from explant cultures of synovial tissue from arthroplasties or biopsies. Once expanded, the synovial fibroblasts were frozen in FBS with 10% DMSO. Synovial fibroblasts were thawed and cultured in complete DMEM (500ml DMEM with 50ml FBS, 5ml pen/strep and 2ml fungizone) with media changes every 2-3 days. Synovial fibroblasts were split at 90% confluency which was typically once a week. The passage number for the fibroblasts used within this study ranged from passage 5 to passage 7. All patient samples were obtained with written informed consent, and the study protocols (Pain Phenotypes and The Underlying Mechanisms in Inflammatory Arthritis [PUMIA] and Psoriatic, Rheumatoid, Inflammatory arthritis; Study of Immune Mechanisms [PRISM]) received approval from the Bromley Research Ethics Committee and Health Research Authority (21/LO/0712) and the Harrow Research Ethics Committee (66106), respectively. For clinical and demographic parameters for all samples refer to Supplementary Excel Table 6.

### Differentiation of iPSC-derived sensory neurons

The sensory neurons utilised within this study were generated from three iPSC lines, Kute4 (HPS1071 4i-kute_4, originally generated by the Cambridge BioResource and Wellcome Trust Sanger Institute from skin fibroblasts of a 25-29 year old female), AD2 (STBCi321-A, formerly known as SFC-AD2-01, originally generated by the StemBANCC consortium at Oxford from skin fibroblasts of a 51-year-old male) and UKB (UKBi013-A-GCamP6f, generated via Sendai virus reprogramming of peripheral blood mononuclear cells from a healthy male donor). The iPSCs were grown and maintained on vitronectin-coated plates in Stemflex medium until 70% confluency when they were passaged using versene. iPSCs were differentiated following the Chambers protocol (Chambers et al., 2012) and replated at day 11, using 75,000 cells per geltrex-coated coverslip in 24 well plates. The neurons matured in N2 media containing NGF, NT-3, GDNF and BDNF at 25ng/ml between day 11-39; after day 39, the concentration of all growth factors was reduced to 10ng/ml. Geltrex was added to the media once a week to prevent the cells detaching. To remove any non-neuronal contaminating cells, 1-3 μM Ara C was added, usually between day 14-30. For the experiments included in this study the neuron age range was day 49 to day 95.

### HD10.6 cell culture

Cells were cultured as previously described (Raymon et al., 1999, Al-Abbasi et al., 2025, Dhillon et al., 2026) with minor modifications. In brief, cultures were maintained at 37 °C in a humidified, 5% CO₂ incubator and verified to be free of mycoplasma before any experiments were performed. To maintain proliferation, cells were grown on Cultrex-coated plates in a proliferation medium of Advanced DMEM/F-12 containing GlutaMAX, B27 supplement, prostaglandin E1, bFGF, and geneticin (G418). Once cultures reached roughly 70–80% confluency, they were passaged and re-seeded at approximately 16,000–19,000 cells/cm², with the exact density depending on the platform used (multiwell plates or microfluidic devices).

Differentiation was begun 24 hours after plating (defined as day 0) by exchanging the proliferation medium for a maturation/differentiation medium based on Neurobasal medium and supplemented with GlutaMAX, B27 supplement, tetracycline (1 µg/mL, to repress v-Myc expression), forskolin, and a panel of neurotrophic factors (NGF, GDNF, NT-3 and CNTF). Half-volume media changes were carried out every 2–4 days, and cultures were maintained for up to 21 days in vitro (DIV21).

### Western blotting

After treatment, the neurons were washed in cold 1X PBS and lysed in 1X XT sample buffer with β-mercaptoethanol and stored at -80°C. Samples were thawed, boiled at 95°C for 5 minutes and loaded onto either a 4-20% mini-PROTEAN® TGX™ gel or a 4_15% Criterion™ tris-HCl gel, depending on the number of samples. Once loaded, the gels were ran at 100V until the dye front ran off the bottom of the gel. The proteins were then transferred onto a PVDF membrane using the Bio-Rad Trans-Blot® Turbo™ system. Membranes were blocked in 5% milk in 1X TBS with 0.1% Tween® 20 (TBS-T) and incubated overnight at 4°C in primary anti-pSTAT3 or anti-GAPDH antibodies diluted in TBS-T. Membranes were then washed 3 times for 5 minutes each in TBS-T and incubated for 1 hour at room temperature in secondary antibody diluted in TBS-T. The membranes were washed again (x3 for 5 minutes) and imaged, using SuperSignal Chemiluminescent Substrates, using a Chemidoc XRS+. Blots were stripped with 1X ReBlot Plus stripping solution for 15 minutes followed by blocking, primary anti-STAT antibody incubation and the same steps as above. Densitometry for each blot was calculated in Fiji, and pSTAT values were normalised to total STAT.

### RNA extraction, cDNA synthesis and qPCR

**Table 2.**
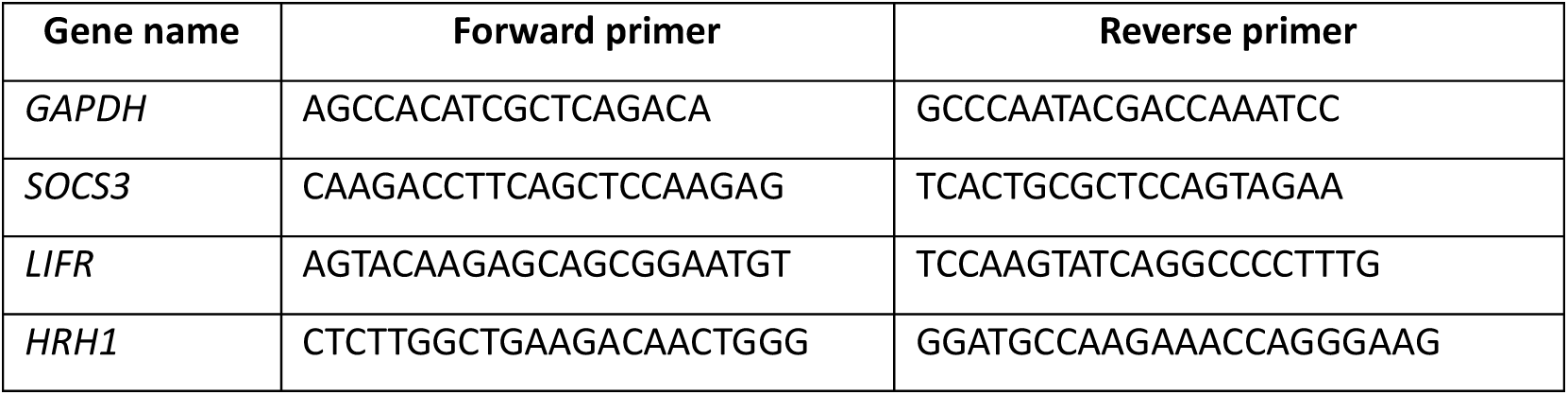
- Primer sequences used within this study.

All cells were lysed directly in 350μl RLT plus buffer with 1% β-mercaptoethanol and stored at -80°C. RNA was extracted using the RNeasy® Plus Micro kit, following manufacturer’s instructions. cDNA synthesis was performed using High-Capacity RT kit and qPCR was performed using the PowerTrack™ SYBR Green Master Mix, both following manufacturer’s instructions. Per reaction between 2-12.5ng/ml cDNA was added. Relative expression to the housekeeping gene GADPH was generated using the delta CT method.

### Fibroblast stimulation for generation of conditioned media

Between 40,000-60,000 synovial fibroblasts were seeded into a 24 well plate in complete DMEM and left overnight to recover and adhere. The following day, the media were changed and each well was supplemented with fresh DMEM media with or without IL-1β or TNF-α with IL-17A (at a final concentration of 10ng/ml) for 24 hours. After the 24 hours, the conditioned media were collected, and any technical repeats were pooled together. The conditioned media were then centrifuged at 8,000 rpm for 3.5 minutes, the supernatants collected and aliquoted into eppendorfs for storage at -80°C.

### Neuron stimulation

**Table 3.**
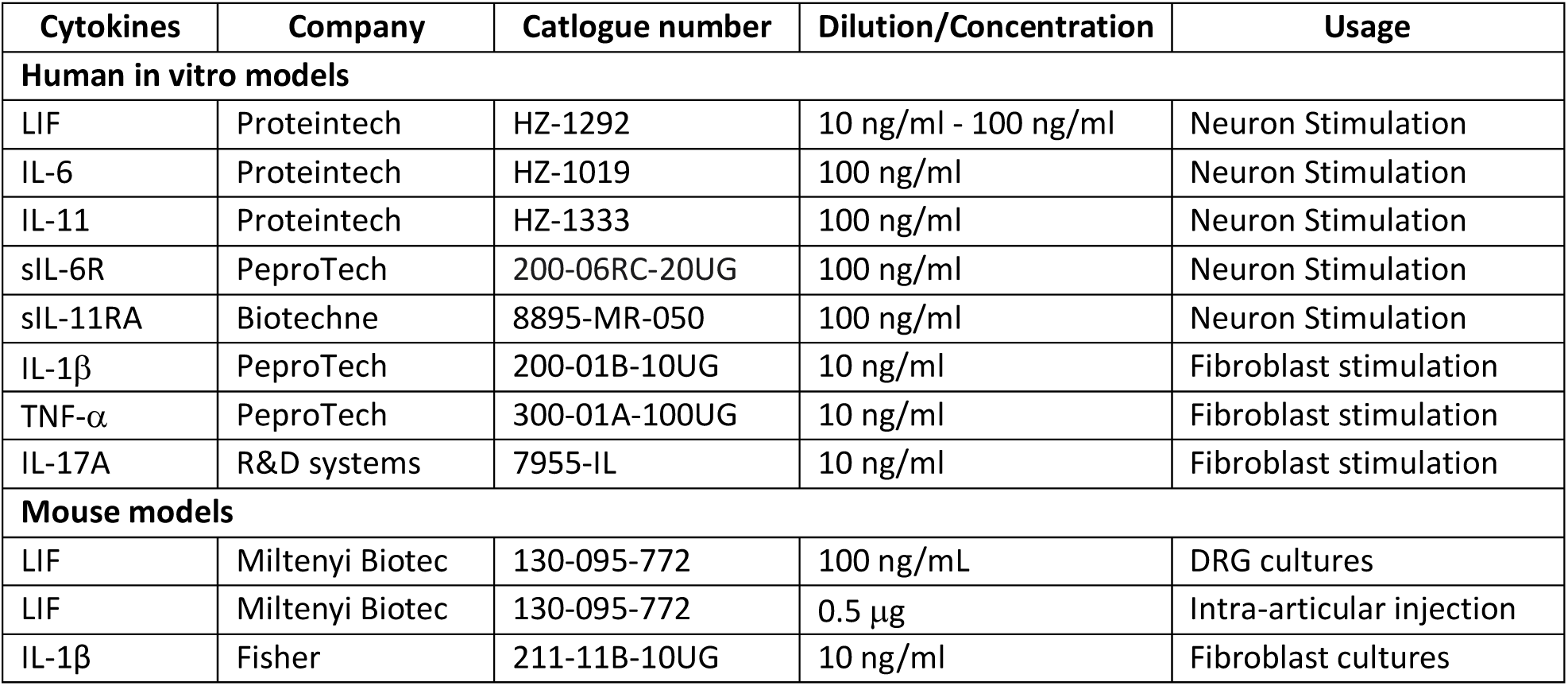
- Recombinant cytokines used within this study.

For all *in vitro* stimulations, neurons were incubated for 24 hours for RNA analysis and between 30 minutes and 1 hour for Western blotting. HD10.6 neurons required additional growth factor starvation (8 hours) prior to stimulation.

For stimulation of neurons with RA SF, the synovial fluid was first centrifuged at 16,000g for 4 minutes and diluted to 10% in N2 media. For fibroblast conditioned media stimulation, the conditioned media was diluted to 10% in complete DMEM.

Soluble receptors were pre-mixed with the relevant cytokines for 1 hour with shaking before adding to the neurons. Neutralising antibodies were pre-mixed with CM or SF for 1 hour with shaking before adding to the neurons. Tofacitinib and SC144 were pre-incubated with the neurons for 1 hour and 24 hours respectively.

Neuronal media was removed and the neurons were supplemented with 300 µl of the prepared stimulation.

### RNA-seq

Neurons were lysed in 350μl RLT plus buffer with 1% β-mercaptoethanol and stored at -80°C. RNA extraction was batch-controlled and performed using the RNeasy plus micro kit. RNA concentration was measured using the Qubit High Sensitivity kit, and RNA integrity was checked using the High Sensitivity RNA Screen Tapes and corresponding kit on the Aligent Tapestation 4200. Samples were shipped on dry ice to Novogene in a single library preparation and sequencing run (∼30M reads) using the Illumina Novaseq X Plus. Reads were pseudo-aligned to the human genome (Homo Sapiens GCRh38) using Kallisto (version 0.51.1). Genes with raw counts of ≥10 within all samples of a given group were considered expressed to filter out lowly expressed genes. A further filter, where transcript per million (TPM) values had to be ≥1 in all samples of at least 1 group was applied to the cytokine and SF analysis. These lists of all expressed genes were then also used as a background for Fisher’s exact test calculations. Differential gene expression was performed using DESeq2 in R, with neuronal differentiation, synovial fluid donor and/or fibroblast line included as factors. Genes were considered differential expressed if they had a false discovery rate (FDR) ≤ 0.05. Full scripts for all analyses are provided in the Supplement and on the Gene Expression Omnibus, where all raw and processed data have also been submitted under accession number GSE347477.

### Neurite outgrowth in HD10.6 neurons

Microfluidic devices were fabricated in-house from epoxy templates as previously described (Clark et al., 2018) and bonded to glass-bottom dishes (WillCo). Before coating with Cultrex extracellular matrix, devices were plasma-treated (Femto plasma cleaner; Diener Electronics, Germany). HD10.6 cells were introduced into the soma compartment in proliferation medium, and differentiation was initiated 24 hours later (day 0) by switching to maturation medium, with half-volume media changes thereafter every 2–4 days. At DIV7, cultures were transduced with pAAV-CAG-tdTomato (Addgene #59462) at an MOI of 100,000 for 7 days to fluorescently label neuronal processes.

At DIV20, live-cell imaging was carried out on a spinning-disk confocal microscope (IXplore IX83 SpinSR Super-Resolution System), acquiring four fields of view from predefined positions within the axonal compartment while cells were held at 37 °C and 5% CO₂. On day 21, axotomy was performed by vacuum aspiration confined to the axonal compartment. The same fields were then reimaged 24 and 48 hours after injury to assess regrowth. Regeneration was quantified in ImageJ by measuring the area covered by tdTomato-positive neurites and normalising it to the corresponding pre-axotomy value.

### Animals

C57BL/6J wild-type mice purchased from Charles River, UK were used for all experiments. Mice were housed in individually ventilated cages on a 12-hour dark/light cycle and given access to food and water ad libitum. Experiments were conducted in accordance with the United Kingdom Home Office Legislation (Scientific Procedures Act, 1986) and were approved by the Home Office to be conducted at King’s College London.

Individual mice were used as the experimental unit throughout. Results are reported in line with the Essential 10 ARRIVE 2.0 Guidelines.

### Inflammatory arthritis mouse model

The Antigen Induced arthritis (AIA) model was induced in male and female C57BL/6J mice as previously described (Goodwin et al., 2024). Mice were 11 weeks old at the start of the experiment (females 17.2-20.8g, males 21.8-25.5g starting weight). Measurements of knee thickness were taken using callipers.

### LIF and Fast Blue intraarticular injections

For downstream immunohistochemistry (IHC) analysis, mice were injected with 1.5μl Fast Blue (2%, in sterile H20; Polysciences) intraarticularly into the ipsilateral knee 7 days before LIF/BSA injections. Injections were performed under ketamine (75mg/kg) and medetomidine (1mg/kg) anaesthesia, administered intraperitoneally. Once anaesthetised, the mice were placed on a heating pad and temperature maintained at 37C. After shaving the knee and wiping it with 70% ethanol, the patellar tendon was identified and marked by running a 27 g needle in the gap beneath the patella. A Hamilton syringe with a 30 g needle attached was inserted through the patellar tendon at a depth of 2–2.5 mm to deliver the tracer fast blue into the articular space. After the injection, mice were kept anaesthetised in a heated chamber for 1hr, to prevent leaking of the tracer. Sterile saline (0.9%, 0.2mL) was administered subcutaneously to keep the mice hydrated. Anaesthesia was then reversed with atipamezole (1mg/kg), and the mice were allowed to recover in the heated chamber before being returned to their home cage. LIF (Miltenyi Biotec, 130-095-775) was reconstituted at 0.2 mg/ml in 0.1% BSA. Mice were injected intraarticularly into the ipsilateral knee with 2.5μl LIF (0.5 μg) or 0.1% BSA under brief isofluorane anaesthesia. Treatment was allocated using block randomisation by a second experimenter, splitting treatment groups evenly based on sex, cage and whether the mouse had also been injected with Fast Blue, and group allocations were concealed from the primary experimenter until the end of the study.

### Behavioural assays

For the weight-bearing assay, mice were acclimatised to the apparatus (Linton Incapacitance tester, modified with red film covering over the transparent Plexiglas) three times before the first measurement. Peanut butter was applied on the sloped surface of the chamber during acclimatisation and behaviour assessment, to encourage the mice to maintain correct placement during the assay. For weight-bearing measurements, mice were placed in the apparatus and allowed to settle. Measurements were taken while the mice were sitting still, facing forward with one hind paw placed on each plate. Five measurements were taken per mouse, and an average was calculated per paw per mouse. For the AIA experiment, it was not possible for the experimenter to be blinded as the swelling is clearly visible. The experimenter was blinded to treatment groups for experiments where LIF was injected intraarticularly.

For the PAM test used to assess joint pressure thresholds, mice were acclimatised to handling and scruffing over two separate sessions prior to any measurements. For the measurements, one experimenter scruffed the mice and gently extended the leg, while the other experimenter applied pressure on the knee, aiming for a force of 50gf/s with a maximum total force of 450gf. Pressure was removed and total force applied recorded once the mouse responded to the stimuli with a sudden leg retraction and/or vocalisation. If a mouse did not respond, pressure was removed once the maximum force was reached (450gf) and that number was recorded. Two measurements were taken per mouse for each timepoint and an average response calculated. Both experimenters were blinded to the treatment groups during the PAM test.

This study did not deviate from conventional pre-clinical sample sizes (N=10 per group), with the rationale that AIA, due to its extensive joint swelling, is expected to have extremely large effects on pain perception. N=10 gives an 80% chance to detect a main effect of f=0.52 with a repeated measures ANOVA and an alpha = 0.05. Based on our *in vitro* data, we further expected LIF to have similarly large effects on neuronal hyper-excitability and pain. The final sample size for the LIF study was smaller, due to the following exclusions: two LIF mice had to be excluded from PAM measurements due to small fighting wounds in the scruff area, meaning that a secure scruff could not be maintained without causing unnecessary pain and distress to the mice; one vehicle mouse had to be excluded from the 5-72hr LIF study due to an unexpected health condition that was not related to treatment.

Data were analysed in GraphPad, with vehicle-injected mice used as the control group in a repeated measures ANOVA for LIF injection studies, and contralateral knees used as a control for ipsilateral knees for the AIA study presented in Figure 7. Final n numbers and statistical results are provided in the Figure legends.

### Mouse tissue collection and processing for immunofluorescent staining

The DRGs of mice previously injected with Fast Blue were collected for immunofluorescent staining. Briefly, mice were transcardially perfused under terminal anaesthesia with 10ml PBS followed by 20ml 4% paraformaldehyde (PFA). Each ipsilateral L4 DRG was collected and incubated in 4% PFA for 2 hours. DRGs were then washed two times in PBS and placed in 30% sucrose for three days. After that, they were cut on a cryostat at a thickness of 10μm and mounted on microscope slides.

Slides were allowed to dry at room temperature and incubated in 100% methanol at -20C for 10min to permeabilise them. Slides were washed with PBS followed by a 2-hour incubation in 6% Donkey serum in 0.2%Triton in PBS. DRG slices were stained with NeuN (Merck, ABN91, chicken), CGRP (Enzo Life Sciences, BML-CA1137-0025, sheep) and pSTAT3 (Cell Signalling, #9145, rabbit) antibodies overnight at room temperature. Slides were washed and incubated with donkey antibodies against chicken (488, A78948), sheep (568, A-21099) and rabbit (647, A32795) diluted at 1:1000 for 2 hours.

Slides were washed and mounted with fluoromount. DRGs were imaged on a Zeiss LSM 710 confocal microscope using a 20× objective (20×/0.8na). A 5 μm z-stack at 1 μm intervals was taken for each region imaged and 4-5 regions were imaged per sample, aiming to image 6 Fast Blue positive cells per sample where possible.

### IHC Image analysis

The experimenter was blinded while acquiring and analysing the images. Images were analysed on FIJI as follows: a maximum intensity projection of the Z stack was created and the channels split and converted to binary images. The threshold was set so that the positive signal was clearly visible with as little background as possible. The binary images of pSTAT3, CGRP or Fast Blue were multiplied with the binary image for NeuN to ensure only neurons were included in the analysis. Outliers were removed (setting the radius to 5 in the “remove outlies” function in FIJI) for all resulting images. The resulting pSTAT3 signal was dilated (2x) and holes filled. ROIs were created based on the resulting pSTAT3 binary image, and overlayed on the NeuN, CGRP and Fast Blue images to count pSTAT3+ cells in all three channels. pSTAT3 negative cells were counted manually for each channel. Sections with no Fast Blue positive cells were excluded from the analysis.

Final sample sizes of n=6 meant that only very large effects would have been detectable at reliable rates (80% chance to detect d=1.79 with an independent t-test and alpha = 0.05).

### Mouse tissue collection and processing for LIF ELISA

Mice were transcardially perfused under terminal anaesthesia with 10ml PBS and knees collected. The knees were chopped up with a scalpel and incubated in digestion buffer (312.5μg/ml Liberase and 50μg/ml DNAse in RPMI) at 37C, 200rpm for 1.5 hours. Cells were centrifuged at 400g for 5 minutes, and supernatant collected.

Supernatants were diluted 1 in 10, and a LIF ELISA was carried out as per manufacturer’s instructions (Abcam; ab238261), running all samples in duplicate.

### Mouse tissue collection and processing for Synovial Fibroblast culture

Legs from mice were collected, skin and muscle removed, keeping the joints intact, and legs frozen in Freezing media (High Glucose DMEM +LGlut, supplemented with 20% FBS and 10% DMSO) and stored at -80C. Legs were thawed on ice and incubated in digestion buffer (0.1mg/ml Libearse and 0.1mg/ml DNAse I in RPMI) at 37C, 200rpm for 30min. The supernatant was discarded, and legs rinsed in RPMI. The joints were prized apart and placed into fresh digestion buffer at 37C, 200rpm for 30min. The supernatant was passed through a 100μm cell strainer, followed by an equal volume of media to rinse. Cells were centrifuged at 400g for 5min and the supernatant discarded. Cells were washed with FACS buffer (2mM EDTA, 15mM HEPES and 0.4% BSA in HBSS) and incubated in antibody cocktail (CD146 - BV605, Ter119 - BV785, CD45 - FITC, CD31 - PerCpCy5.5, PDPN - PE, Thy1 - PECy7) + Fc block for 30min on ice. Cells were washed and resuspended in FACS buffer +DAPI (10 ng/ml). Sublining synovial fibroblasts (Ter119-DAPI-CD45-CD31-CD146-PDPN+Thy1+) were sorted using an Aria 2 with a 100μm nozzle into 100% FBS. Cells were centrifuged at 400g for 5min and resuspended in plating media (High glucose DMEM +GlutaMax, supplemented with 15% FBS, 1% P/S, 50μM βMercaproethanol) and plated at 1,000 cells per well in a 96-well plate. Cells were allowed to proliferate for two weeks and treated with IL-1β (Fisher; 211-11B-10UG; 10ng/ml) or vehicle for 24 hours. Conditioned media was collected and diluted 1:2 to be used in the LIF ELISA as recommended by the manufacturers.

### Primary mouse sensory neuron culture and analysis

Glass coverslips coated with 0.1 mg/mL poly-L-lysine (Sigma-Aldrich, #P8954) were plated, washed, and dried overnight in a laminar flow hood. Adult mice were anesthetised and perfused with 10 mL of ice-cold PBS. Following decapitation, a total laminectomy was performed along the vertebral column to expose the spinal cord and dorsal roots. DRGs from cervical to lumbar levels were microdissected bilaterally and collected into Ham’s F12 medium (Sigma-Aldrich, #N6658) on ice. Ganglia were washed with fresh F12 and digested in an enzymatic solution containing 3 mg/mL dispase II (Roche, #4942078001), 0.8 mg/mL collagenase (Sigma-Aldrich, #C9891), and 30 μg/mL DNase I (Roche, #10104159001) at 37°C for 1 h under gentle agitation. DRGs were mechanically dissociated in F12 by sequential trituration through fire-polished glass Pasteur pipettes and pulse-centrifuged (up to 800 rpm), with the resulting supernatant collected. This process was repeated three times, and the supernatant was passed through a 70 µm nylon cell strainer. To remove non-neuronal debris and myelin fragments, the cell suspension was pelleted (1,200 rpm, 10 min), resuspended in F12, and carefully layered on top of a 15% bovine serum albumin (BSA) (Sigma-Aldrich, #A3294) cushion in PBS. Following centrifugation (1,000 rpm, 10 min), the upper debris layer and supernatant were aspirated, and the neuronal pellet was resuspended in complete culture medium (Ham’s F12 supplemented with 10% FBS (Gibco, #10500-064), 1% Antibiotic-Antimycotic (ThermoFisher, #15240096), and 1% N2 supplement (ThermoFisher, 17502048)). 2 hours prior to cell seeding, coverslips were coated with a 10 µL drop of 40 µg/mL mouse laminin (Sigma-Aldrich, #L2020) diluted in F12 at 37°C. Cells were seeded at a density of 5,000 cells per well. Plated neurons were allowed to adhere for 1 h at 37°C in a humidified 5% CO2 incubator before wells were flooded with 500 µL of complete culture medium. After 24 hours, cultures were treated with recombinant mouse LIF (100 ng/mL; Miltenyi Biotec, #130-095-772) or vehicle control for 1 h at 37°C prior to fixation for immunocytochemistry or lysis for Western blot analysis.

For immunocytochemistry, cells were fixed in 2% paraformaldehyde (PFA) for 10 min at 37°C. Cells were permeabilised in 100% methanol at –20°C for 10 min, followed by blocking with 10% donkey serum (Abcam, #ab7475) in PBS containing 0.2% Triton X-100 (Sigma-Aldrich, #T9284) for 1 h at room temperature. Coverslips were then incubated overnight at 4°C with primary antibodies against NeuN (1:750; Merck, #ABN91) and pSTAT3 (1:100; Cell Signalling, #9145) in 10% donkey serum PBS-T. Secondary antibodies, donkey anti-chicken 488 (1:1000; Invitrogen #A78948) and donkey anti-rabbit 647 (1:1000; Invitrogen, #A32795), were applied for 2 h at room temperature. PBS washes were performed between each step. Coverslips were then mounted on glass slides using DAPI-containing mounting medium (Invitrogen, #00-4959-52).

For Western blot, cultured DRG neurons were washed once with PBS, harvested in 4X Laemmli buffer (Bio-Rad, # 1610747), and denatured at 95°C for 5 min. Protein samples and a ladder (Thermo Scientific, #26619) were resolved on Novex Tris-Glycine Mini Protein Gels (Invitrogen, #XP08165BOX) at 100V for 90 min. Separated proteins were transferred onto a methanol-activated 0.45 μm PVDF membrane (Thermo Scientific, #10617354) at 100V for 60 min. For detection of pSTAT3, the membrane was incubated with the anti-pSTAT3 primary antibody (1:2000; Cell Signalling, #9145) diluted in Western blotting accelerator (Proteintech, #PR20039) for 20 min at room temperature, washed 3 times with Tris-buffered saline containing 0.1% Tween-20 (TBS-T) (Sigma-Aldrich, #P9416), and incubated with an HRP-conjugated secondary antibody (1:5000; Abcam, #ab205718) for 15 min at room temperature. After washing, blots were visualised using the ECL Plus HRP Substrate Kit (Abbkine, #K22030) and captured on an iBright Imaging System (Invitrogen). To detect total STAT3, the membrane was stripped using ReBlot Plus Strong Antibody Stripping Solution (Millipore, #2504) for 15 min, washed, and then restained with anti-STAT3 primary antibody (1:1000; Cell Signalling, 4904).

## Supporting information

Cytokines RNA-seq R script

Fibroblast CM RNA-seq R script

Synovial Fluid RNA-seq R script

## Acknowledgements

We thank Trishalee Gungoosingh for her help with numerous media changes throughout the differentiation process.

We acknowledge King’s College London as the source of HPSI0714i-kute_4 human iPSC line, which was generated under the Human Induced Pluripotent Stem Cell Initiative funded by a grant from the Wellcome Trust and Medical Research Council, supported by the Wellcome Trust (WT098051) and the NIHR/Wellcome Trust Clinical Research Facility, and acknowledge Life Science Technologies Corporation as the provider of CytoTune. This research was supported by Research and Development, Guy’s and St Thomas’ NHS Foundation Trust. The views expressed are those of the author(s) and not necessarily those of the NHS.

By using the AD2 iPSC line, the research leading to these results has received support from the Innovative Medicines Initiative Joint Undertaking under grant agreement n° 115439, resources of which are composed of financial contribution from the European Union’s Seventh Framework Programme (FP7/2007-2013) and EFPIA companies’ in-kind contribution.

The UKBi013-A-GCamP6f iPSC line was generated as part of an Innovative Medicines Initiative 2 Joint Undertaking under Grant Agreement no. 116072. This Joint Undertaking has received the support from the European Union’s Horizon 2020 research and innovation.

This research was supported by the Advanced Cytometry Platform (Flow Core) Research and Development, Guy’s and St Thomas’ NHS Foundation Trust. The views expressed are those of the authors and not necessarily those of the NHS

## Authors contributions

RR and YL conducted the majority of the *in vitro* iPSC-derived neuron experiments, supported by IZ, IM, NA and KF. RR and FD performed the RNA-seq analysis. JVW performed all the mouse experiments, supported by IZ, RC, GG, ACM & TT. MAA performed experiments involving the HD 10.6 DRG line, supported by RR and supervised by DB. ZRL, FB, SR, FH and BK were involved in recruitment, consenting, collecting and/or processing of clinical samples. FD and LST conceptualised and co-supervised the project, and together with RR and JVW drafted the manuscript. All authors provided intellectual support and advice and edited and/or reviewed the final draft of the manuscript.

