## Supplementary Figures for "LIF is a pro-algesic mediator in human and mouse models of arthritis-related pain"

### Supplementary Figure 1

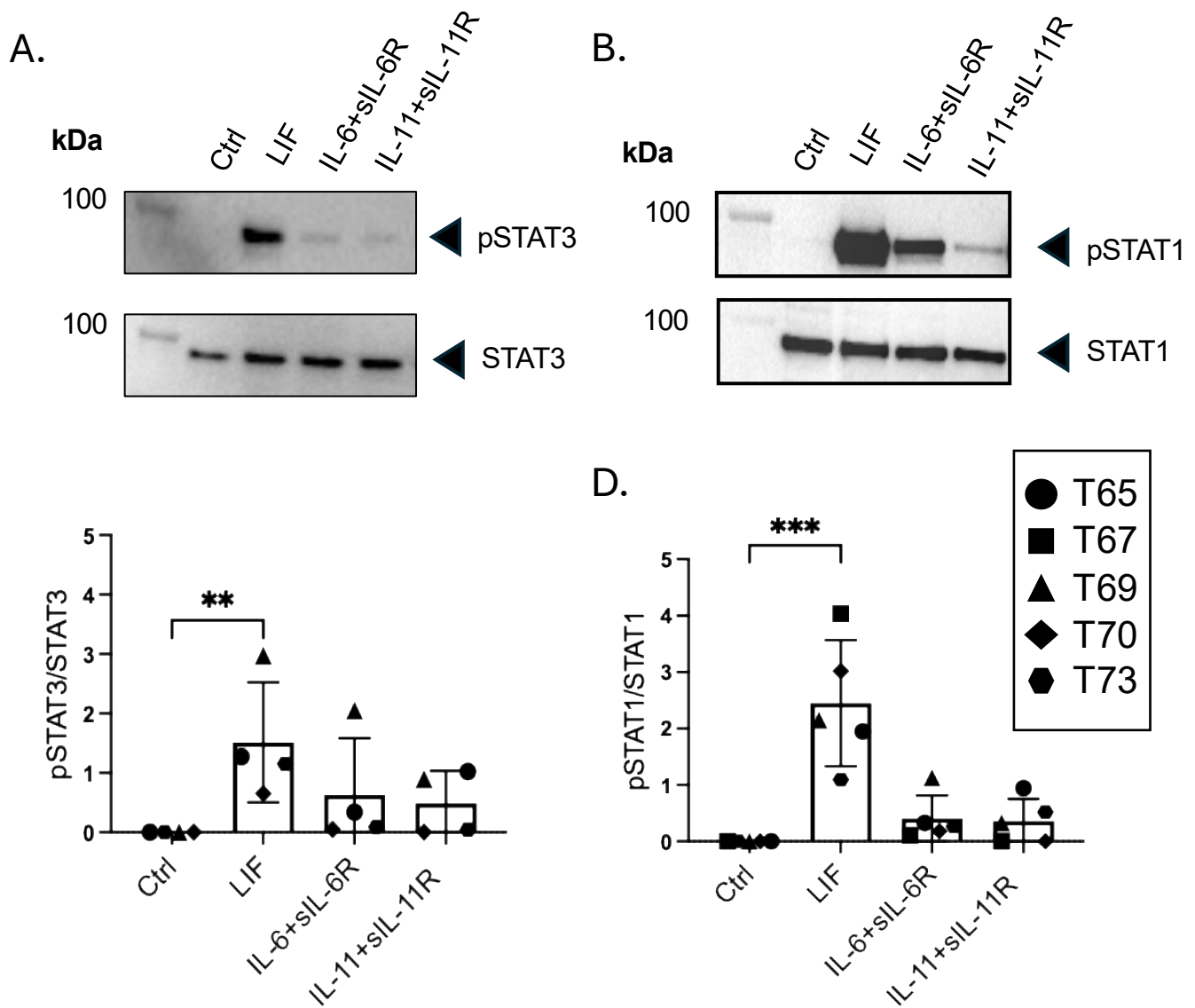

**Supplementary Figure 1: LIF induces higher levels of pSTAT3 and pSTAT1 in sensory neurons than IL-6 and IL-11.** A, B. Representative Western blots of pSTAT3 (A) and pSTAT1 (B) in iPSC-derived sensory neurons (generated from the Kute 4 iPSC line) after 30 minutes incubation with LIF (10 ng/ml), IL-6+sIL-6R (100 ng/ml) or IL-11+sIL-11R (100 ng/ml) with corresponding densitometry shown in C,D, n=5 different iPSC-derived neuronal differentiations (T65-T73). Statistical analysis was performed using a Kruskal Wallis test followed by Dunn's multiple comparison test comparing cytokine means to the control (Ctrl) mean, *padj* = 0.006, *padj* = 0.005.

#### Supplementary Figure 2

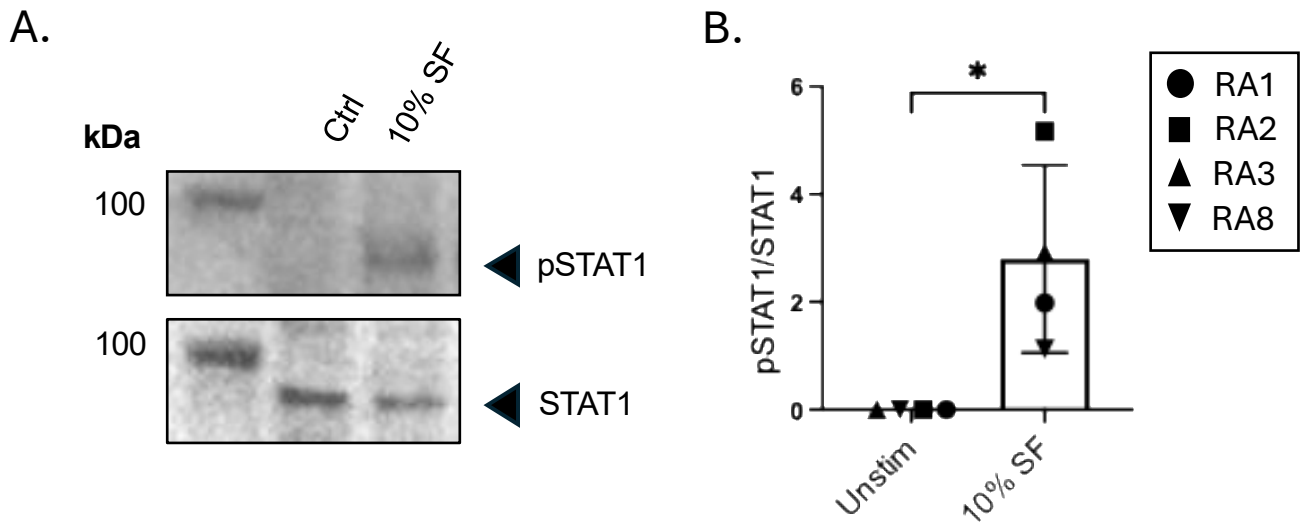

**Supplementary Figure 2: RA SF induces neuronal pSTAT1.** A. Representative Western blot showing pSTAT1 in iPSC-derived sensory neurons incubated for 1 hour with 10% RA SF. B. Corresponding densitometry of pSTAT1/STAT1 in sensory neurons from $n=1$ neuronal differentiation across $n=1$ iPSC line (K4) upon treatment with $n=4$ different RA SF donor samples. Statistical analysis was performed using a Mann-Whitney U test, $p = 0.0286$.

### Supplementary Figure 3

A.

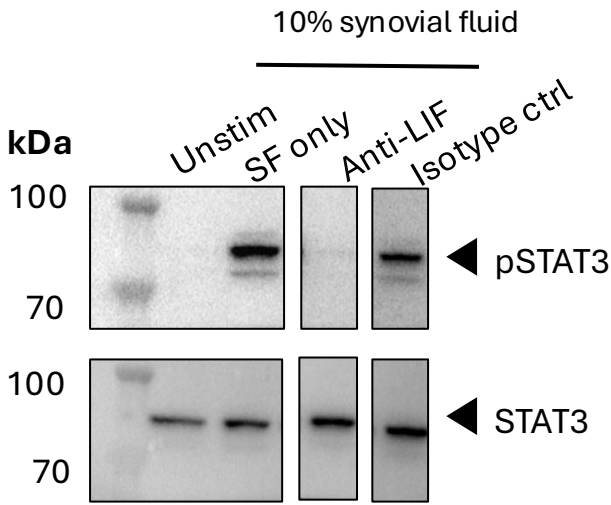

B.

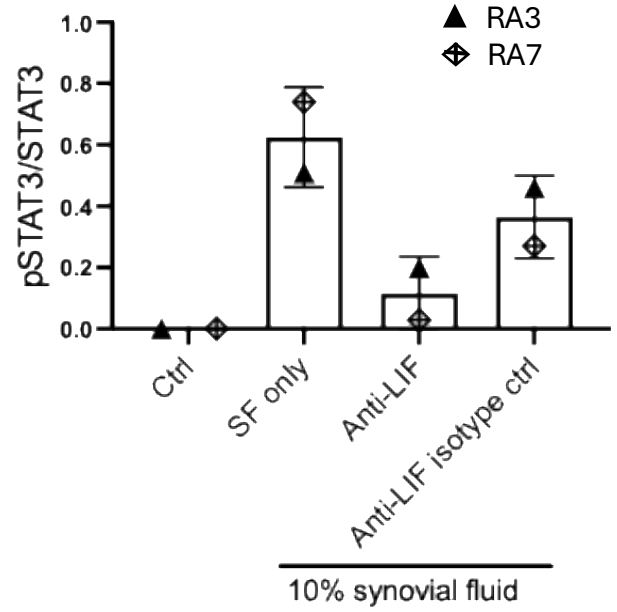

**Supplementary Figure 3: Effect of anti-LIF vs. isotype control Ab on RA SF-induced neuronal pSTAT3.** A. Representative example of a Western blot showing pSTAT3 levels in iPSC-derived sensory neurons after a 1hr incubation with 10% v/v RA SF alone, anti-LIF (4 μg/ml) or anti-LIF isotype control (4 μg/ml). B. Corresponding densitometry of n=1 neuronal differentiation in n=1 iPSC line (AD2) exposed to n=2 RA SF.

### Supplementary Figure 4

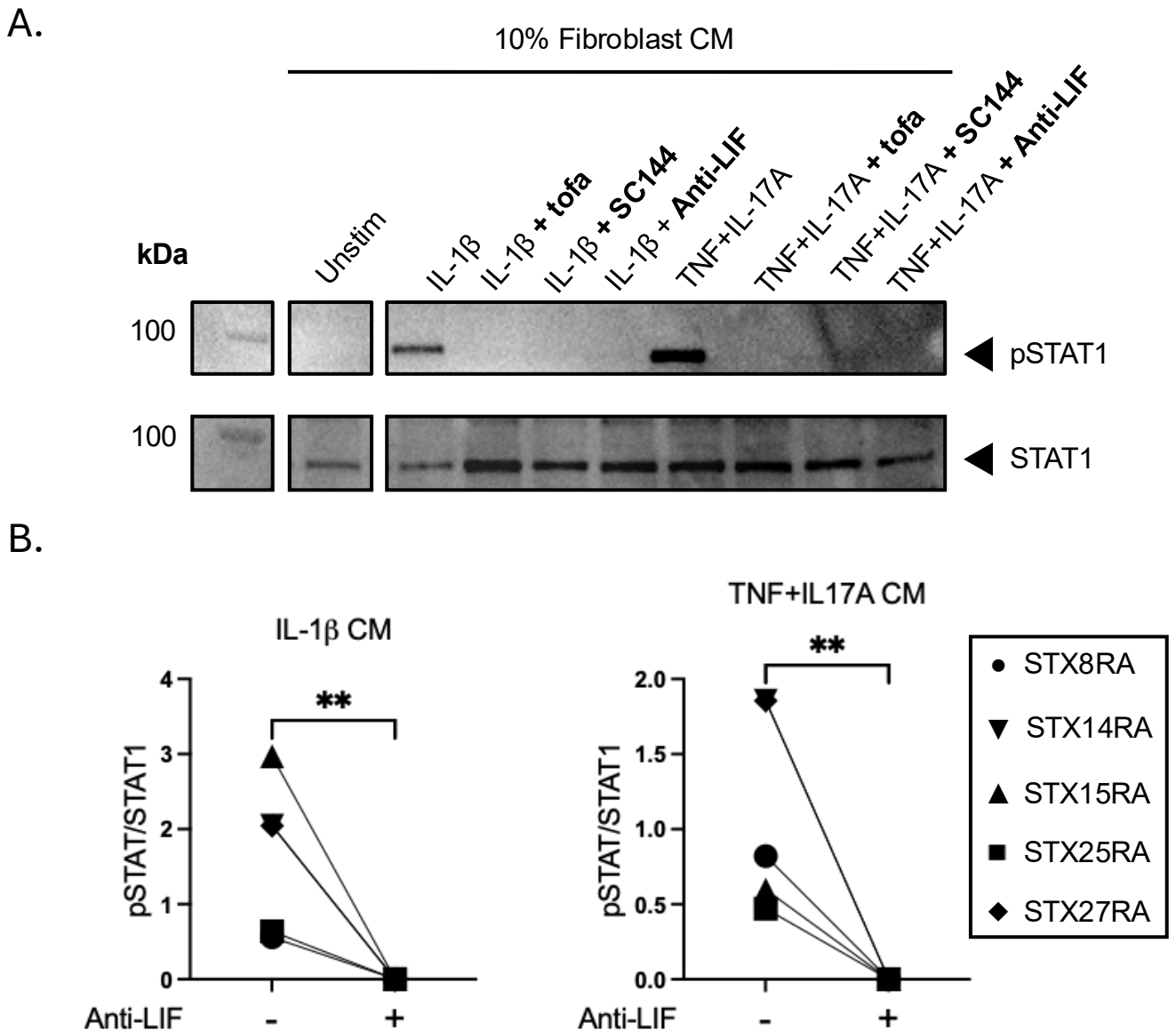

**Supplementary Figure 4: RA fibroblast CM induces neuronal pSTAT1 in a LIF-dependent manner.** A. Representative Western blot showing pSTAT1 in iPSC-derived sensory neurons after 1hr exposure to 10% v/v conditioned media from unstimulated or cytokine treated fibroblasts, in the absence or presence of tofacitinib (2 μM), the gp130 inhibitor SC144 (20 μM) or anti-LIF (4 μg/ml). B. Corresponding densitometry analysis of n=3 different neuronal differentiations across n=1 iPSC line, exposed to n=5 RA synovial fibroblast CM. Statistical analysis was performed using a Mann-Whitney U test, $p = 0.0079$, $p = 0.0079$.

### Supplementary Figure 5

A.

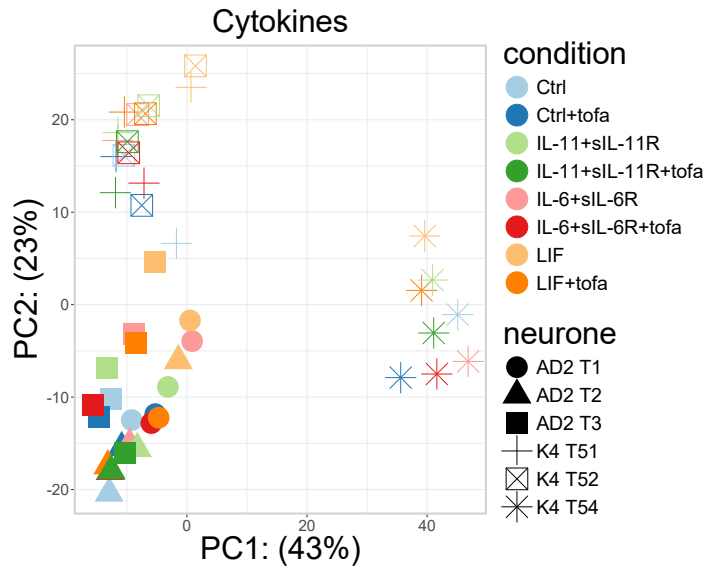

B.

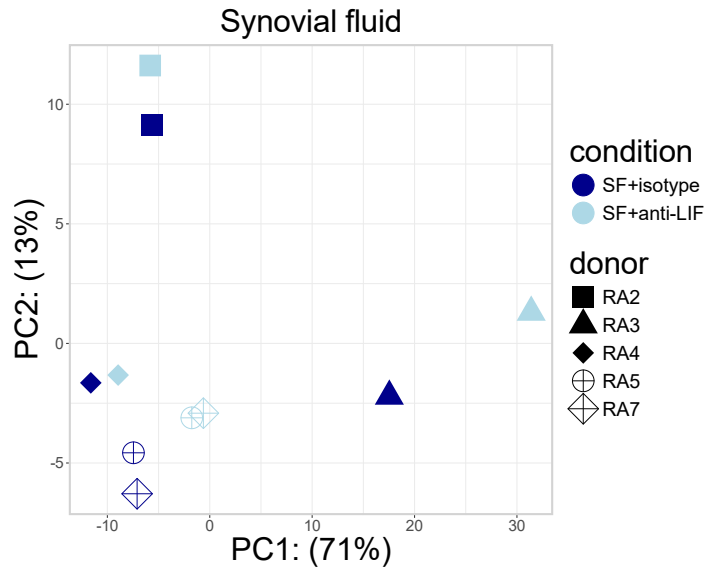

C.

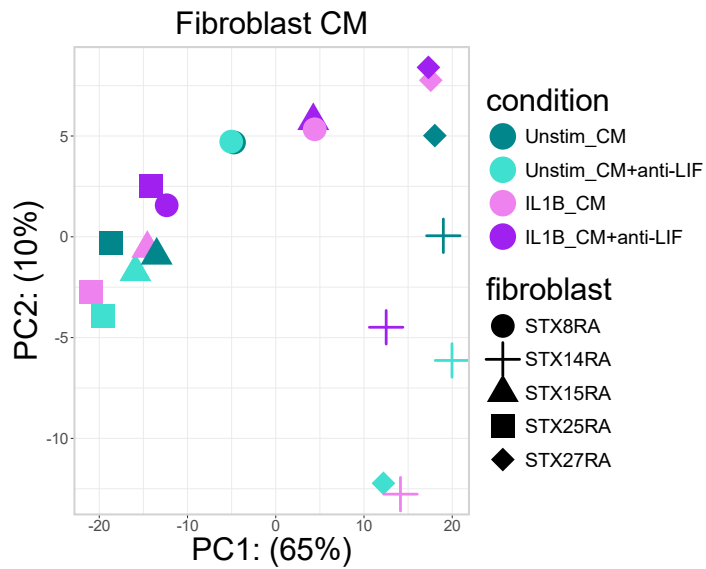

**Supplementary Figure 5: PCA plots for RNA-sequencing of iPSC-derived sensory neurons.** PCA plots showing transcriptional variation in sensory neurons after 24 hr exposure to: (A) media, IL-11 + sIL-11R, IL-6 + sIL-6R or LIF (all at 100ng/ml) in the absence or presence of tofacitinib (2 μM), (B) 10% v/v RA SF with anti-LIF (4 μg/ml) or isotype control (4 μg/ml), or (C) 10% v/v RA synovial fibroblast CM with or without anti-LIF. Different symbols indicate n=3 different neuronal differentiations each (T1-T54) from two different iPSC lines (AD2 and K4, A), n=5 different SF donors (RA2-7, B) and CM from n=5 different donor-derived fibroblast lines (STX8RA-27RA, C).

### Supplementary Figure 6

A.

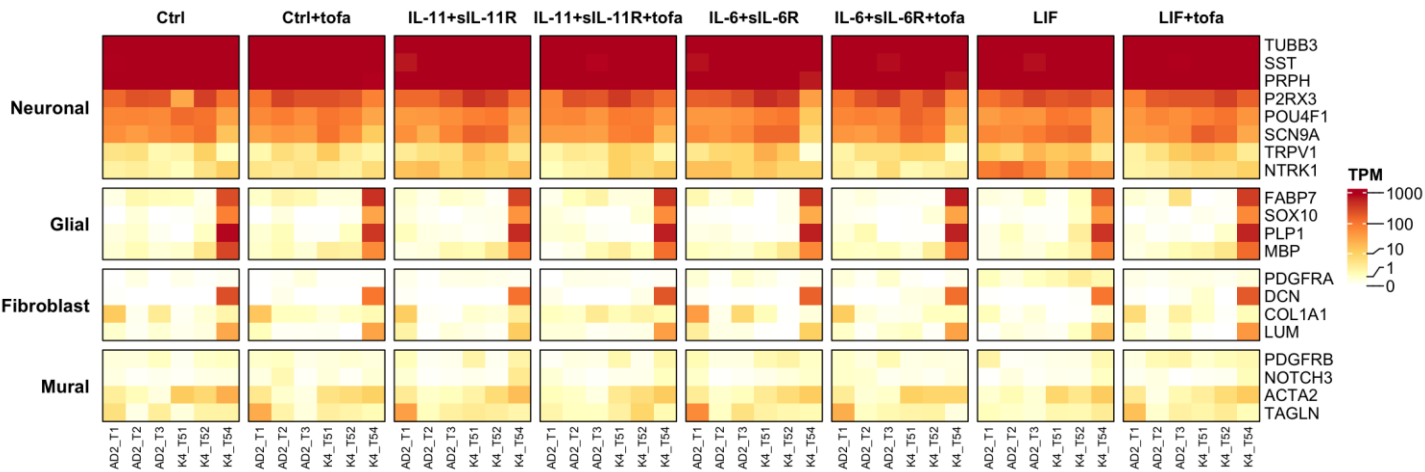

B.

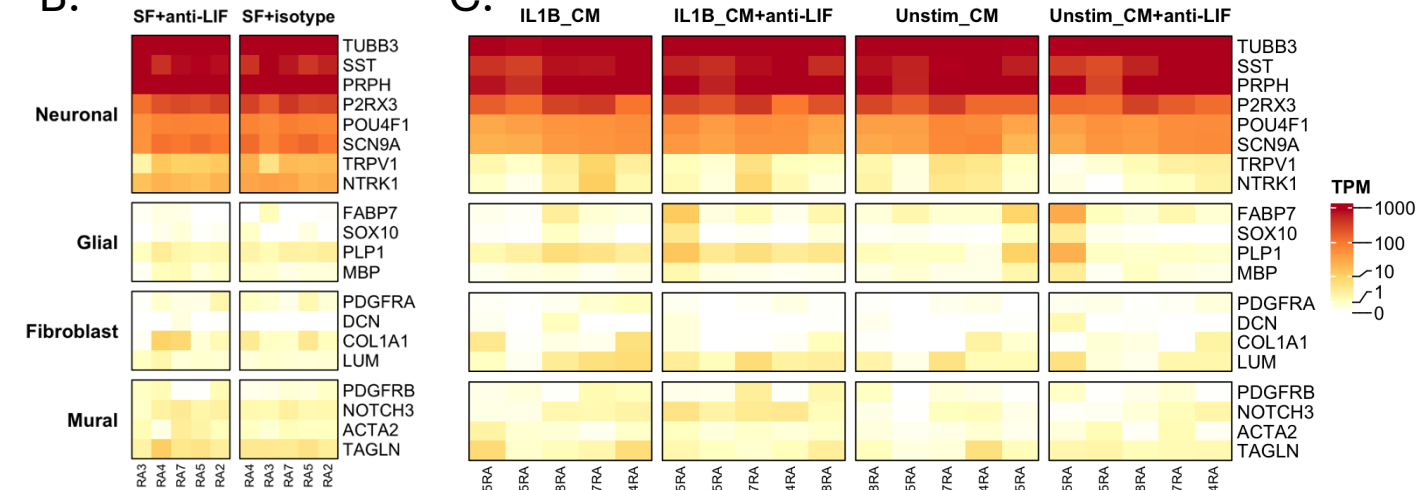

C.

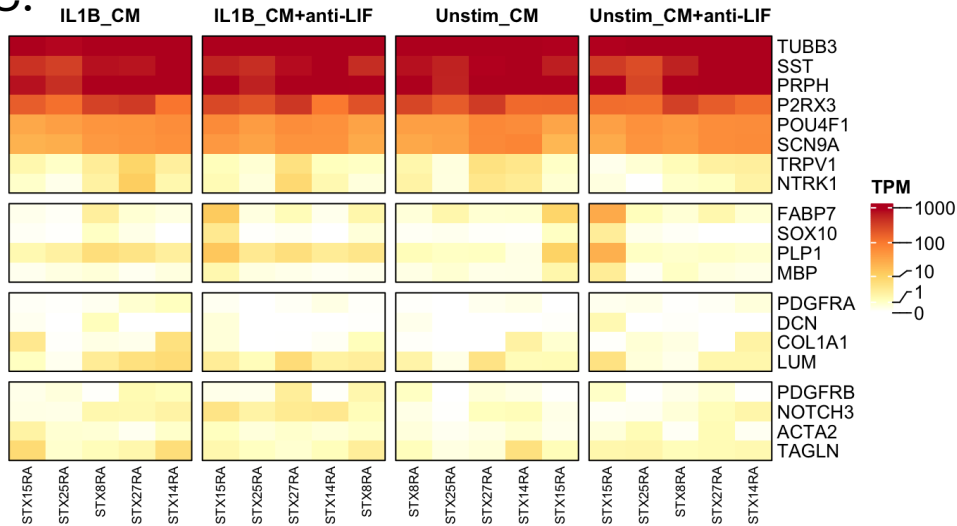

**Supplementary Figure 6: Heatmap of marker genes demonstrates expected expression of neuronal genes and negligible levels of non-neuronal gene expression.** Three different experiments using two different iPSC lines each were conducted where iPSC-derived sensory neurons were treated with cytokines (A), synovial fluids (SF, B), or conditioned media (CM) from fibroblasts (C). Blocks indicate n=3 different neuronal differentiations each (T1-T54) from two different iPSC lines (AD2 and K4, A), n=5 different SF donors (RA2-7, B) and CM from n=5 different donor-derived fibroblast lines (STX8RA-27RA, C). RNA extractions proceeded by experiment in batch-controlled fashion, while library preparation and sequencing were conducted on all samples at once. Neuronal gene expression was in keeping with previous literature, e.g. with high levels of the sensory neuron specific transcription factor, BRN3A (POU4F1). Negligible levels of non-neuronal gene expression were observed in all differentiation but one (K4\_T54).

### Supplementary Figure 7

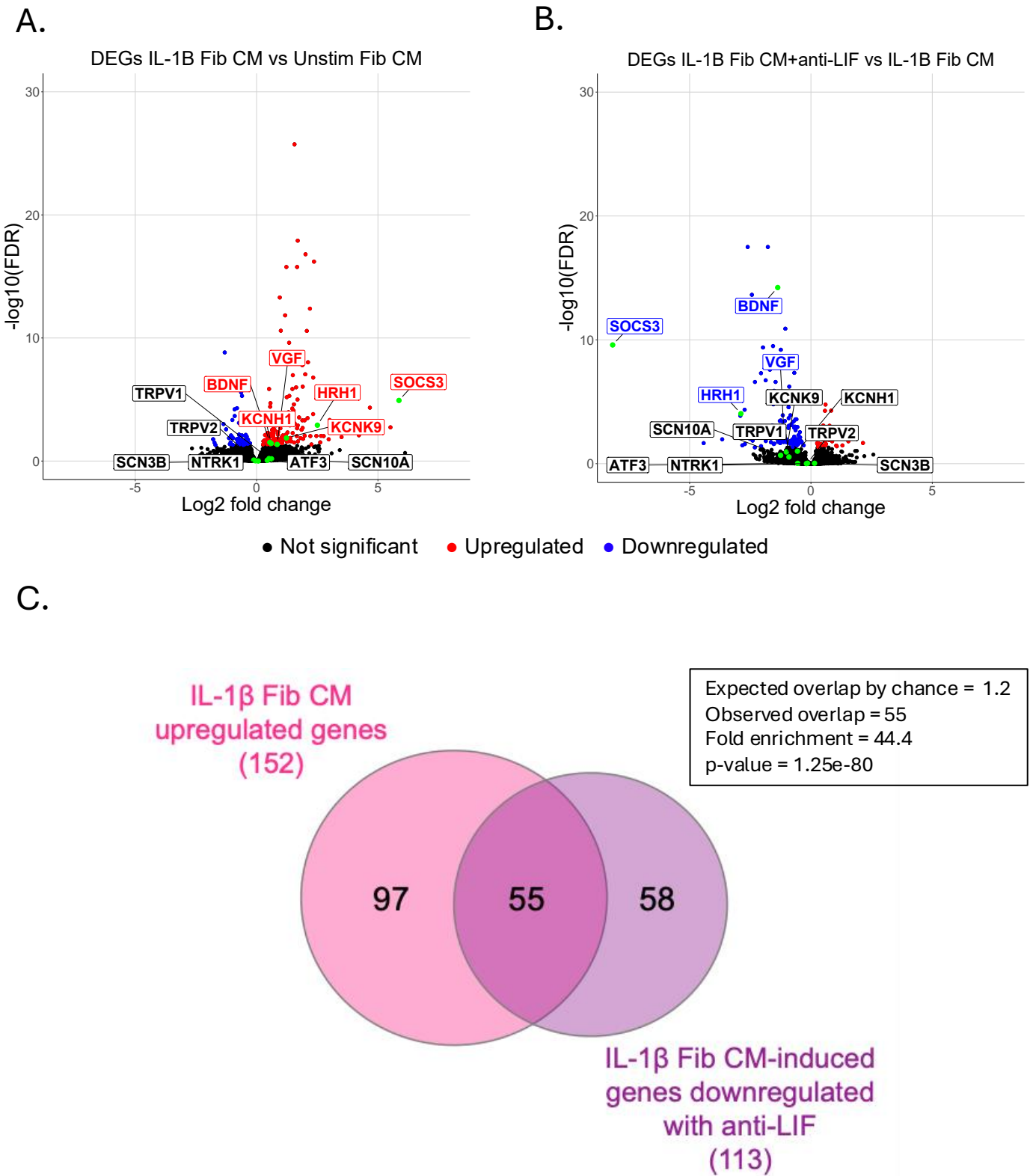

**Supplementary Figure 7: Pairwise comparisons to determine neuronal genes induced by IL-1β-treated fibroblast CM but downregulated with anti-LIF.** A. Volcano plots showing differentially expressed genes from pairwise comparison of A. IL-1β-treated fibroblast CM vs unstimulated fibroblast CM (significant DEGs FDR ≤ 0.05, Wald test) or B. IL-1β-treated fibroblast CM with anti-LIF vs IL-1β-treated fibroblast CM alone (significant DEGs FDR ≤ 0.05, Wald test). Hyperexcitability and/or peripheral sensitisation-associated genes are highlighted with a green dot and either blue (downregulated), red (upregulated), or black (non-significant) text. C. Venn diagram showing the overlap between IL-1β-treated fibroblast CM upregulated genes and IL-1β-treated fibroblast CM-induced genes downregulated with anti-LIF. Statistical analysis was performed using a Fisher's exact test.

### Supplementary Figure 8

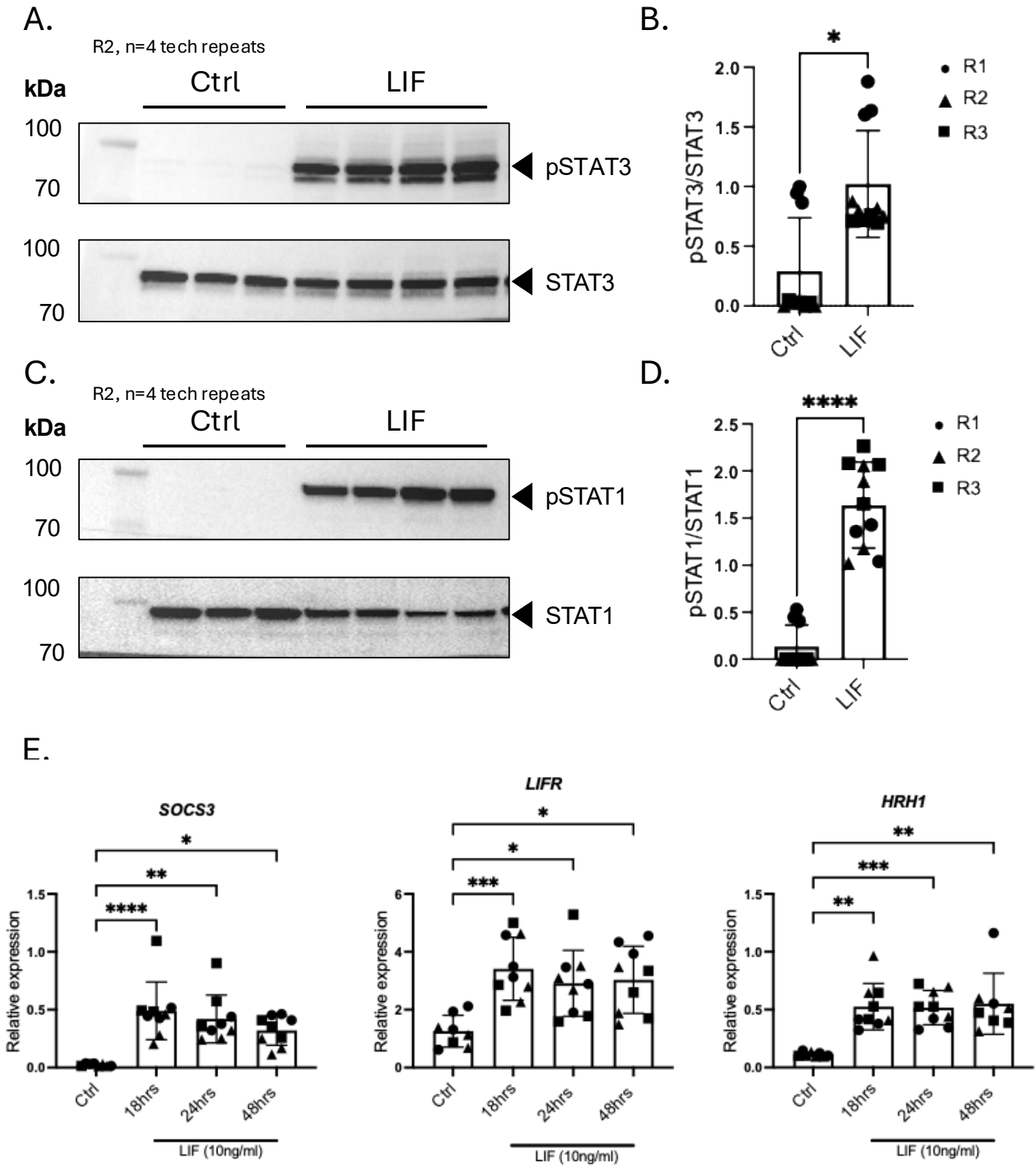

**Supplementary Figure 8: HD10.6 neurons induce neuronal pSTAT3 and pSTAT1 in response to LIF.** A-D. Representative Western blots showing pSTAT3 (A) and pSTAT1 (C) in HD10.6 neurons after 30-minute incubation with LIF (10ng/ml) and corresponding densitometry (B, D) of n=3 neuronal differentiations with at least 3 technical repeats of each condition. Statistical analysis was performed using a Mann-Whitney U test, $p = 0.024$, $p = <0.0001$. E. *SOC3S*, *LIFR* and *HRH1* expression in HD10.6 neurons after 24 hrs incubation with LIF (10ng/ml). All cells were starved of all growth factors for 8 hours prior to stimulation. Statistical analysis was performed using a Kruskal Wallis test followed by Dunn's multiple comparison test comparing timepoint means to the control (Ctrl) mean, Ctrl vs. 18hrs $padj = <0.0001$, $padj = 0.0008$, $padj = 0.0019$, Ctrl vs. 24hrs $padj = 0.0018$, $padj = 0.0171$, $padj = 0.0006$, Ctrl vs. 48hrs $padj = 0.0143$, $padj = 0.0121$, $padj = 0.0028$.

### Supplementary Figure 9

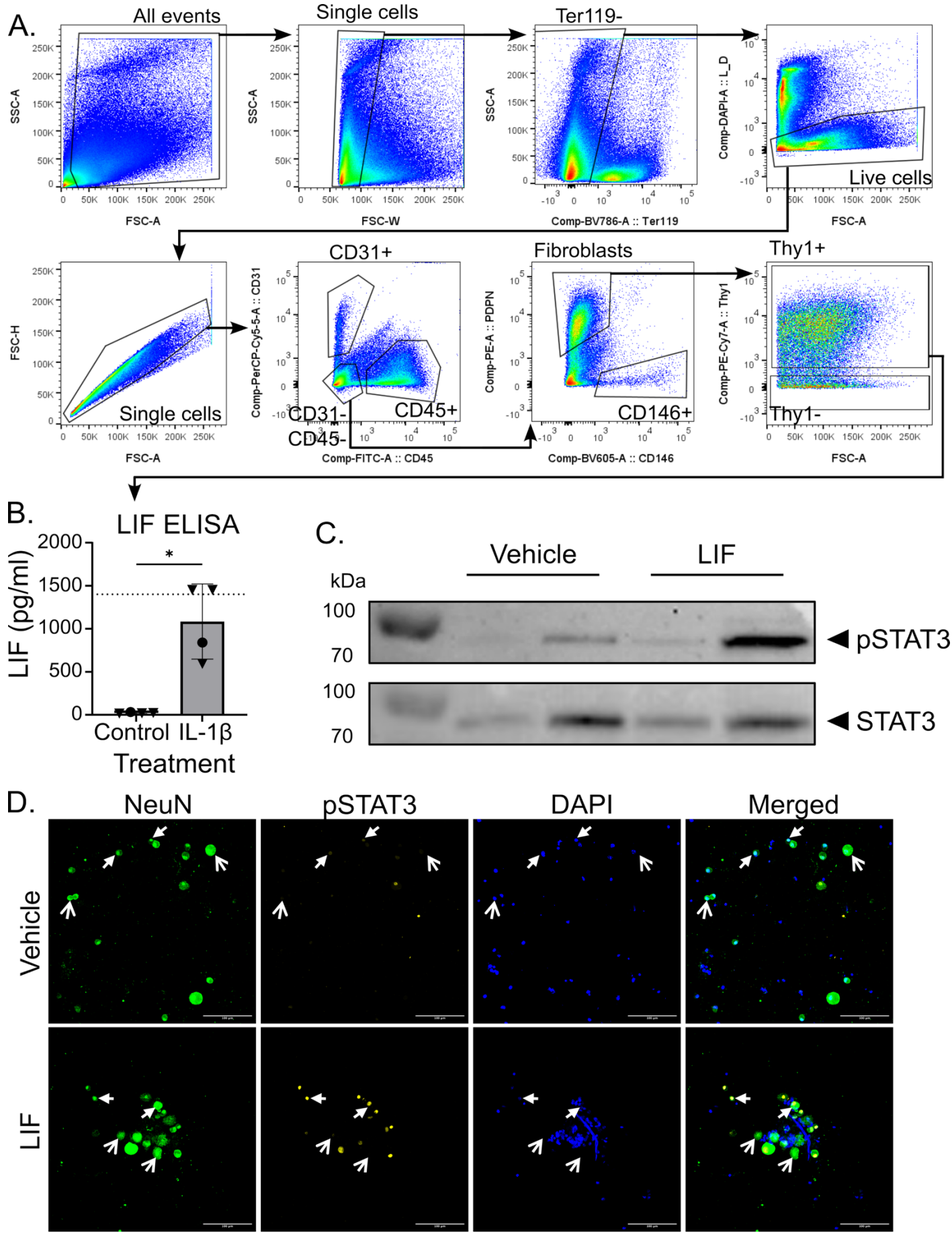

#### Supplementary Figure 9

##### **Supplementary Figure 9: Stimulated mouse fibroblasts release LIF, and primary mouse sensory neurons respond to LIF by pSTAT3 induction, mirroring results from human cellular models.**

A. Gating strategy used to isolate synovial sublining fibroblasts from mouse knees using FACS, for tissue culture. B. LIF measured in the supernatant of cultured synovial sublining fibroblasts treated with 10 ng/ml IL-1β or vehicle for 24 hours, measured using an ELISA. N=4. Each symbol represents a well of primary fibroblasts derived from a different mouse; Welch's t-test was used; error bars show SD. circles = males, triangles = females. C. Western blot for pSTAT3 and total STAT3 from adult mouse primary sensory neurons, grown for 24 hours and subsequently incubated in recombinant mouse LIF (100 ng/mL) or vehicle control 1 hour prior to protein extraction. Results from two individual mice are shown. D. Representative immunocytochemistry images of cultured adult mouse primary sensory neurons treated as in C. Cells were immunostained for pSTAT3 (Tyr705), the neuronal marker NeuN, and DAPI to visualise nuclei. Individual and merged channels are shown. Closed arrows indicate pSTAT3-positive neurons; open arrows indicate pSTAT3-negative neurons. Scale bar: 100 μm.

### Supplementary Figure 10

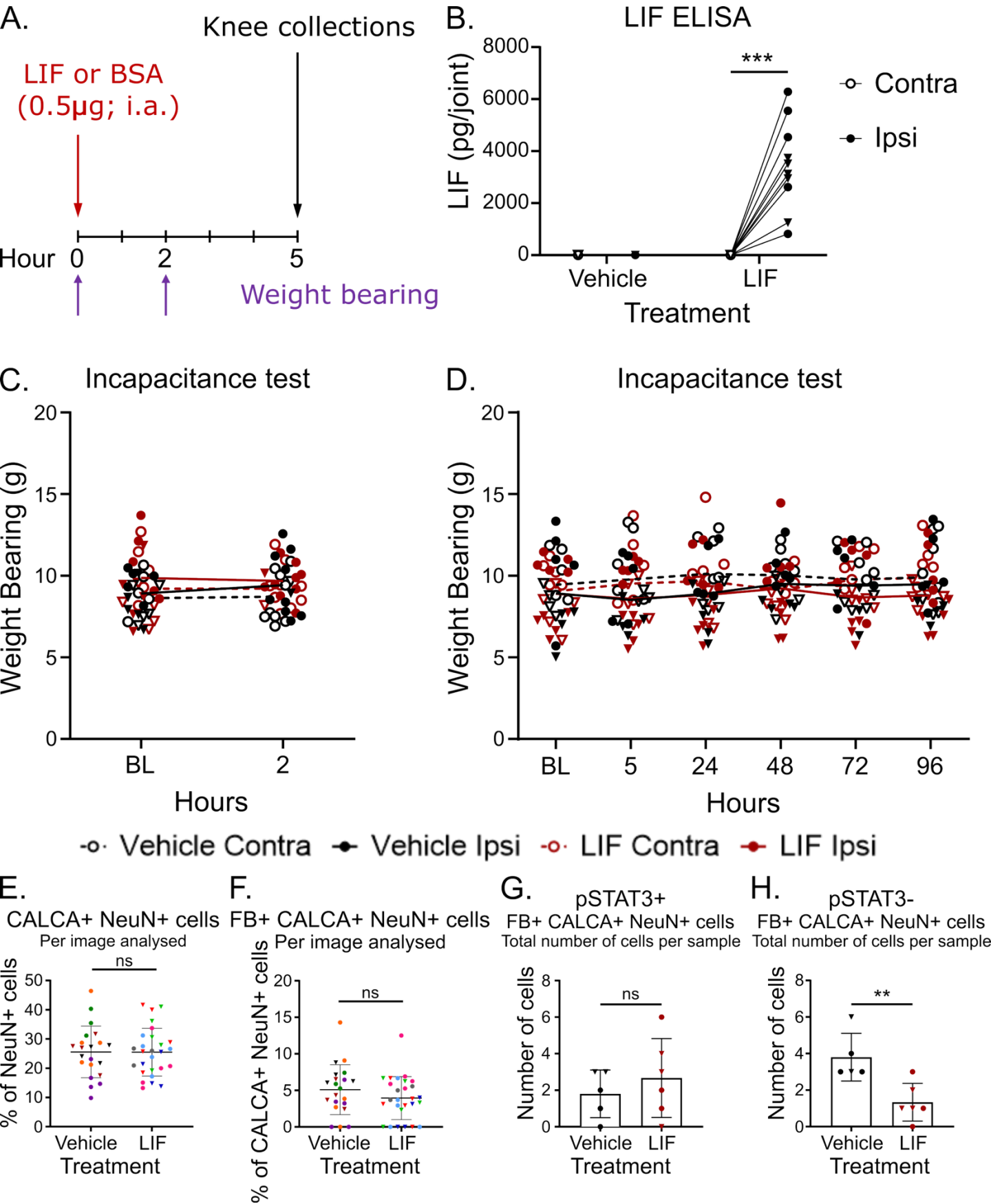

#### Supplementary Figure 10

**Supplementary Figure 10: Intraarticular LIF injection has no effect on weight bearing in mice.** A. Schematic representation of experimental plan: LIF (0.5µg) or vehicle (0.1% BSA) was injected in the ipsilateral knee of wildtype mice, and an incapacitance test was carried out to assess weight bearing at baseline (BL) and 2 hours after LIF injection. Knees were collected 5 hours after injection for downstream processing. B. LIF measured in the supernatant of digested mouse knees collected 5 hours after intraarticular (i.a.) LIF injection, measured using an ELISA. n=10. Each dot is an individual mouse. 2-way ANOVA with uncorrected Fisher's LSD (Treatment: \*\*\*, Knee: \*\*\*, Treatment x Knee: \*\*\*). C. Weight placed on each leg in mice injected with LIF or vehicle at 2 hours. n=10. Each dot is an individual mouse. D. Weight bearing results for experiment presented in Fig. 8 of the main manuscript. Vehicle: n=9, LIF: n=10. Each dot is an individual mouse. E. Percentage of CALCA+ neurons among NeuN neurons in the L4 DRG of mice 4 days after intraarticular injection with LIF or vehicle. Each symbol represents a section (n=21 vehicle and n=27 LIF sections from n=5 vehicle and n=6 LIF mice). Stats were applied to the mean value per mouse. F. Percentage of FastBlue+ neurons among CALCA and NeuN double positive cells per image analysed. Each symbol represents one image, each colour represents one mouse. Each dot represents an image analysed (n=21 vehicle and n=27 LIF sections from n=5 vehicle and n=6 LIF mice). Stats were applied to the mean value per mouse. G-H. Total number of FastBlue, CALCA and NeuN triple positive cells per mouse, that were also pSTAT3 positive (G) or negative (H). Each symbol represents an individual mouse. Vehicle: n=5, LIF: n=6. Welch's test; error bars show SD. Circles = males, Triangles = females.
