## Supplementary material for "LIF is a pro-algesic mediator in human and mouse models of arthritis-related pain": Cytokines RNA-seq R script

LIF\_paper\_cytokines


### LIF\_paper\_cytokines

###### Rosie Ross

#### 2026-08-21

```
knitr::opts_knit$set(
  root.dir = "/Users/franziskadenk/Library/CloudStorage/Dropbox/WTCA/kallisto_output/kallisto_for_paper")

knitr::opts_chunk$set(
  echo = TRUE,
  warning = FALSE,
  message = FALSE)

library(DESeq2)
```

```
## Loading required package: S4Vectors
```

```
## Loading required package: stats4
```

```
## Loading required package: BiocGenerics
```

```
## 
## Attaching package: 'BiocGenerics'
```

```
## The following objects are masked from 'package:stats':
## 
##     IQR, mad, sd, var, xtabs
```

```
## The following objects are masked from 'package:base':
## 
##     anyDuplicated, aperm, append, as.data.frame, basename, cbind,
##     colnames, dirname, do.call, duplicated, eval, evalq, Filter, Find,
##     get, grep, grepl, intersect, is.unsorted, lapply, Map, mapply,
##     match, mget, order, paste, pmax, pmax.int, pmin, pmin.int,
##     Position, rank, rbind, Reduce, rownames, sapply, saveRDS, setdiff,
##     table, tapply, union, unique, unsplit, which.max, which.min
```

```
## 
## Attaching package: 'S4Vectors'
```

```
## The following object is masked from 'package:utils':
## 
##     findMatches
```

```
## The following objects are masked from 'package:base':
## 
##     expand.grid, I, unname
```

```
## Loading required package: IRanges
```

```
## Loading required package: GenomicRanges
```

```
## Loading required package: GenomeInfoDb
```

```
## Loading required package: SummarizedExperiment
```

```
## Loading required package: MatrixGenerics
```

```
## Loading required package: matrixStats
```

```
## 
## Attaching package: 'MatrixGenerics'
```

```
## The following objects are masked from 'package:matrixStats':
## 
##     colAlls, colAnyNAs, colAnys, colAvgsPerRowSet, colCollapse,
##     colCounts, colCummaxs, colCummins, colCumprods, colCumsums,
##     colDiffs, colIQRDiffs, colIQRs, colLogSumExps, colMadDiffs,
##     colMads, colMaxs, colMeans2, colMedians, colMins, colOrderStats,
##     colProds, colQuantiles, colRanges, colRanks, colSdDiffs, colSds,
##     colSums2, colTabulates, colVarDiffs, colVars, colWeightedMads,
##     colWeightedMeans, colWeightedMedians, colWeightedSds,
##     colWeightedVars, rowAlls, rowAnyNAs, rowAnys, rowAvgsPerColSet,
##     rowCollapse, rowCounts, rowCummaxs, rowCummins, rowCumprods,
##     rowCumsums, rowDiffs, rowIQRDiffs, rowIQRs, rowLogSumExps,
##     rowMadDiffs, rowMads, rowMaxs, rowMeans2, rowMedians, rowMins,
##     rowOrderStats, rowProds, rowQuantiles, rowRanges, rowRanks,
##     rowSdDiffs, rowSds, rowSums2, rowTabulates, rowVarDiffs, rowVars,
##     rowWeightedMads, rowWeightedMeans, rowWeightedMedians,
##     rowWeightedSds, rowWeightedVars
```

```
## Loading required package: Biobase
```

```
## Welcome to Bioconductor
## 
##     Vignettes contain introductory material; view with
##     'browseVignettes()'. To cite Bioconductor, see
##     'citation("Biobase")', and for packages 'citation("pkgname")'.
```

```
## 
## Attaching package: 'Biobase'
```

```
## The following object is masked from 'package:MatrixGenerics':
## 
##     rowMedians
```

```
## The following objects are masked from 'package:matrixStats':
## 
##     anyMissing, rowMedians
```

```
library(ggplot2)
```

```
## Warning: package 'ggplot2' was built under R version 4.4.3
```

```
library(org.Hs.eg.db)
```

```
## Loading required package: AnnotationDbi
```

```
##
```

```
library(AnnotationDbi)
library(devEMF)
library(ggrepel)
```

```
## Warning: package 'ggrepel' was built under R version 4.4.3
```

```
library(tidyverse)
```

```
## ── Attaching core tidyverse packages ──────────────────────── tidyverse 2.0.0 ──
## ✔ dplyr     1.1.4     ✔ readr     2.1.5
## ✔ forcats   1.0.0     ✔ stringr   1.5.1
## ✔ lubridate 1.9.4     ✔ tibble    3.2.1
## ✔ purrr     1.0.2     ✔ tidyr     1.3.1
```

```
## ── Conflicts ────────────────────────────────────────── tidyverse_conflicts() ──
## ✖ lubridate::%within%() masks IRanges::%within%()
## ✖ dplyr::collapse()     masks IRanges::collapse()
## ✖ dplyr::combine()      masks Biobase::combine(), BiocGenerics::combine()
## ✖ dplyr::count()        masks matrixStats::count()
## ✖ dplyr::desc()         masks IRanges::desc()
## ✖ tidyr::expand()       masks S4Vectors::expand()
## ✖ dplyr::filter()       masks stats::filter()
## ✖ dplyr::first()        masks S4Vectors::first()
## ✖ dplyr::lag()          masks stats::lag()
## ✖ ggplot2::Position()   masks BiocGenerics::Position(), base::Position()
## ✖ purrr::reduce()       masks GenomicRanges::reduce(), IRanges::reduce()
## ✖ dplyr::rename()       masks S4Vectors::rename()
## ✖ lubridate::second()   masks S4Vectors::second()
## ✖ lubridate::second<-() masks S4Vectors::second<-()
## ✖ dplyr::select()       masks AnnotationDbi::select()
## ✖ dplyr::slice()        masks IRanges::slice()
## ℹ Use the conflicted package (<http://conflicted.r-lib.org/>) to force all conflicts to become errors
```

```
library(reshape2)
```

```
## 
## Attaching package: 'reshape2'
## 
## The following object is masked from 'package:tidyr':
## 
##     smiths
```

```
library(clusterProfiler)
```

```
## 
## clusterProfiler v4.14.6 Learn more at https://yulab-smu.top/contribution-knowledge-mining/
## 
## Please cite:
## 
## T Wu, E Hu, S Xu, M Chen, P Guo, Z Dai, T Feng, L Zhou, W Tang, L Zhan,
## X Fu, S Liu, X Bo, and G Yu. clusterProfiler 4.0: A universal
## enrichment tool for interpreting omics data. The Innovation. 2021,
## 2(3):100141
## 
## Attaching package: 'clusterProfiler'
## 
## The following object is masked from 'package:purrr':
## 
##     simplify
## 
## The following object is masked from 'package:AnnotationDbi':
## 
##     select
## 
## The following object is masked from 'package:IRanges':
## 
##     slice
## 
## The following object is masked from 'package:S4Vectors':
## 
##     rename
## 
## The following object is masked from 'package:stats':
## 
##     filter
```

```
library(biomaRt)
library(dplyr)
library(purrr)
library(tximport)
library(grid) 
library(enrichplot)
```

```
## enrichplot v1.26.6 Learn more at https://yulab-smu.top/contribution-knowledge-mining/
## 
## Please cite:
## 
## Guangchuang Yu, Fei Li, Yide Qin, Xiaochen Bo, Yibo Wu and Shengqi
## Wang. GOSemSim: an R package for measuring semantic similarity among GO
## terms and gene products. Bioinformatics. 2010, 26(7):976-978
```

```
library(ComplexHeatmap)
```

```
## ========================================
## ComplexHeatmap version 2.22.0
## Bioconductor page: http://bioconductor.org/packages/ComplexHeatmap/
## Github page: https://github.com/jokergoo/ComplexHeatmap
## Documentation: http://jokergoo.github.io/ComplexHeatmap-reference
## 
## If you use it in published research, please cite either one:
## - Gu, Z. Complex Heatmap Visualization. iMeta 2022.
## - Gu, Z. Complex heatmaps reveal patterns and correlations in multidimensional 
##     genomic data. Bioinformatics 2016.
## 
## 
## The new InteractiveComplexHeatmap package can directly export static 
## complex heatmaps into an interactive Shiny app with zero effort. Have a try!
## 
## This message can be suppressed by:
##   suppressPackageStartupMessages(library(ComplexHeatmap))
## ========================================
```

```
library(SuperExactTest)
```

```
## 
## Attaching package: 'SuperExactTest'
## 
## The following objects are masked from 'package:lubridate':
## 
##     intersect, union
## 
## The following objects are masked from 'package:dplyr':
## 
##     intersect, union
## 
## The following objects are masked from 'package:GenomicRanges':
## 
##     intersect, union
## 
## The following object is masked from 'package:GenomeInfoDb':
## 
##     intersect
## 
## The following objects are masked from 'package:IRanges':
## 
##     intersect, union
## 
## The following objects are masked from 'package:S4Vectors':
## 
##     intersect, union
## 
## The following objects are masked from 'package:BiocGenerics':
## 
##     intersect, union
## 
## The following objects are masked from 'package:base':
## 
##     intersect, union
```

```
theme_set(theme_gray(base_family = "Arial"))
```

Uploading and processing aligned files (abundance.tsv files from
Kallisto)

```
base_dir = "/Users/franziskadenk/Library/CloudStorage/Dropbox/WTCA/kallisto_output/kallisto_for_paper"

# List all subdirectories 
folders = list.dirs(base_dir, recursive = FALSE, full.names = TRUE)

# Filter for directories starting with AD2_, T51_, T52_, T53_, or T54_
target_folders = folders[basename(folders) %>% str_detect("^(AD2_|T51_|T52_|T53_|T54_)")]

# Construct full paths to abundance.tsv files
files = file.path(target_folders, "abundance.tsv")
names(files) = basename(target_folders)

# Filter out any that don't exist (just in case)
files = files[file.exists(files)]

# Double-check
all(file.exists(files))  # Should be TRUE
```

```
## [1] TRUE
```

```
# Connect to Ensembl
mart <- useEnsembl(biomart = "genes", dataset = "hsapiens_gene_ensembl", host = "https://may2025.archive.ensembl.org")

# Use one abundance.tsv file to get transcript IDs
sample_file = read_tsv(files[1])
ensembl_ids = sample_file$target_id

# Remove version suffix (e.g., ENS...1.2 -> ENS...1)
ensembl_ids_clean = sub("\\..*", "", ensembl_ids)

# for gene symbols 
tx2gene = getBM(attributes = c("ensembl_transcript_id", "external_gene_name"),
                filters = "ensembl_transcript_id",
                values = ensembl_ids_clean,
                mart = mart)

txi <- tximport(files,
                type = "kallisto",
                tx2gene = tx2gene,
                ignoreTxVersion = TRUE)

# Save txi
saveRDS(txi, file = "txi_cytokine.rds")
write.csv(txi$abundance, file = "txi_cytokines_TPM.csv")
write.csv(txi$counts, file = "txi_cytokines_counts.csv")
```

```
# read in sample sheet 
sample_sheet_cyto = read.table("sample_sheet.txt", header = TRUE, row.names = 1, sep="\t")
sample_sheet_cyto = sample_sheet_cyto[colnames(txi$counts), ]

# sample names in sample_sheet need to match colnames in txi$counts including the order
all(colnames(txi$counts) == rownames(sample_sheet_cyto)) # Should return TRUE
```

```
## [1] TRUE
```

```
#makes sure the group information is a factor
sample_sheet_cyto$condition = factor(sample_sheet_cyto$condition)
sample_sheet_cyto$neurone = factor(sample_sheet_cyto$neurone)

# run DESeq from Tximport 
dds = DESeqDataSetFromTximport(txi,
                               colData = sample_sheet_cyto,
                               design = ~ neurone + condition)
```

```
library(circlize)
# ============================================================
# 1. Marker genes
# ============================================================

marker_groups <- list(
  Neuronal = c(
    "TUBB3", "SST", "PRPH", "P2RX3", "POU4F1",
    "SCN9A", "TRPV1", "NTRK1"
  ),
  
  Glial = c(
    "FABP7", "SOX10", "PLP1", "MBP"
  ),
  
  Fibroblast = c(
    "PDGFRA", "DCN", "COL1A1", "LUM"
  ),
  
  Mural = c(
    "PDGFRB", "NOTCH3", "ACTA2", "TAGLN"
  )
)

marker_genes <- unlist(marker_groups, use.names = FALSE)


# ============================================================
# 2. TPM matrix
# ============================================================

tpm <- txi$abundance

# Make sure all dds samples are present in txi
stopifnot(all(colnames(dds) %in% colnames(tpm)))

# Put TPM columns in same order as dds
tpm <- tpm[, colnames(dds), drop = FALSE]


# ============================================================
# 3. Check marker genes
# ============================================================

marker_present <- marker_genes[
  marker_genes %in% rownames(tpm)
]

marker_missing <- setdiff(
  marker_genes,
  rownames(tpm)
)

cat("Missing marker genes:\n")
```

```
## Missing marker genes:
```

```
print(marker_missing)
```

```
## character(0)
```

```
# ============================================================
# 4. Subset to marker genes
# ============================================================

marker_tpm <- tpm[
  marker_present,
  ,
  drop = FALSE
]

# ============================================================
# 5. Log-transform TPM for display
# ============================================================

marker_mat <- log2(marker_tpm + 1)

# ============================================================
# 6. Define gene groups
# ============================================================

gene_group <- rep(
  names(marker_groups),
  lengths(marker_groups)
)

names(gene_group) <- unlist(
  marker_groups,
  use.names = FALSE
)

gene_group <- gene_group[
  rownames(marker_mat)
]

gene_group <- factor(
  gene_group,
  levels = c(
    "Neuronal",
    "Glial",
    "Fibroblast",
    "Mural"
  )
)


# ============================================================
# 7. Condition information from dds
# ============================================================

condition <- factor(dds$condition)

# ============================================================
# 8. Order samples by condition
# ============================================================

column_order <- order(condition)

marker_mat <- marker_mat[
  ,
  column_order,
  drop = FALSE
]

condition <- condition[
  column_order
]

# ============================================================
# 9. Define TPM legend breakpoints
# ============================================================

tpm_breaks <- c(
  0,
  1,
  10,
  100,
  1000
)

log_breaks <- log2(tpm_breaks + 1)

# ============================================================
# 10. Colour scale
# ============================================================

max_log <- max(marker_mat, na.rm = TRUE)

col_fun <- colorRamp2(
  log_breaks,
  c(
    "white",
    "#FFFFCC",
    "#FED976",
    "#FD8D3C",
    "#BD0026"
  )
)

# ============================================================
# 11. Heatmap
# ============================================================

ht <- Heatmap(
  marker_mat,
  
  name = "TPM",
  
  col = col_fun,
  
  # Legend is positioned according to log2(TPM+1),
  # but labelled in actual TPM units
  heatmap_legend_param = list(
    at = log_breaks,
    labels = tpm_breaks,
    title = "TPM"
  ),
  
  
  # ----------------------------------------------------------
  # ROWS
  # ----------------------------------------------------------
  
  row_split = gene_group,
  
  cluster_rows = FALSE,
  
  show_row_names = TRUE,
  
  row_names_gp = gpar(
    fontsize = 10
  ),
  
  row_title_gp = gpar(
    fontsize = 11,
    fontface = "bold"
  ),
  
  row_title_rot = 0,
  
  
  # ----------------------------------------------------------
  # COLUMNS
  # ----------------------------------------------------------
  
  column_split = condition,
  
  cluster_columns = FALSE,
  
  cluster_column_slices = FALSE,
  
  column_labels = as.character(dds$neurone[column_order]),
show_column_names = TRUE,
  
  column_names_gp = gpar(
    fontsize = 7
  ),
  
  column_names_rot = 90,
  
  column_title_gp = gpar(
    fontsize = 10,
    fontface = "bold"
  ),
  
  
  # ----------------------------------------------------------
  # FORMATTING
  # ----------------------------------------------------------
  
  border = TRUE,
  
  column_gap = unit(
    2,
    "mm"
  ),
  
  row_gap = unit(
    2,
    "mm"
  )
)


draw(ht)
```

```
pdf(
  "marker_heatmap.pdf",
  width = 7,
  height = 4
)

draw(ht)

dev.off()
```

```
## quartz_off_screen 
##                 2
```

DESeq analysis - generating a model accounting for iPSC line and
condition

```
#filter counts >=10 counts in >=6 samples in at least 1 condition

raw_counts = counts(dds)
by_cond = split(seq_len(ncol(dds)), colData(dds)$condition)

min_count = 10
req_n = 6

pass_by_cond = lapply(by_cond, function(idx) {
  rowSums(raw_counts[, idx, drop = FALSE] >= min_count) >= req_n
})

keep_any = Reduce(`|`, pass_by_cond)

cat("Kept", sum(keep_any), "genes out of", nrow(dds),
    "that have ≥", min_count, "counts in", req_n,
    "samples in at least one Condition\n")
```

```
## Kept 14210 genes out of 25796 that have ≥ 10 counts in 6 samples in at least one Condition
```

```
dds = dds[keep_any, ]

# save dds as RDS files
saveRDS(dds, file = "dds_cytokine.rds")
dds_wald = DESeq(dds)
saveRDS(dds_wald, file = "dds_wald_cytokine.rds")
```

Creating PCA plot

```
# creating PCA plot (uses top 500 most variable genes)
pca_data = plotPCA(vst(dds), intgroup = c("neurone", "condition"), returnData = TRUE)
percentVar = round(100*attr(pca_data, "percentVar"))

# Set the factor levels to define legend order
pca_data$condition = factor(pca_data$condition,
                            levels = c("Ctrl", "Ctrl+tofa","IL-11+sIL-11R", "IL-11+sIL-11R+tofa", 
                                       "IL-6+sIL-6R", "IL-6+sIL-6R+tofa", "LIF", "LIF+tofa"))
# plot PCA
pca_plot = ggplot(pca_data, aes(x = PC1, y = PC2, colour = condition, shape = neurone)) +
  geom_point(size = 10)+ 
  scale_colour_brewer(palette = "Paired")+
  guides (
    colour = guide_legend(order = 1),
    shape = guide_legend(order =2))+
  labs(title = "Cytokines", 
       x = paste0("PC1: (", percentVar[1], "%)"),
       y = paste0("PC2: (", percentVar[2], "%)"))+
  theme(
    plot.title = element_text(hjust = 0.5, size = 32), 
    axis.title = element_text(size = 32),
    axis.text  = element_text(size = 16),
    panel.background = element_rect(fill = "white"),
    panel.grid = element_line(colour = "grey92"), 
    panel.border = element_rect(colour = "lightgrey", fill = NA, linewidth = 2),
    legend.title = element_text(size = 32),
    legend.text  = element_text(size = 20),
    legend.key.size = unit(1, "cm"))

emf("PCA_plot.emf", height=8, width=10) 
print(pca_plot)
dev.off()
```

```
## quartz_off_screen 
##                 2
```

```
pca_plot
```

Generating and saving useful files including: 1. Abundance (TPM) 2.
Raw counts 3. vst count (vst = variance stabilising transformation) 4.
Normalised counts (size factor, accounting for seq\_depth)

```
# Extracting abundance 
abundance = txi$abundance

# Ensures every row had a unique name 
rownames(abundance) = ifelse(rownames(abundance) == "" | is.na(rownames(abundance)),
                             paste0("Unknown_", seq_along(rownames(abundance))),
                             rownames(abundance))

# Re-order sample sheet and abundance by condition
order_index = order(sample_sheet_cyto$condition)
sample_sheet_cyto = sample_sheet_cyto[order_index,]
abundance = abundance[,order_index]

# generate filtered tpm file >=1 in all samples of at least 1 condition 

by_cond = split(seq_len(ncol(abundance)), sample_sheet_cyto$condition)

min_samples = 5   # require 5 samples
min_tpm = 1       # TPM threshold

pass_by_cond = lapply(by_cond, function(idx) {
  rowSums(abundance[, idx, drop = FALSE] >= min_tpm) >= min_samples
})

keep_any = Reduce(`|`, pass_by_cond)

abundance_filtered = abundance[keep_any, , drop = FALSE]

abundance_out_fil = as.data.frame(abundance_filtered) %>%
  rownames_to_column(var = "gene")

stopifnot(all(rownames(sample_sheet_cyto) == colnames(abundance_filtered)))

# Save filtered abundance as txt file
write.table(abundance_out_fil, file="abundance_cytokines_filtered.txt", sep = "\t", row.names = FALSE, quote = FALSE)

# Saving raw counts
# Extracting raw counts 
raw_counts = txi$counts
# Ensures every row had a unique name 
rownames(raw_counts) = ifelse(rownames(raw_counts) == "" | is.na(rownames(raw_counts)),
                              paste0("Unknown_", seq_along(rownames(raw_counts))),
                              rownames(raw_counts))

# Re-order raw_counts by condition
raw_counts = raw_counts[,order_index]

# Add gene column for row names for saving 
raw_counts_out = as.data.frame(raw_counts) %>%
  rownames_to_column(var = "gene")
stopifnot(all(rownames(sample_sheet_cyto) == colnames(raw_counts)))

# Save raw counts as txt file
write.table(raw_counts_out, file="raw_counts_cytokines.txt", sep = "\t", row.names = FALSE, quote = FALSE)

# Saving vst counts 
# Extracting vst counts 
NT_counts = vst(dds)

# Ensures every row had a unique name 
rownames(NT_counts) = ifelse(rownames(NT_counts) == "" | is.na(rownames(NT_counts)),
                             paste0("Unknown_", seq_along(rownames(NT_counts))),
                             rownames(NT_counts))
# Extract numeric matix 
NT_counts = assay(NT_counts)

# Re-order NT_counts by condition
NT_counts = NT_counts[,order_index]

# Add gene column for row names for saving 
NT_counts_out = as.data.frame(NT_counts) %>%
  rownames_to_column(var = "gene")
stopifnot(all(rownames(sample_sheet_cyto) == colnames(NT_counts)))

# Save NT_counts as txt file
write.table(NT_counts_out, file="NT_counts_cytokines.txt", sep = "\t", row.names = FALSE, quote = FALSE)

# Saving normalised counts
# Extract the normalised counts
norm_counts = as.data.frame(counts(dds_wald, normalized=TRUE)) # using dds_wald here as need size factors for the normalisation

# Ensures every row had a unique name 
rownames(norm_counts) <- ifelse(rownames(norm_counts) == "" | is.na(rownames(norm_counts)),
                                paste0("Unknown_", seq_along(rownames(norm_counts))),
                                rownames(norm_counts))

#round the data (makes it easier to look at)
norm_counts = round(norm_counts,2)

# Re-order by condition
norm_counts = norm_counts[,order_index]

# Add gene column for row names for saving 
norm_counts_out = as.data.frame(norm_counts) %>%
  rownames_to_column(var = "gene")
stopifnot(all(rownames(sample_sheet_cyto) == colnames(norm_counts)))

# Save norm_counts as txt file
write.table(norm_counts_out, file="norm_counts_cytokines.txt", sep = "\t", row.names = FALSE, quote = FALSE)
```

Pairwise comparison - 3 comparisons 1. LIF vs Ctrl 2. IL-11 vs Ctrl
3. IL-6 vs Ctrl Generates volcano plot highlighting genes of
interest

```
# 1. LIF vs Ctrl 
de = results(dds_wald, c("condition","LIF","Ctrl")) 

# save to disk
de_out = as.data.frame(de)

# Ensures every row had a unique name 
rownames(de_out) = ifelse(rownames(de_out) == "" | is.na(rownames(de_out)),
                             paste0("Unknown_", seq_along(rownames(de_out))),
                             rownames(de_out))

# Add gene column for row names for saving
de_out = de_out %>% rownames_to_column(var = "gene")

# merge with TPM (abundance)
de_fil = merge(de_out, abundance_out_fil, by = "gene")

#add log10 p value
de_fil$mlog10p = -log10(de_fil$padj)

#order master by padj
sorted_order = order(de_fil[,"padj"], decreasing=FALSE) 
de_fila = de_fil[sorted_order,]

# Save results 
write.table(de_fil, file='lif_ctrl_filtered.txt', row.names = FALSE, quote = FALSE, sep = "\t")


# subsetting data based on significance (norm & transformed counts)
de_fil_siga = subset(de_fil, padj < 0.05)
de_fil_sig_upa = subset(de_fil, padj < 0.05 & (log2FoldChange) >0)
de_fil_sig_downa = subset(de_fil, padj < 0.05 & (log2FoldChange) < 0)
write.table(de_fil_siga, file="sig_lif_ctrl.txt", row.names=FALSE, quote = FALSE, sep = "\t")
write.table(de_fil_sig_downa, file="sig_down_lif_ctrl.txt", row.names=FALSE, quote = FALSE, sep = "\t")
write.table(de_fil_sig_upa, file="sig_up_lif_ctrl.txt", row.names=FALSE, quote = FALSE, sep = "\t")

# Genes you want to highlight/label
genes_interest <- c(
  "TRPV1", "TRPV2", "NTRK1", "HRH1", "VGF", "BDNF",
  "KCNH1", "KCNK9", "SCN10A", "SCN3B", "ATF3", "SOCS3"
)

# Subset DE results
de_interest <- de_fila[de_fila$gene %in% genes_interest, ]

labels <- de_interest

labels$label_colour <- ifelse(
  labels$padj < 0.05 & labels$log2FoldChange > 0, "red",
  ifelse(
    labels$padj < 0.05 & labels$log2FoldChange < 0,
    "blue",
    "black"
  )
)

# Creating volcano plot 
vol_plota <- ggplot(de_fila, aes(x = log2FoldChange, y = mlog10p)) +
  
  # All genes
  geom_point(colour = "black", size = 2) +
  
  # Significant upregulated
  geom_point(
    data = de_fil_sig_upa,
    colour = "red",
    size = 2
  ) +
  
  # Significant downregulated
  geom_point(
    data = de_fil_sig_downa,
    colour = "blue",
    size = 2
  ) +
  
  # Highlight genes of interest
  geom_point(
    data = de_interest,
    colour = "green",
    size = 3
  ) +
  
  # Labels
  geom_label_repel(
    data = labels,
    aes(label = gene),
    colour = labels$label_colour,
    size = 8,
    fontface = "bold",
    force = 30,
    force_pull = 0.5,
    max.overlaps = Inf,
    box.padding = 1,
    point.padding = 1,
    segment.colour = "black",
    show.legend = FALSE
  ) +
  
  labs(
    title = "DEGs LIF vs Ctrl", x="Log2 fold change",
    x = "Log2 fold change",
    y = "-log10(FDR)"
  ) +
  
  coord_cartesian(
    xlim = c(-10, 10),
    ylim = c(0, 80),
    clip = "off"
  ) +
  
  theme(axis.title.x=element_text(size = 30),
        plot.title = element_text(size = 30, hjust = 0.5), 
        axis.title.y = element_text(size = 30),
        axis.text.x = element_text(size = 20),
        axis.text.y = element_text(size = 20),
        axis.line.x = element_line(colour = "black"),
        axis.line.y = element_line(colour = "black"),
        panel.background = element_blank(),
        panel.grid.major = element_line(colour = "lightgrey"))

emf("Vol_plot_lif_ctrl_genesinterest.emf", height=10, width=10) 
print(vol_plota)
dev.off()
```

```
## quartz_off_screen 
##                 2
```

```
#2. IL-11 vs Ctrl 
de = results(dds_wald, c("condition","IL-11+sIL-11R","Ctrl")) 

# save to disk
de_out = as.data.frame(de)

# Ensures every row had a unique name 
rownames(de_out) = ifelse(rownames(de_out) == "" | is.na(rownames(de_out)),
                          paste0("Unknown_", seq_along(rownames(de_out))),
                          rownames(de_out))

# Add gene column for row names for saving
de_out = de_out %>% rownames_to_column(var = "gene")


# merge with TPM (abundance)
de_fil = merge(de_out, abundance_out_fil, by = "gene")

#add log10 p value
de_fil$mlog10p = -log10(de_fil$padj)

#order master by padj
sorted_order = order(de_fil[,"padj"], decreasing=FALSE) 
de_filb = de_fil[sorted_order,]

write.table(de_filb, file='il11_ctrl_filtered.txt', row.names = FALSE, quote = FALSE, sep = "\t")

# subsetting data based on significance (norm & transformed counts)
de_fil_sigb = subset(de_filb, padj < 0.05)
de_fil_sig_upb = subset(de_filb, padj < 0.05 & (log2FoldChange) >0)
de_fil_sig_downb = subset(de_filb, padj < 0.05 & (log2FoldChange) < 0)
write.table(de_fil_sigb, file="sig_il11_ctrl.txt", row.names=FALSE, quote = FALSE, sep = "\t")
write.table(de_fil_sig_downb, file="sig_down_il11_ctrl.txt", row.names=FALSE, quote = FALSE, sep = "\t")
write.table(de_fil_sig_upb, file="sig_up_il11_ctrl.txt", row.names=FALSE, quote = FALSE, sep = "\t")

# Genes you want to highlight/label
genes_interest <- c(
  "TRPV1", "TRPV2", "NTRK1", "HRH1", "VGF", "BDNF",
  "KCNH1", "KCNK9", "SCN10A", "SCN3B", "ATF3", "SOCS3"
)

# Subset DE results
de_interest <- de_filb[de_filb$gene %in% genes_interest, ]

labels <- de_interest

labels$label_colour <- ifelse(
  labels$padj < 0.05 & labels$log2FoldChange > 0, "red",
  ifelse(
    labels$padj < 0.05 & labels$log2FoldChange < 0,
    "blue",
    "black"
  )
)

# Creating volcano plot 
vol_plotb <- ggplot(de_filb, aes(x = log2FoldChange, y = mlog10p)) +
  
  # All genes
  geom_point(colour = "black", size = 2) +
  
  # Significant upregulated
  geom_point(
    data = de_fil_sig_upb,
    colour = "red",
    size = 2
  ) +
  
  # Significant downregulated
  geom_point(
    data = de_fil_sig_downb,
    colour = "blue",
    size = 2
  ) +
  
  # Highlight genes of interest
  geom_point(
    data = de_interest,
    colour = "green",
    size = 3
  ) +
  
  # Labels
  geom_label_repel(
    data = labels,
    aes(label = gene),
    colour = labels$label_colour,
    size = 8,
    fontface = "bold",
    force = 30,
    force_pull = 0.5,
    max.overlaps = Inf,
    box.padding = 1,
    point.padding = 1,
    segment.colour = "black",
    show.legend = FALSE
  ) +
  
  labs(
    title = "DEGs IL-11 vs Ctrl", x="Log2 fold change",
    x = "Log2 fold change",
    y = "-log10(FDR)"
  ) +
  
  coord_cartesian(
    xlim = c(-10, 10),
    ylim = c(0, 80),
    clip = "off"
  ) +
  
  theme(axis.title.x=element_text(size = 30),
        plot.title = element_text(size = 30, hjust = 0.5), 
        axis.title.y = element_text(size = 30),
        axis.text.x = element_text(size = 20),
        axis.text.y = element_text(size = 20),
        axis.line.x = element_line(colour = "black"),
        axis.line.y = element_line(colour = "black"),
        panel.background = element_blank(),
        panel.grid.major = element_line(colour = "lightgrey"))

emf("Vol_plot_il11_ctrl_genesinterest.emf", height=10, width=10) 
print(vol_plotb)
dev.off()
```

```
## quartz_off_screen 
##                 2
```

```
de = results(dds_wald, c("condition","IL-6+sIL-6R","Ctrl")) 

# save to disk
de_out = as.data.frame(de)

# Ensures every row had a unique name 
rownames(de_out) = ifelse(rownames(de_out) == "" | is.na(rownames(de_out)),
                          paste0("Unknown_", seq_along(rownames(de_out))),
                          rownames(de_out))

# Add gene column for row names for saving
de_out = de_out %>% rownames_to_column(var = "gene")

# merge with TPM (abundance)
de_fil = merge(de_out, abundance_out_fil, by = "gene")

#add log10 p value
de_fil$mlog10p = -log10(de_fil$padj)

#order master by padj
sorted_order = order(de_fil[,"padj"], decreasing=FALSE) 
de_filc = de_fil[sorted_order,]

write.table(de_filc, file='il6_ctrl_filtered.txt', row.names = FALSE, quote = FALSE, sep = "\t")

# subsetting data based on significance (norm & transformed counts)
de_fil_sigc = subset(de_filc, padj < 0.05)
de_fil_sig_upc = subset(de_filc, padj < 0.05 & (log2FoldChange) >0)
de_fil_sig_downc = subset(de_filc, padj < 0.05 & (log2FoldChange) < 0)
write.table(de_fil_sigc, file="sig_il6_ctrl.txt", row.names=FALSE, quote = FALSE, sep = "\t")
write.table(de_fil_sig_downc, file="sig_down_il6_ctrl.txt", row.names=FALSE, quote = FALSE, sep = "\t")
write.table(de_fil_sig_upc, file="sig_up_il6_ctrl.txt", row.names=FALSE, quote = FALSE, sep = "\t")

# Genes you want to highlight/label
genes_interest <- c(
  "TRPV1", "TRPV2", "NTRK1", "HRH1", "VGF", "BDNF",
  "KCNH1", "KCNK9", "SCN10A", "SCN3B", "ATF3", "SOCS3"
)

# Subset DE results
de_interest <- de_filc[de_filc$gene %in% genes_interest, ]

labels <- de_interest

labels$label_colour <- ifelse(
  labels$padj < 0.05 & labels$log2FoldChange > 0, "red",
  ifelse(
    labels$padj < 0.05 & labels$log2FoldChange < 0,
    "blue",
    "black"
  )
)

# Creating volcano plot 
vol_plotc <- ggplot(de_filc, aes(x = log2FoldChange, y = mlog10p)) +
  
  # All genes
  geom_point(colour = "black", size = 2) +
  
  # Significant upregulated
  geom_point(
    data = de_fil_sig_upc,
    colour = "red",
    size = 2
  ) +
  
  # Significant downregulated
  geom_point(
    data = de_fil_sig_downc,
    colour = "blue",
    size = 2
  ) +
  
  # Highlight genes of interest
  geom_point(
    data = de_interest,
    colour = "green",
    size = 3
  ) +
  
  # Labels
  geom_label_repel(
    data = labels,
    aes(label = gene),
    colour = labels$label_colour,
    size = 8,
    fontface = "bold",
    force = 30,
    force_pull = 0.5,
    max.overlaps = Inf,
    box.padding = 1,
    point.padding = 1,
    segment.colour = "black",
    show.legend = FALSE
  ) +
  
  labs(
    title = "DEGs IL-6 vs Ctrl", x="Log2 fold change",
    x = "Log2 fold change",
    y = "-log10(FDR)"
  ) +
  
  coord_cartesian(
    xlim = c(-10, 10),
    ylim = c(0, 80),
    clip = "off"
  ) +
  
  theme(axis.title.x=element_text(size = 30),
        plot.title = element_text(size = 30, hjust = 0.5), 
        axis.title.y = element_text(size = 30),
        axis.text.x = element_text(size = 20),
        axis.text.y = element_text(size = 20),
        axis.line.x = element_line(colour = "black"),
        axis.line.y = element_line(colour = "black"),
        panel.background = element_blank(),
        panel.grid.major = element_line(colour = "lightgrey"))

emf("Vol_plot_il6_ctrl_genesinterest.emf", height=10, width=10) 
print(vol_plotc)
dev.off()
```

```
## quartz_off_screen 
##                 2
```

```
vol_plota
```

```
vol_plotb
```

```
vol_plotc
```

Generating gene of interest graphs

```
gene_cyto = abundance_filtered[row.names(abundance_filtered) %in% c("TRPV1", "TRPV2", "NTRK1", 
                                                  "HRH1", "VGF", "BDNF", "KCNH1", "KCNK9", 
                                                  "SCN10A", "SCN3B", "ATF3", "SOCS3"), ]
gene_cyto = row.names(gene_cyto)
gene_cyto = abundance_filtered[gene_cyto,]

gene_cyto = data.frame(t(gene_cyto))
gene_cyto$condition = sample_sheet_cyto$condition
gene_cyto$neurone = sample_sheet_cyto$neurone

gene_cyto_m = melt(gene_cyto, id.vars = c("condition", "neurone"))


# Subset relevant conditions
sub_gene_cyto <- subset(gene_cyto_m,
                        condition %in% c("LIF", "LIF+tofa"))

# Desired order
gene_order <- c("TRPV1", "TRPV2", "NTRK1", 
                "HRH1", "VGF", "BDNF", "KCNH1", "KCNK9", 
                "SCN10A", "SCN3B", "ATF3", "SOCS3")

# Make variable a factor with levels in the desired order
sub_gene_cyto$variable <- factor(sub_gene_cyto$variable, levels = gene_order)


gene_cyto_plot = ggplot(sub_gene_cyto, aes(x = condition, y = value, group = neurone))+ 
  geom_line(colour="black", alpha=0.6)+
  geom_point(aes(shape = neurone, colour = condition), size = 4,
             position = position_jitter(width = 0.05, height = 0)) +
  scale_colour_manual(values = c(
    "LIF" = "gold",
    "LIF+tofa" = "darkorange"))+
  facet_wrap(~variable, scales = "free_y",nrow = 3)+
  labs(title="", x="", y="Abundance (TPM)")+
  #guides(colour="none")+
  theme(plot.title = element_text(size = 18, hjust = 0.5), 
        axis.title.y = element_text(size = 18),
        axis.text.x = element_text(size = 18, angle = 45, hjust = 1),
        legend.key.size = unit(1, "cm"),
        legend.text = element_text(size=18),
        legend.title = element_text(size=14),
        panel.background = element_rect(fill ="white"),  
        panel.grid = element_line(colour = "lightgrey"),
        axis.ticks = element_line(colour = "lightgrey"), 
        panel.border = element_rect(colour = "black", fill = NA),
        strip.text = element_text(size = 20),
        strip.background = element_blank())

emf("lif_liftofa_genes_plot_paper.emf", height=10, width=10) 
print(gene_cyto_plot)
dev.off()
```

```
## quartz_off_screen 
##                 2
```

```
gene_cyto_plot
```

Super exact test - determining where gene overlap is statistically
significant

```
#how to determine universe number (the number of genes that could have appeared in both analyses)
lif_genes = rownames(de_fila)
il11_genes = rownames(de_filb)
il6_genes = rownames(de_filc)


universe_no = Reduce(intersect, list(lif_genes, il6_genes, il11_genes))

length(universe_no)
```

```
## [1] 12845
```

```
sets <- list(
  A = rownames(de_fil_sig_upa),
  B = rownames(de_fil_sig_upb),
  C = rownames(de_fil_sig_upc)
)

sets$B <- sets$B[sets$B != "" & !is.na(sets$B)]

res <- supertest(sets, n = 12856)
stats_res <- data.frame(
  Combination = names(res$overlap.sizes),
  Observed = unname(res$overlap.sizes),
  Expected = unname(res$overlap.expected),
  Fold.enrichment = unname(res$overlap.sizes) / unname(res$overlap.expected),
  P.value = unname(res$P.value)
)


write.table(stats_res, file="super_exact_test_results_cyto.txt", sep = "\t", row.names = FALSE, quote = FALSE)
```
