## Supplementary material for "LIF is a pro-algesic mediator in human and mouse models of arthritis-related pain": Fibroblast CM RNA-seq R script

LIF\_paper\_fibCM


### LIF\_paper\_fibCM

###### Rosie Ross

#### 2026-08-25

```
knitr::opts_knit$set(
  root.dir = "/Users/franziskadenk/Library/CloudStorage/Dropbox/WTCA/kallisto_output/kallisto_for_paper")
    
knitr::opts_chunk$set(
  echo = TRUE,
  warning = FALSE,
  message = FALSE
)

library(DESeq2)
```

```
library(biomaRt)
library(dplyr)
library(purrr)
library(tximport)
library(grid) 
library(enrichplot)
```

```
## enrichplot v1.26.6 Learn more at https://yulab-smu.top/contribution-knowledge-mining/
## 
## Please cite:
## 
## Guangchuang Yu, Li-Gen Wang, and Qing-Yu He. ChIPseeker: an
## R/Bioconductor package for ChIP peak annotation, comparison and
## visualization. Bioinformatics. 2015, 31(14):2382-2383
```

```
library(ComplexHeatmap)
```

```
## ========================================
## ComplexHeatmap version 2.22.0
## Bioconductor page: http://bioconductor.org/packages/ComplexHeatmap/
## Github page: https://github.com/jokergoo/ComplexHeatmap
## Documentation: http://jokergoo.github.io/ComplexHeatmap-reference
## 
## If you use it in published research, please cite either one:
## - Gu, Z. Complex Heatmap Visualization. iMeta 2022.
## - Gu, Z. Complex heatmaps reveal patterns and correlations in multidimensional 
##     genomic data. Bioinformatics 2016.
## 
## 
## The new InteractiveComplexHeatmap package can directly export static 
## complex heatmaps into an interactive Shiny app with zero effort. Have a try!
## 
## This message can be suppressed by:
##   suppressPackageStartupMessages(library(ComplexHeatmap))
## ========================================
```

```
library(SuperExactTest)
```

```
## 
## Attaching package: 'SuperExactTest'
## 
## The following objects are masked from 'package:lubridate':
## 
##     intersect, union
## 
## The following objects are masked from 'package:dplyr':
## 
##     intersect, union
## 
## The following objects are masked from 'package:GenomicRanges':
## 
##     intersect, union
## 
## The following object is masked from 'package:GenomeInfoDb':
## 
##     intersect
## 
## The following objects are masked from 'package:IRanges':
## 
##     intersect, union
## 
## The following objects are masked from 'package:S4Vectors':
## 
##     intersect, union
## 
## The following objects are masked from 'package:BiocGenerics':
## 
##     intersect, union
## 
## The following objects are masked from 'package:base':
## 
##     intersect, union
```

```
theme_set(theme_gray(base_family = "Arial"))
```

Uploading and processing aligned files (abundance.tsv files from
Kallisto)

```
base_dir = "/Users/franziskadenk/Library/CloudStorage/Dropbox/WTCA/kallisto_output/kallisto_for_paper"

# List all subdirectories 
folders = list.dirs(base_dir, recursive = FALSE, full.names = TRUE)

# Filter for directories starting with "RR_" 
rr_folders = folders[basename(folders) %>% str_detect("^RR_\\d+")]

# Construct full paths to abundance.tsv files
files = file.path(rr_folders, "abundance.tsv")
names(files) = basename(rr_folders)

# Filter out any that don't exist (just in case)
files = files[file.exists(files)]

# Double-check
all(file.exists(files))  # Should be TRUE
```

```
## [1] TRUE
```

```
# Connect to Ensembl
mart <- useEnsembl(biomart = "genes", dataset = "hsapiens_gene_ensembl", host = "https://may2025.archive.ensembl.org")

# Use one abundance.tsv file to get transcript IDs
sample_file = read_tsv(files[1])
ensembl_ids = sample_file$target_id

# Remove version suffix (e.g., ENS...1.2 -> ENS...1)
ensembl_ids_clean <- sub("\\..*", "", ensembl_ids)

# for gene symbols 
tx2gene = getBM(attributes = c("ensembl_transcript_id", "external_gene_name"),
                filters = "ensembl_transcript_id",
                values = ensembl_ids_clean,
                mart = mart)
txi = tximport(files,
                type = "kallisto",
                tx2gene = tx2gene,
                ignoreTxVersion = TRUE)
# Save txi 
saveRDS(txi, file = "txi_fibCM.rds")
write.csv(txi$abundance, file = "txi_fibCM_TPM.csv")
write.csv(txi$counts, file = "txi_fibCM_counts.csv")
```

```
# read in sample sheet 
sample_sheet_fibcm = read.table("sample_sheet_2.txt", header = TRUE, row.names = 1, sep="\t")
sample_sheet_fibcm = sample_sheet_fibcm[colnames(txi$counts), ]

# sample names in sample_sheet need to match colnames in txi$counts including the order
all(colnames(txi$counts) == rownames(sample_sheet_fibcm)) # Should return TRUE
```

```
## [1] TRUE
```

```
# makes sure the group information is a factor
sample_sheet_fibcm$condition = factor(sample_sheet_fibcm$condition)
sample_sheet_fibcm$fibroblast = factor(sample_sheet_fibcm$fibroblast)

# run DESeq from Tximport (accounting for fibroblast + condition)
dds = DESeqDataSetFromTximport(txi,
                               colData = sample_sheet_fibcm,
                               design = ~ fibroblast + condition)
```

```
library(circlize)
# ============================================================
# 1. Marker genes
# ============================================================

marker_groups <- list(
  Neuronal = c(
    "TUBB3", "SST", "PRPH", "P2RX3", "POU4F1",
    "SCN9A", "TRPV1", "NTRK1"
  ),
  
  Glial = c(
    "FABP7", "SOX10", "PLP1", "MBP"
  ),
  
  Fibroblast = c(
    "PDGFRA", "DCN", "COL1A1", "LUM"
  ),
  
  Mural = c(
    "PDGFRB", "NOTCH3", "ACTA2", "TAGLN"
  )
)

marker_genes <- unlist(marker_groups, use.names = FALSE)


# ============================================================
# 2. TPM matrix
# ============================================================

tpm <- txi$abundance

# Make sure all dds samples are present in txi
stopifnot(all(colnames(dds) %in% colnames(tpm)))

# Put TPM columns in same order as dds
tpm <- tpm[, colnames(dds), drop = FALSE]


# ============================================================
# 3. Check marker genes
# ============================================================

marker_present <- marker_genes[
  marker_genes %in% rownames(tpm)
]

marker_missing <- setdiff(
  marker_genes,
  rownames(tpm)
)

cat("Missing marker genes:\n")
```

```
## Missing marker genes:
```

```
print(marker_missing)
```

```
## character(0)
```

```
# ============================================================
# 4. Subset to marker genes
# ============================================================

marker_tpm <- tpm[
  marker_present,
  ,
  drop = FALSE
]

# ============================================================
# 5. Log-transform TPM for display
# ============================================================

marker_mat <- log2(marker_tpm + 1)

# ============================================================
# 6. Define gene groups
# ============================================================

gene_group <- rep(
  names(marker_groups),
  lengths(marker_groups)
)

names(gene_group) <- unlist(
  marker_groups,
  use.names = FALSE
)

gene_group <- gene_group[
  rownames(marker_mat)
]

gene_group <- factor(
  gene_group,
  levels = c(
    "Neuronal",
    "Glial",
    "Fibroblast",
    "Mural"
  )
)


# ============================================================
# 7. Condition information from dds
# ============================================================

condition <- factor(dds$condition)

# ============================================================
# 8. Order samples by condition
# ============================================================

column_order <- order(condition)

marker_mat <- marker_mat[
  ,
  column_order,
  drop = FALSE
]

condition <- condition[
  column_order
]

# ============================================================
# 9. Define TPM legend breakpoints
# ============================================================

tpm_breaks <- c(
  0,
  1,
  10,
  100,
  1000
)

log_breaks <- log2(tpm_breaks + 1)

# ============================================================
# 10. Colour scale
# ============================================================

max_log <- max(marker_mat, na.rm = TRUE)

col_fun <- colorRamp2(
  log_breaks,
  c(
    "white",
    "#FFFFCC",
    "#FED976",
    "#FD8D3C",
    "#BD0026"
  )
)

# ============================================================
# 11. Heatmap
# ============================================================

ht <- Heatmap(
  marker_mat,
  
  name = "TPM",
  
  col = col_fun,
  
  # Legend is positioned according to log2(TPM+1),
  # but labelled in actual TPM units
  heatmap_legend_param = list(
    at = log_breaks,
    labels = tpm_breaks,
    title = "TPM"
  ),
  
  
  # ----------------------------------------------------------
  # ROWS
  # ----------------------------------------------------------
  
  row_split = gene_group,
  
  cluster_rows = FALSE,
  
  show_row_names = TRUE,
  
  row_names_gp = gpar(
    fontsize = 10
  ),
  
  row_title_gp = gpar(
    fontsize = 11,
    fontface = "bold"
  ),
  
  row_title_rot = 0,
  
  
  # ----------------------------------------------------------
  # COLUMNS
  # ----------------------------------------------------------
  
  column_split = condition,
  
  cluster_columns = FALSE,
  
  cluster_column_slices = FALSE,
  
    column_labels = as.character(dds$fibroblast[column_order]),
show_column_names = TRUE,
  
  column_names_gp = gpar(
    fontsize = 7
  ),
  
  column_names_rot = 90,
  
  column_title_gp = gpar(
    fontsize = 10,
    fontface = "bold"
  ),
  
  
  # ----------------------------------------------------------
  # FORMATTING
  # ----------------------------------------------------------
  
  border = TRUE,
  
  column_gap = unit(
    2,
    "mm"
  ),
  
  row_gap = unit(
    2,
    "mm"
  )
)


draw(ht)
```

```
pdf(
  "marker_heatmap.pdf",
  width = 7,
  height = 4
)

draw(ht)

dev.off()
```

```
## quartz_off_screen 
##                 2
```

DESeq analysis - generating a model accounting for patient fibroblast
and condition

```
#filter counts >=10 counts in >=5 samples in at least 1 condition

raw_counts = counts(dds)
by_cond = split(seq_len(ncol(dds)), colData(dds)$condition)

min_count = 10
req_n = 5

pass_by_cond = lapply(by_cond, function(idx) {
  rowSums(raw_counts[, idx, drop = FALSE] >= min_count) >= req_n
})

keep_any = Reduce(`|`, pass_by_cond)

cat("Kept", sum(keep_any), "genes out of", nrow(dds),
    "that have ≥", min_count, "counts in", req_n,
    "samples in at least one Condition\n")
```

```
## Kept 13831 genes out of 25796 that have ≥ 10 counts in 5 samples in at least one Condition
```

```
dds = dds[keep_any, ]

# save dds as RDS files
saveRDS(dds, file = "dds_fibCM.rds")
dds_wald = DESeq(dds)
saveRDS(dds_wald, file = "dds_wald_fibCM.rds")
```

Creating PCA plot

```
# creating PCA plot (uses top 500 most variable genes)
pca_data = plotPCA(vst(dds), intgroup = c("fibroblast", "condition"), returnData = TRUE)
percentVar = round(100*attr(pca_data, "percentVar"))


# Set the factor levels to define legend order
pca_data$condition = factor(pca_data$condition,
                            levels = c("Unstim_CM", "Unstim_CM+anti-LIF","IL1B_CM", "IL1B_CM+anti-LIF"))
pca_data$fibroblast = factor(pca_data$fibroblast,
                            levels = c("STX8RA", "STX14RA","STX15RA", "STX25RA","STX27RA"))


# plot PCA
pca_plot = ggplot(pca_data, aes(x = PC1, y = PC2, colour = condition, shape = fibroblast)) +
  geom_point(size = 10, stroke = 2)+ 
  guides (
    colour = guide_legend(order = 1),
    shape = guide_legend(order =2))+
  scale_colour_manual(values = c("Unstim_CM" = "turquoise4" , 
                                 "Unstim_CM+anti-LIF" = "turquoise", 
                                 "IL1B_CM" = "violet",
                                 "IL1B_CM+anti-LIF"= "purple"))+
  scale_shape_manual(values = c("STX8RA" = 16, 
                                "STX15RA" = 17, 
                                "STX25RA" = 15, 
                                "STX14RA" = 3, 
                                "STX27RA" = 18))+
  labs(title = "Fibroblast CM", 
       x = paste0("PC1: (", percentVar[1], "%)"),
       y = paste0("PC2: (", percentVar[2], "%)"))+
  theme(
    plot.title = element_text(hjust = 0.5, size = 32), 
    axis.title = element_text(size = 32),
    axis.text  = element_text(size = 16),
    panel.background = element_rect(fill = "white"),
    panel.grid = element_line(colour = "grey92"), 
    panel.border = element_rect(colour = "lightgrey", fill = NA, linewidth = 2),
    legend.title = element_text(size = 32),
    legend.text  = element_text(size = 20),
    legend.key.size = unit(1.2, "cm"))

emf("PCA_plot.emf", height=8, width=10) 
print(pca_plot)
dev.off()
```

```
## quartz_off_screen 
##                 2
```

```
pca_plot
```

Generating and saving useful files including: 1. Abundance (TPM) 2.
Raw counts 3. vst count (vst = variance stabilising transformation) 4.
Normalised counts (size factor, accounting for seq\_depth)

```
# Extracting abundance 
abundance = txi$abundance
# Ensures every row had a unique name 
rownames(abundance) = ifelse(rownames(abundance) == "" | is.na(rownames(abundance)),
                             paste0("Unknown_", seq_along(rownames(abundance))),
                             rownames(abundance))

# Re-order sample sheet and NT_counts by condition
order_index = order(sample_sheet_fibcm$condition)
sample_sheet_fibcm = sample_sheet_fibcm[order_index,]
abundance = abundance[,order_index]

# Add gene column for row names for saving 
abundance_out = as.data.frame(abundance) %>%
  rownames_to_column(var = "gene")
stopifnot(all(rownames(sample_sheet_fibcm) == colnames(abundance)))

# Save abundance as txt file
write.table(abundance_out, file="abundance_fibCM.txt", sep = "\t", row.names = FALSE, quote = FALSE)

#Extracting raw counts 
raw_counts = txi$counts
# Ensures every row had a unique name 
rownames(raw_counts) = ifelse(rownames(raw_counts) == "" | is.na(rownames(raw_counts)),
                             paste0("Unknown_", seq_along(rownames(raw_counts))),
                             rownames(raw_counts))

# Re-order raw_counts by condition
raw_counts = raw_counts[,order_index]

# Add gene column for row names for saving 
raw_counts_out = as.data.frame(raw_counts) %>%
  rownames_to_column(var = "gene")
stopifnot(all(rownames(sample_sheet_fibcm) == colnames(raw_counts)))

# Save raw counts as txt file
write.table(raw_counts_out, file="raw_counts_fibCM.txt", sep = "\t", row.names = FALSE, quote = FALSE)

# Extracting vst counts
NT_counts = vst(dds)

# Ensures every row had a unique name 
rownames(NT_counts) = ifelse(rownames(NT_counts) == "" | is.na(rownames(NT_counts)),
                              paste0("Unknown_", seq_along(rownames(NT_counts))),
                              rownames(NT_counts))
# Extract numeric matix 
NT_counts = assay(NT_counts)

# Re-order NT_counts by condition
NT_counts = NT_counts[,order_index]

# Add gene column for row names for saving 
NT_counts_out = as.data.frame(NT_counts) %>%
  rownames_to_column(var = "gene")
stopifnot(all(rownames(sample_sheet_fibcm) == colnames(NT_counts)))

# Save NT_counts as txt file
write.table(NT_counts_out, file="NT_counts_fibCM.txt", sep = "\t", row.names = FALSE, quote = FALSE)

# Extract the normalised counts
norm_counts = as.data.frame(counts(dds_wald, normalized=TRUE)) # using dds_wald as need size factor for normalisation

# Ensures every row had a unique name 
rownames(norm_counts) <- ifelse(rownames(norm_counts) == "" | is.na(rownames(norm_counts)),
                                paste0("Unknown_", seq_along(rownames(norm_counts))),
                                rownames(norm_counts))

#round the data (makes it easier to look at)
norm_counts = round(norm_counts,2)

# Re-order by condition
norm_counts = norm_counts[,order_index]

# Add gene column for row names for saving 
norm_counts_out = as.data.frame(norm_counts) %>%
  rownames_to_column(var = "gene")
stopifnot(all(rownames(sample_sheet_fibcm) == colnames(norm_counts)))

# Save norm_counts as txt file
write.table(norm_counts_out, file="norm_counts_fibCM.txt", sep = "\t", row.names = FALSE, quote = FALSE)
```

Pairwise comparison - 2 comparisons 1. IL1B vs Unstim 2. IL1B+AL vs
IL1B Generates volcano plot highlighting genes of interest

```
# 1. IL1B vs Unstim
de = results(dds_wald, c("condition","IL1B_CM","Unstim_CM")) 

# save to disk
de_out = as.data.frame(de)

# Ensures every row had a unique name 
rownames(de_out) = ifelse(rownames(de_out) == "" | is.na(rownames(de_out)),
                          paste0("Unknown_", seq_along(rownames(de_out))),
                          rownames(de_out))

# Add gene column for row names for saving
de_out = de_out %>% rownames_to_column(var = "gene")

# merge with TPM (abundance)
de_nonfil_a = merge(de_out, abundance_out, by = "gene")

#add log10 p value
de_nonfil_a$mlog10p = -log10(de_nonfil_a$padj)

#order master by padj
sorted_order = order(de_nonfil_a[,"padj"], decreasing=FALSE)    
de_nonfil_a = de_nonfil_a[sorted_order,]

write.table(de_nonfil_a, file="il1b_unstim.txt", row.names=FALSE, quote = FALSE, sep = "\t")

# subsetting data based on significance (norm & transformed counts)
de_nonfil_a_sig = subset(de_nonfil_a, padj < 0.05)
de_nonfil_a_sig_up = subset(de_nonfil_a, padj < 0.05 & (log2FoldChange) >0)
de_nonfil_a_sig_down = subset(de_nonfil_a, padj < 0.05 & (log2FoldChange) < 0)
write.table(de_nonfil_a_sig, file="sig_il1b_unstim.txt", row.names=FALSE, quote = FALSE, sep = "\t")
write.table(de_nonfil_a_sig_down, file="sig_down_il1b_unstim.txt", row.names=FALSE, quote = FALSE, sep = "\t")
write.table(de_nonfil_a_sig_up, file="sig_up_il1b_unstim.txt", row.names=FALSE, quote = FALSE, sep = "\t")


# Genes you want to highlight/label
genes_interest = c(
  "TRPV1", "TRPV2", "NTRK1", "HRH1", "VGF", "BDNF",
  "KCNH1", "KCNK9", "SCN10A", "SCN3B", "ATF3", "SOCS3"
)

# Subset DE results
de_interest = de_nonfil_a[de_nonfil_a$gene %in% genes_interest, ]

labels = de_interest

labels$label_colour = ifelse(
  labels$padj < 0.05 & labels$log2FoldChange > 0, "red",
  ifelse(
    labels$padj < 0.05 & labels$log2FoldChange < 0,
    "blue",
    "black"
  )
)

# creating volcano plot
vol_plot = ggplot(de_nonfil_a, aes(x = log2FoldChange, y = mlog10p)) +
  
  # All genes
  geom_point(colour = "black", size = 2) +
  
  # Significant upregulated
  geom_point(
    data = de_nonfil_a_sig_up,
    colour = "red",
    size = 2
  ) +
  
  # Significant downregulated
  geom_point(
    data = de_nonfil_a_sig_down,
    colour = "blue",
    size = 2
  ) +
  
  # Highlight genes of interest
  geom_point(
    data = de_interest,
    colour = "green",
    size = 3
  ) +
  
  # Labels
  geom_label_repel(
    data = labels,
    aes(label = gene),
    colour = labels$label_colour,
    size = 8,
    fontface = "bold",
    force = 30,
    force_pull = 0.5,
    max.overlaps = Inf,
    box.padding = 1,
    point.padding = 1,
    segment.colour = "black",
    show.legend = FALSE
  ) +
  
  labs(
    title = "DEGs IL-1B Fib CM vs Unstim Fib CM", x="Log2 fold change",
    x = "Log2 fold change",
    y = "-log10(FDR)"
  ) +
  
  coord_cartesian(
    xlim = c(-8, 8),
    ylim = c(0, 30),
    clip = "off"
  ) +
  
  theme(axis.title.x=element_text(size = 30),
        plot.title = element_text(size = 30, hjust = 0.5), 
        axis.title.y = element_text(size = 30),
        axis.text.x = element_text(size = 20),
        axis.text.y = element_text(size = 20),
        axis.line.x = element_line(colour = "black"),
        axis.line.y = element_line(colour = "black"),
        panel.background = element_blank(),
        panel.grid.major = element_line(colour = "lightgrey"))

emf("Vol_plot_il1b_unstim_genesinterest.emf", height=10, width=10) 
print(vol_plot)
dev.off()
```

```
## quartz_off_screen 
##                 2
```

```
vol_plot
```

```
# 2. IL1B+AL vs IL1B
de = results(dds_wald, c("condition","IL1B_CM+anti-LIF","IL1B_CM")) 

# save to disk
de_out = as.data.frame(de)

# Ensures every row had a unique name 
rownames(de_out) = ifelse(rownames(de_out) == "" | is.na(rownames(de_out)),
                          paste0("Unknown_", seq_along(rownames(de_out))),
                          rownames(de_out))

# Add gene column for row names for saving
de_out = de_out %>% rownames_to_column(var = "gene")

# merge with TPM (abundance)
de_nonfil_b = merge(de_out, abundance_out, by = "gene")

#add log10 p value
de_nonfil_b$mlog10p = -log10(de_nonfil_b$padj)

#order master by padj
sorted_order = order(de_nonfil_b[,"padj"], decreasing=FALSE)    
de_nonfil_b = de_nonfil_b[sorted_order,]

write.table(de_nonfil_b, file="il1bAL_il1b.txt", row.names=FALSE, quote = FALSE, sep = "\t")


# subsetting data based on significance (norm & transformed counts)
de_nonfil_b_sig = subset(de_nonfil_b, padj < 0.05)
de_nonfil_b_sig_up = subset(de_nonfil_b, padj < 0.05 & (log2FoldChange) >0)
de_nonfil_b_sig_down = subset(de_nonfil_b, padj < 0.05 & (log2FoldChange) < 0)
write.table(de_nonfil_b_sig, file="sig_il1bAL_il1b.txt", row.names=FALSE, quote = FALSE, sep = "\t")
write.table(de_nonfil_b_sig_down, file="sig_down_il1bAL_il1b.txt", row.names=FALSE, quote = FALSE, sep = "\t")
write.table(de_nonfil_b_sig_up, file="sig_up_il1bAL_il1b.txt", row.names=FALSE, quote = FALSE, sep = "\t")

# Genes you want to highlight/label
genes_interest = c(
  "TRPV1", "TRPV2", "NTRK1", "HRH1", "VGF", "BDNF",
  "KCNH1", "KCNK9", "SCN10A", "SCN3B", "ATF3", "SOCS3"
)

# Subset DE results
de_interest = de_nonfil_b[de_nonfil_b$gene %in% genes_interest, ]

labels = de_interest

labels$label_colour = ifelse(
  labels$padj < 0.05 & labels$log2FoldChange > 0, "red",
  ifelse(
    labels$padj < 0.05 & labels$log2FoldChange < 0,
    "blue",
    "black"
  )
)

# creating volcano plot
vol_plot = ggplot(de_nonfil_a, aes(x = log2FoldChange, y = mlog10p)) +
  
  # All genes
  geom_point(colour = "black", size = 2) +
  
  # Significant upregulated
  geom_point(
    data = de_nonfil_b_sig_up,
    colour = "red",
    size = 2
  ) +
  
  # Significant downregulated
  geom_point(
    data = de_nonfil_b_sig_down,
    colour = "blue",
    size = 2
  ) +
  
  # Highlight genes of interest
  geom_point(
    data = de_interest,
    colour = "green",
    size = 3
  ) +
  
  # Labels
  geom_label_repel(
    data = labels,
    aes(label = gene),
    colour = labels$label_colour,
    size = 8,
    fontface = "bold",
    force = 30,
    force_pull = 0.5,
    max.overlaps = Inf,
    box.padding = 1,
    point.padding = 1,
    segment.colour = "black",
    show.legend = FALSE
  ) +
  
  labs(
    title = "DEGs IL-1B Fib CM+anti-LIF vs IL-1B Fib CM", x="Log2 fold change",
    x = "Log2 fold change",
    y = "-log10(FDR)"
  ) +
  
  coord_cartesian(
    xlim = c(-8, 8),
    ylim = c(0, 30),
    clip = "off"
  ) +
  
  theme(axis.title.x=element_text(size = 30),
        plot.title = element_text(size = 30, hjust = 0.5), 
        axis.title.y = element_text(size = 30),
        axis.text.x = element_text(size = 20),
        axis.text.y = element_text(size = 20),
        axis.line.x = element_line(colour = "black"),
        axis.line.y = element_line(colour = "black"),
        panel.background = element_blank(),
        panel.grid.major = element_line(colour = "lightgrey"))

emf("Vol_plot_il1bAL_il1b_genesinterest.emf", height=10, width=10) 
print(vol_plot)
dev.off()
```

```
## quartz_off_screen 
##                 2
```

```
vol_plot
```

Heatmap of hyper-excitability and JAK/STAT genes - LIF/SF/FibCM
analysis

```
# Load cytokine and SF files needed 

abundance_cyto  = read.table("/Users/franziskadenk/Library/CloudStorage/Dropbox/WTCA/kallisto_output/kallisto_for_paper/abundance_cytokines_filtered.txt", header = TRUE, sep = "\t", row.names = 1)
sample_sheet_cyto  = read.table("/Users/franziskadenk/Library/CloudStorage/Dropbox/WTCA/kallisto_output/kallisto_for_paper/sample_sheet.txt", header = TRUE, sep = "\t")


abundance_sf = read.table("/Users/franziskadenk/Library/CloudStorage/Dropbox/WTCA/kallisto_output/kallisto_for_paper/abundance_SF_filtered.txt", header = TRUE, sep = "\t", row.names = 1)
sample_sheet_sf = read.table("/Users/franziskadenk/Library/CloudStorage/Dropbox/WTCA/kallisto_output/kallisto_for_paper/sample_sheet_3.txt", header = TRUE, sep = "\t")

# Gene list

genes <- c(
  "VGF","BDNF","HRH1", "JUNB","LIFR","CDKN1A","CISH","IL13RA1","IL6ST","SOCS3"
)

# Condition filters

keep_cyto  <- c("Ctrl","LIF")
keep_sf    <- c("SF+isotype","SF+anti-LIF")
keep_fibcm <- c("IL1B_CM","IL1B_CM+anti-LIF")

map_cyto  <- setNames(sample_sheet_cyto$condition,  sample_sheet_cyto$ID)
map_sf    <- setNames(sample_sheet_sf$condition,    sample_sheet_sf$ID)
map_fibcm <- setNames(sample_sheet_fibcm$condition, rownames(sample_sheet_fibcm))


# Extraction function (keeps replicates)

extract_data_keep_samples <- function(df, map, keep_conditions, genes) {
  df <- df[rownames(df) %in% genes, , drop = FALSE]
  valid <- intersect(names(map), colnames(df))
  selected <- valid[ map[valid] %in% keep_conditions ]
  df <- df[, selected, drop = FALSE]
  cond_vec <- unname(map[colnames(df)])
  list(mat = df, cond = cond_vec)
}

cyto_res  <- extract_data_keep_samples(abundance_cyto,  map_cyto,  keep_cyto,  genes)
sf_res    <- extract_data_keep_samples(abundance_sf,    map_sf,    keep_sf,    genes)
fibcm_res <- extract_data_keep_samples(abundance, map_fibcm, keep_fibcm, genes)


# Transform + scale EACH dataset separately


safe_scale <- function(mat) {
  t(apply(mat, 1, function(x) {
    if (sd(x) == 0) {
      rep(0, length(x))
    } else {
      (x - mean(x)) / sd(x)
    }
  }))
}

# log2 transform
cyto_log2  <- log2(cyto_res$mat + 1)
sf_log2    <- log2(sf_res$mat + 1)
fibcm_log2 <- log2(fibcm_res$mat + 1)

# z-score per gene within each dataset
cyto_scaled  <- safe_scale(cyto_log2)
sf_scaled    <- safe_scale(sf_log2)
fibcm_scaled <- safe_scale(fibcm_log2)

# Combine (already scaled matrices)

combined <- cbind(cyto_scaled, sf_scaled, fibcm_scaled)
condition_per_col <- c(
  as.character(cyto_res$cond),
  as.character(sf_res$cond),
  as.character(fibcm_res$cond)
)

# Replace IL1B_CM with FibCM
condition_per_col <- recode(condition_per_col,
                            "IL1B_CM" = "IL-1B Fib CM",
                            "IL1B_CM+anti-LIF" = "IL-1B Fib CM+anti-LIF")


desired_order <- c("Ctrl","LIF","SF+isotype","SF+anti-LIF","IL-1B Fib CM","IL-1B Fib CM+anti-LIF")

ord <- order(factor(condition_per_col, levels = desired_order), colnames(combined))
combined <- combined[, ord, drop = FALSE]
condition_per_col <- condition_per_col[ord]


# Row groups

row_group <- rep("Other", nrow(combined))
names(row_group) <- rownames(combined)

neuronal <- c(
  "VGF","BDNF","HRH1", "JUNB"
)

jakstat <- c(
  "LIFR","CDKN1A","CISH","IL13RA1","IL6ST","SOCS3"
)

row_group[names(row_group) %in% neuronal] <- "Hyperexcitability"
row_group[names(row_group) %in% jakstat]  <- "JAK/STAT"
row_group <- factor(row_group, levels = c("Hyperexcitability","JAK/STAT"))


# Column split labels with replicate counts

cond_factor <- factor(condition_per_col, levels = desired_order)
present_levels <- desired_order[desired_order %in% unique(cond_factor)]
cond_factor <- factor(condition_per_col, levels = present_levels)

cond_counts <- table(cond_factor)[present_levels]
col_split_titles <- paste0(present_levels, " (n=", cond_counts, ")")

cond_factor_labels <- factor(
  cond_factor,
  levels = present_levels,
  labels = col_split_titles
)


# Annotation

condition_colours <- c(
  "Ctrl" = "grey",
  "LIF" = "gold",
  "SF+isotype" = "darkblue",
  "SF+anti-LIF" = "lightblue",
  "IL-1B Fib CM" = "violet",
  "IL-1B Fib CM+anti-LIF" = "purple"
)

ha <- HeatmapAnnotation(
  condition = cond_factor,
  col = list(condition = condition_colours),
  show_annotation_name = FALSE,
  show_legend = FALSE   # removes condition legend
)


# Build heatmap

ht <- Heatmap(
  combined,
  name = "Expression (z-score)",
  row_split = row_group,
  column_split = cond_factor,
  cluster_rows = FALSE,
  cluster_columns = FALSE,
  show_row_names = TRUE,
  show_column_names = FALSE,
  top_annotation = ha,
  row_title_rot = 0,
  row_gap = unit(2, "mm"),
  column_gap = unit(2, "mm"),
  heatmap_legend_param = list(
    title_gp = gpar(fontsize = 16, fontfamily = "Arial"),
    labels_gp = gpar(fontsize = 16, fontfamily = "Arial"),
    grid_width = unit(1, "cm")
  ),
  row_title_gp = gpar(fontsize = 16, fontfamily = "Arial"),
  row_names_gp = gpar(fontsize = 14, fontfamily = "Arial"),
  column_title_gp = gpar(fontsize = 16, fontfamily = "Arial"),
  column_title_rot = 45,
  border = TRUE  # important: clean outer borders
)


# Draw + add black borders

emf("LIF_SF_FIBCMgenes_acrossall_NEW.emf", height = 5, width = 12)

ht_drawn <- draw(
  ht,
  merge_legends = TRUE,
  heatmap_legend_side = "right",   # expression legend right side
  annotation_legend_side = "right"
)

n_row_slices <- nlevels(row_group)
n_col_slices <- nlevels(cond_factor)

for (ri in seq_len(n_row_slices)) {
  for (ci in seq_len(n_col_slices)) {
    decorate_heatmap_body("Expression (z-score)", 
                          row_slice = ri, 
                          column_slice = ci, {
                            grid.rect(gp = gpar(col = "black", fill = NA, lwd = 2))
                          })
  }
}

dev.off()
```

```
## quartz_off_screen 
##                 2
```

```
ht_drawn
```

Fisher exact test - determining where gene overlap is statisically
significant

```
#how to determine universe number (the number of genes that could have appeared in both analyses)
#FibCM analysis
il1bup_genes = rownames(de_nonfil_a)
il1bALdown_genes = rownames(de_nonfil_b)

universe_no = intersect(il1bup_genes, il1bALdown_genes)

length(universe_no)
```

```
## [1] 13831
```

```
# values for test 
universe = 13851
listA = 152
listB = 113
observed = 55

# test calculations 
expected = (listA * listB) / universe
fold = observed / expected

pval = phyper(
  observed - 1,
  listA,
  universe - listA,
  listB,
  lower.tail = FALSE
)

print(expected)
```

```
## [1] 1.240055
```

```
print(fold)
```

```
## [1] 44.35288
```

```
print(pval)
```

```
## [1] 1.254661e-80
```

```
#LIF vs SF vs FibCM analysis 

# Open cytokine and SF files that are needed 
lif_up  = read.table("/Users/franziskadenk/Library/CloudStorage/Dropbox/WTCA/kallisto_output/kallisto_for_paper/sig_up_lif_ctrl.txt", header = TRUE, sep = "\t", row.names = 1)

SFAL_down = read.table("/Users/franziskadenk/Library/CloudStorage/Dropbox/WTCA/kallisto_output/kallisto_for_paper/sig_down_SFAL_SF.txt", header = TRUE, sep = "\t", row.names = 1)

fibcm_genes <- data.frame(
  gene = intersect(
    de_nonfil_a_sig_up$gene,
    de_nonfil_b_sig_down$gene
  )
)


sets <- list(
  A = rownames(lif_up),
  B = rownames(SFAL_down),
  C = fibcm_genes$gene
)

sets$C <- sets$C[sets$C != "" & !is.na(sets$C)]

res <- supertest(sets, n = 12206)
stats_res <- data.frame(
  Combination = names(res$overlap.sizes),
  Observed = unname(res$overlap.sizes),
  Expected = unname(res$overlap.expected),
  Fold.enrichment = unname(res$overlap.sizes) / unname(res$overlap.expected),
  P.value = unname(res$P.value)
)


write.table(stats_res, file="super_exact_test_results.txt", sep = "\t", row.names = FALSE, quote = FALSE)
```
